# Diffusion and Viscosity in Mixed Protein Solutions

**DOI:** 10.1101/2024.10.10.617612

**Authors:** Spencer Wozniak, Michael Feig

## Abstract

The viscosity and diffusion properties of crowded protein systems were investigated with molecular dynamics simulations of SH3 mixtures with different crowders, and results were compared with experimental data. The simulations accurately reproduced experimental trends across a wide range of protein concentrations, including highly crowded environments up to 300 g/L. Notably, viscosity increased with crowding but varied little between different crowder types, while diffusion rates were significantly reduced depending on protein-protein interaction strength. Analysis using the Stokes-Einstein relation indicated that the reduction in diffusion exceeded what was expected from viscosity changes alone, with the additional slow-down attributable to transient cluster formation driven by weakly attractive interactions. Contact kinetics analysis further revealed that longer-lived interactions contributed more significantly to reduced diffusion rates than short-lived interactions. This study also highlights the accuracy of current computational methodologies for capturing the dynamics of proteins in highly concentrated solutions and provides insights into the molecular mechanisms affecting protein mobility in crowded environments.

## INTRODUCTION

Biological environments consist of dense solutions of macromolecules. Inside cells, biomolecular concentrations may be as high as 400 g/L^1^, corresponding to approximately 30% of a cell’s volume being occupied by macromolecules. Despite these high concentrations, living biological samples generally retain fluid behavior, though it is well-established that molecular diffusion is generally reduced^2^ compared to dilute conditions, and this reduction in diffusion has functional consequences, as many biological processes depend on the rate of molecular interactions. However, predicting the diffusive behavior of a specific biomolecule within a dense macromolecular environment remains a considerable challenge.

The Stokes-Einstein relations link translational diffusion, Dt, and rotational diffusion, Dr, to (shear)viscosity, η, of the environment according to:

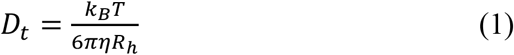

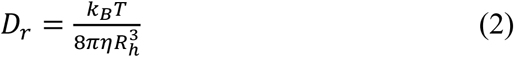

where *R*_*h*_ is the hydrodynamic radius of the molecule, *k*_*B*_ is the Boltzmann constant, and *T* is the temperature. A straightforward interpretation of **Eqs. 1** and **2** suggests that reduced diffusion at high molecular concentration arises from increased viscosity. Indeed, increased viscosity, measured independently of diffusion, has been reported in biological cells^3, 4^, cell lysates^5, 6^, and concentrated protein solutions^7^.

To explain reduced diffusion at high concentrations due to increased viscosity, generalized Stokes-Einstein relationships have been proposed instead of **Eqs. 1** and **2**^8^. The most relevant relates the long-time self-diffusion rate, DL, to the zero-shear viscosity, η:

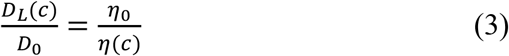

where *D*_0_ and η_0_ represent diffusion and viscosity of infinitely dilute systems, and *D*_*L*_(c) and η(c) are the corresponding values at a finite concentration *c*. **Eq. 3** effectively assumes that the hydrodynamic radius for long-time diffusion is not influenced by concentration. While this assumption holds for non-interacting hard spheres, it may not apply to biological macromolecules that are subject to attractive interactions^9-12^. In such cases, clusters may form, leading to a decrease in diffusion rates because of effectively increased hydrodynamic radius Rh, with potentially different effects on translational *vs*. rotational diffusion^13^ according to **Eqs. 1** and **2**^13-19^. Here, we distinguish clustering from oligomerization, considering clustering as less specific and more transient, though both lead to similar consequences for diffusion.

To fully understand the diffusive behavior of biological macromolecules under concentrated conditions, it is therefore necessary to separately analyze diffusion, viscosity, and the nature of transient interactions that may promote clustering. Simultaneous measurements of self-diffusion and viscosity for concentrated protein solutions under the same conditions are only available for some systems, particularly homotypic solutions of model proteins like bovine serum albumin (BSA)^13, 20-22^, lysozyme^5, 13, 22-25^, and some antibodies^26-28^ that are of technological interest. Experimental data on cluster formation for systems where both diffusion and viscosity were measured is only available for lysozyme^29^ and antibody^26^ solutions.

Recently, computer simulations have been utilized as an alternative to separately analyze translational diffusion, rotational diffusion, viscosity, and clustering for concentrated solutions of lysozyme, GB3, ubiquitin, and villin; for concentrated solutions of ubiquitin^16^; for a DNA methyltransferase and a putative lipoprotein^17^; and for a model bacterial cytoplasm with a mixture of *E. coli* proteins^30^. Additional computational studies have explored the connection between diffusion and clustering for concentrated biomolecular systems^14, 15, 31, 32^, though they did not explicitly calculate viscosities. These computational studies generally agree that significant intermolecular interactions and clustering occur, causing reduced diffusion. However, simulations are subject to force fields that may overstabilize protein-protein interactions^33, 34^, and experimental validation of both diffusion and viscosity for the systems that are being studied is needed to verify the computational interpretations of **Eq. 1** and **2** for concentrated systems.

Motivated by recent experiments^35^, we conducted an atomistic computer simulation study with the SH3 protein as a probe molecule in concentrated protein solutions consisting of either GB1, lysozyme, BSA, or ovalbumin. Experimental values for translational and rotational diffusion of SH3 were available up to protein concentrations of 200 g/L^35^. Additional data was available for SH3 diffusion in 50 g/L urea or 300 g/L sucrose^35^, providing a comparison between protein crowders and small molecule crowders. Experimental viscosity and diffusion data was also available for concentrated BSA^5, 20, 21, 25, 36, 37^, lysozyme^5, 13, 22^-^24, 38, 39^, and ovalbumin^5, 39^ solutions, as well as concentrated solutions of urea^40^ and sucrose^41^. The availability of extensive experimental data allowed us to validate computer simulations of concentrated systems, enabling interpretation of **Eqs. 1** and **2** over a wide range of concentrations, focusing not just on self-crowding but also protein mixtures and the distinction between protein and small molecule crowders.

### Systems

We studied proteins in aqueous solution at various concentrations and in the presence of different crowders. Following recent experiments^35^, we examined the T22G mutant of the N-terminal SH3 domain of drosophila drk (SH3) in the presence of different crowders up to approximately 300 g/L. In alignment with the experimental work, the crowders included the B1 domain of streptococcal immunoglobulin protein G with the T2Q mutation (GB1), hen egg white lysozyme (LYS), bovine serum albumin (BSA), chicken ovalbumin (OVA), sucrose, and urea. We chose the number of crowder molecules to achieve the desired concentrations while keeping the size of the simulated systems small enough to allow for microsecond-scale sampling with the computational resources available to us. We also simulated single copies of all proteins and crowders as dilute condition references, along with systems containing only water, with and without 150 mM NaCl salt. The systems studied are summarized in **Table 1**. Details on the simulated amino acid sequences are given in **Table S1** and snapshots of the equilibrated crowded systems are shown in **Figure 1**.

**Table 1.**
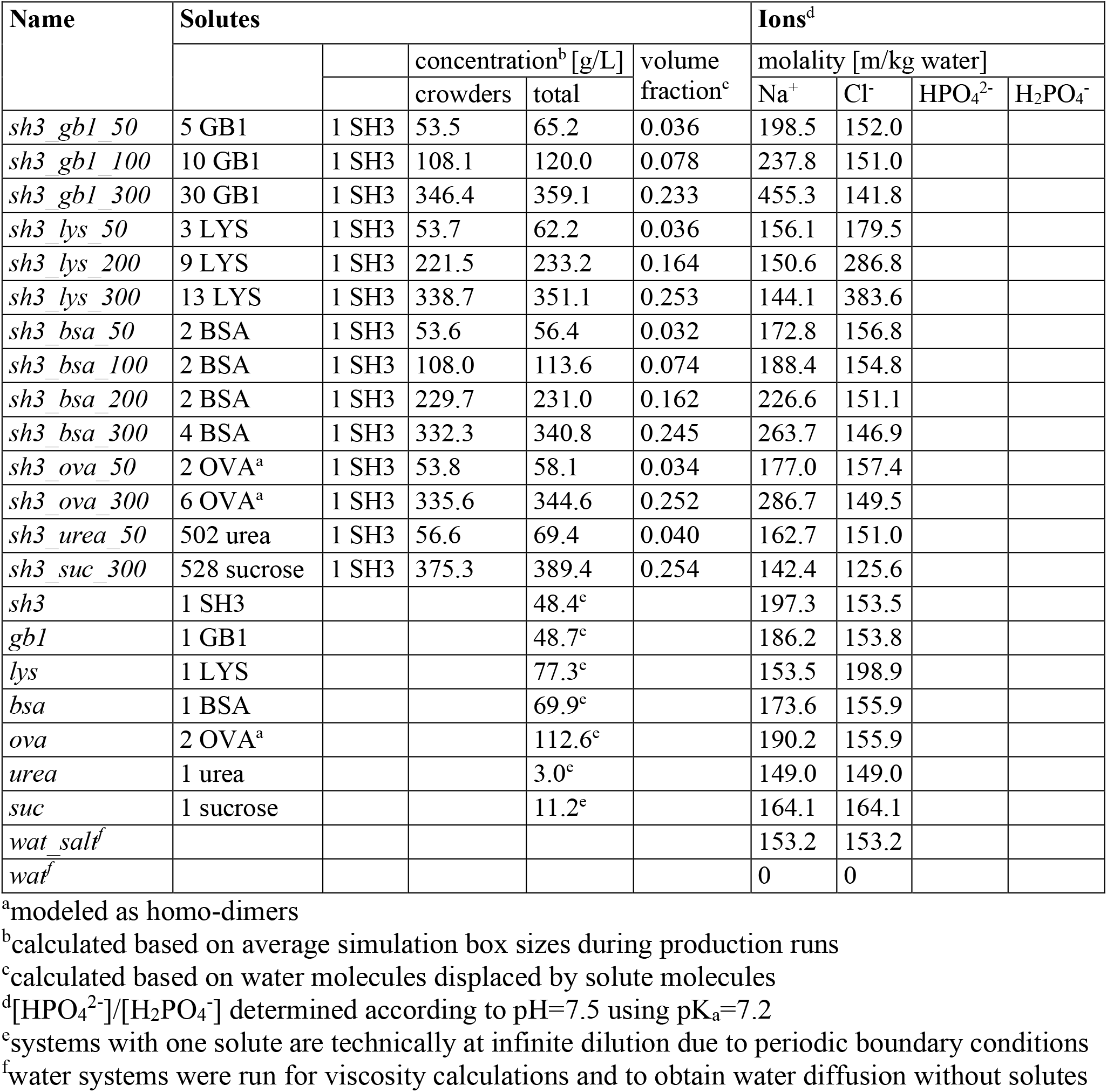
Simulated systems.

| Name | Solutes |  |  |  |  | Ions<sup>d</sup> |  |  |  |
| --- | --- | --- | --- | --- | --- | --- | --- | --- | --- |
|  |  |  | concentration<sup>b</sup> [g/L] |  | volume fraction<sup>c</sup> | molality [m/kg water] |  |  |  |
|  |  |  | crowders | total |  | Na<sup>+</sup> | Cl<sup>-</sup> | HPO<sub>4</sub><sup>2-</sup> | H<sub>2</sub>PO<sub>4</sub><sup>-</sup> |
| *sh3_gbl_50* | 5 GB1 | 1 SH3 | 53.5 | 65.2 | 0.036 | 198.5 | 152.0 |  |  |
| *sh3_gbl_100* | 10 GB1 | 1 SH3 | 108.1 | 120.0 | 0.078 | 237.8 | 151.0 |  |  |
| *sh3_gbl_300* | 30 GB1 | 1 SH3 | 346.4 | 359.1 | 0.233 | 455.3 | 141.8 |  |  |
| *sh3_lys_50* | 3 LYS | 1 SH3 | 53.7 | 62.2 | 0.036 | 156.1 | 179.5 |  |  |
| *sh3_lys_200* | 9 LYS | 1 SH3 | 221.5 | 233.2 | 0.164 | 150.6 | 286.8 |  |  |
| *sh3_lys_300* | 13 LYS | 1 SH3 | 338.7 | 351.1 | 0.253 | 144.1 | 383.6 |  |  |
| *sh3_bsa_50* | 2 BSA | 1 SH3 | 53.6 | 56.4 | 0.032 | 172.8 | 156.8 |  |  |
| *sh3_bsa_100* | 2 BSA | 1 SH3 | 108.0 | 113.6 | 0.074 | 188.4 | 154.8 |  |  |
| *sh3_bsa_200* | 2 BSA | 1 SH3 | 229.7 | 231.0 | 0.162 | 226.6 | 151.1 |  |  |
| *sh3_bsa_300* | 4 BSA | 1 SH3 | 332.3 | 340.8 | 0.245 | 263.7 | 146.9 |  |  |
| *sh3_ova_50* | 2 OVA<sup>a</sup> | 1 SH3 | 53.8 | 58.1 | 0.034 | 177.0 | 157.4 |  |  |
| *sh3_ova_300* | 6 OVA<sup>a</sup> | 1 SH3 | 335.6 | 344.6 | 0.252 | 286.7 | 149.5 |  |  |
| *sh3_urea_50* | 502 urea | 1 SH3 | 56.6 | 69.4 | 0.040 | 162.7 | 151.0 |  |  |
| *sh3_suc_300* | 528 sucrose | 1 SH3 | 375.3 | 389.4 | 0.254 | 142.4 | 125.6 |  |  |
| *sh3* | 1 SH3 |  |  | 48.4<sup>e</sup> |  | 197.3 | 153.5 |  |  |
| *gbl* | 1 GB1 |  |  | 48.7<sup>e</sup> |  | 186.2 | 153.8 |  |  |
| *lys* | 1 LYS |  |  | 77.3<sup>e</sup> |  | 153.5 | 198.9 |  |  |
| *bsa* | 1 BSA |  |  | 69.9<sup>e</sup> |  | 173.6 | 155.9 |  |  |
| *ova* | 2 OVA<sup>a</sup> |  |  | 112.6<sup>e</sup> |  | 190.2 | 155.9 |  |  |
| *urea* | 1 urea |  |  | 3.0<sup>e</sup> |  | 149.0 | 149.0 |  |  |
| *suc* | 1 sucrose |  |  | 11.2<sup>e</sup> |  | 164.1 | 164.1 |  |  |
| *wat_salt<sup>f</sup>* |  |  |  |  |  | 153.2 | 153.2 |  |  |
| *wat<sup>f</sup>* |  |  |  |  |  | 0 | 0 |  |  |

<sup>a</sup>modeled as homo-dimers<sup>b</sup>calculated based on average simulation box sizes during production runs<sup>c</sup>calculated based on water molecules displaced by solute molecules<sup>d</sup>[HPO<sub>4</sub><sup>2-</sup>]/[H<sub>2</sub>PO<sub>4</sub><sup>-</sup>] determined according to pH=7.5 using pK<sub>a</sub>=7.2<sup>e</sup>systems with one solute are technically at infinite dilution due to periodic boundary conditions<sup>f</sup>water systems were run for viscosity calculations and to obtain water diffusion without solutes

### Molecular dynamics simulations

The systems listed in **Table 1** and shown in **Figure 1** were modeled in atomistic detail and subjected to molecular dynamics (MD) simulations using explicit solvent. The simulations covered microsecond time scales, with each system simulated in triplicate to allow estimation of statistical uncertainties. **Table S2** provides further information about the system sizes, simulation times, and simulation conditions.

**Figure 1.**
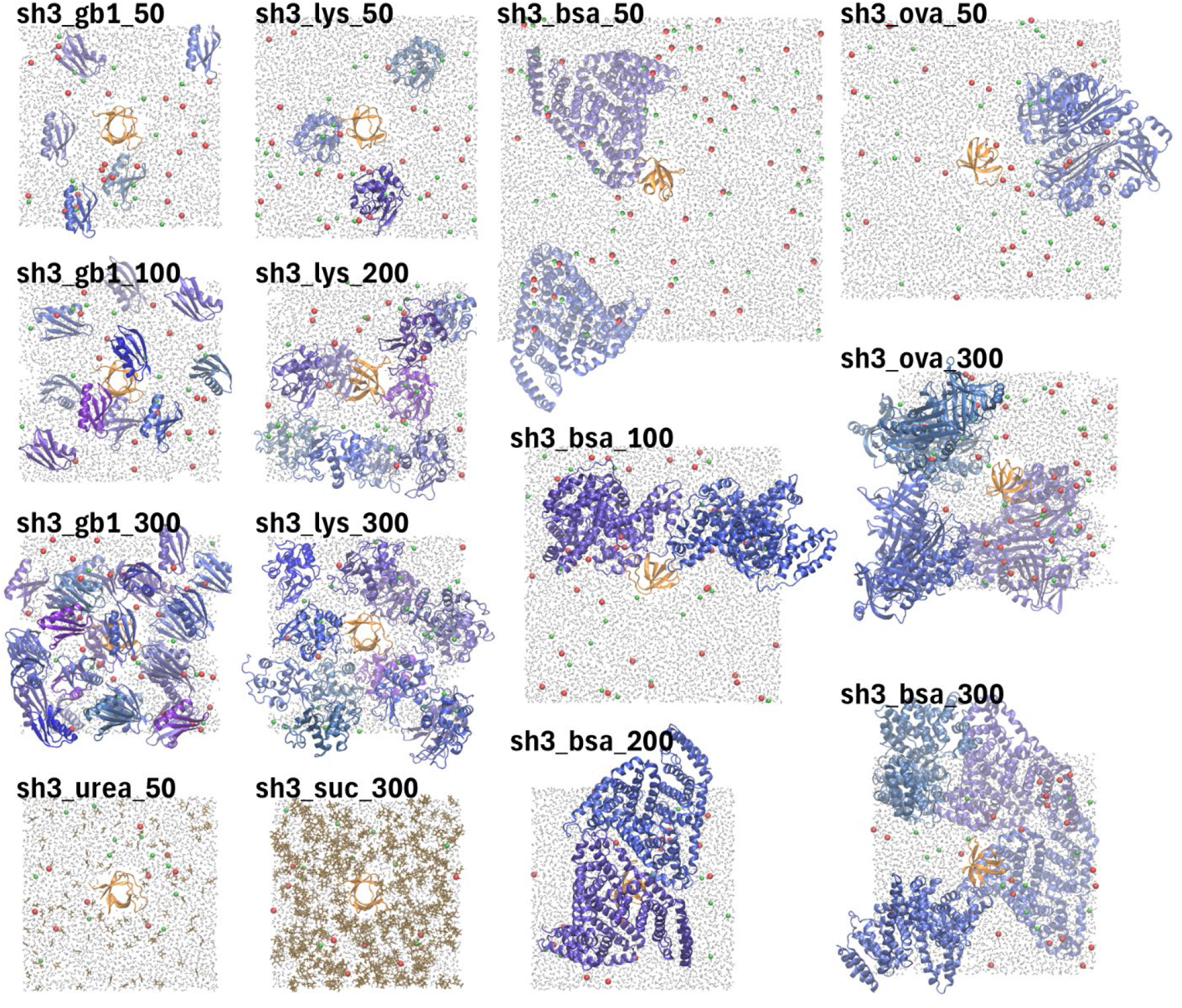
Crowded protein systems. Snapshots of the crowded systems after initial equilibration. Labels correspond to the names given in **Table 1**. SH3 is rendered in orange and protein crowders (GB1, LYS, BSA, OVA) are colored in shades of blue. Urea and sucrose are shown in brown, water molecules in grey, and ions in red (Na^+^), green (Cl^-^), or purple (phosphate). For clarity, only a slice of water and ions is shown.

Initial coordinates of proteins were obtained from experimental structures in the Protein Data Bank (see **Table S3**). Standard ionization states at pH 7 were assumed for all amino acids, and disulfide bonds were included for lysozyme, BSA, and ovalbumin according to their experimental structures. For systems with multiple solutes, the solutes were oriented and placed randomly in a simulation box, with box sizes chosen to match target concentrations (see **Table S2**). For monomeric systems, the solute molecule was centered at the origin. Following the initial placement of solutes, water molecules were added to fill the box, and ions were added last by replacing water molecules. The SH3-crowder systems were set up with the CHARMM-GUI multicomponent server^42^, while monomeric systems were set up using the MMTSB Tool Set^43^.

The solvated systems were energy-minimized and equilibrated before production simulations were started. The equilibration of SH3 in the presence of crowders was carried out by the CHARMM-GUI server^44^. The monomeric systems were first minimized using CHARMM^45^ version c46b2 over 100 steps of steepest descent, followed by 1000 steps with the gradient-based adopted-basis Newton-Raphson algorithm. Subsequent equilibration was then carried out using openMM (version 8.0)^46^ via 10 ps NVT simulations at 5, 10, and 20 K followed by 20 ps each at 50 K, 100 K, 150 K, 200 K, 250 K, and 298K. Production simulations were carried out in the NPT ensemble at 298K and 1 bar using openMM via python scripts, leveraging GPU hardware.

The CHARMM c36m force field^47^ was used to describe the proteins in all systems. Water was modeled with the CHARMM-modified TIP3P model^48, 49^ and ions were modeled according to parameters from Roux *et al*. including corrections via NBFIX^50-52^. Phosphate and urea were modeled according to cgenFF^53^ and sucrose was modeled using the CHARMM36 carbohydrate force field^54^. Electrostatic interactions were modeled via particle-mesh Ewald summation^55^, using a direct space cutoff of 12 Å with a switching function starting from 10 Å. The same cutoff and switching function were also applied for Lennard-Jones interactions. The integration time step during the production phase was 2 fs. Bonds involving hydrogen atoms were kept rigid. The temperature was maintained using the Langevin thermostat with a friction coefficient of 0.01 ps^-1^, and pressure was maintained using the Monte Carlo barostat implemented in openMM.

A second set of simulations was carried out to obtain high-frequency pressure tensor data for estimating viscosities. These simulations were restarted from 150 snapshots extracted from each of the systems studied here and continued for 1-2 ns, during which the pressure tensor was recorded every other simulation step (*i*.*e*. every 4 fs). Relatively short trajectory lengths were sufficient for estimating viscosity, provided that many independent trajectories were available for obtaining averaged pressure fluctuations^56^. For most systems, 1 ns of sampling was sufficient for converged viscosity estimates, though sampling was extended to 2 ns per snapshot for some of the larger crowded systems to improve convergence. Because pressure tensor data cannot be obtained easily from openMM, these simulations were run using NAMD^57^, version 3.0b7, using 20 cores per simulation on Intel CPUs. Simulation parameters were consistent with those used in the openMM simulations, except that the NVT ensemble was applied to fix the box size, and a stochastic velocity rescaling thermostat^58^ with a rescaling period of 1 ps was applied to maintain a temperature of 298 K.

### Convergence

The time needed to reach equilibrium was estimated based on the number of interactions between proteins and crowders, which is the most relevant factor in determining diffusive properties. Specifically, we determined the number of contacts between the SH3 protein and surrounding crowders according to a minimum heavy atom distance of less than 5 Å. From the replicate time series of these interactions, short-time averages over 400 ns intervals a_400_(t) = [*t,t*+400ns] were compared with long-time averages a_max_(t) = [*t,t*_max_] up to the maximum length of a given trajectory, which varied between 1 and 2 µs (see **Table S2**). A Z-score was then calculated according to

|a_400_ − a_max_|/(*SEM*(a_400_) + *SEM*(a_max_)) based on the standard errors of the mean (SEM) for both the short-time and long-time averages. A Z-score around 1 was used as an indication that equilibrium was reached, meaning the difference between short- and long-time averages is within one standard error. The calculated Z-scores as a function of simulation time are shown in **Fig. S1**. After 400 ns, Z-scores remained below 1 in most systems, with only a few exceptions where scores slightly exceeded 1 at certain time points. Thus, we determined that equilibrium distributions were generally reached after 400 ns, and subsequent analysis was carried out omitting the first 400 ns of each trajectory for consistency across different systems.

### Contact analysis

Contacts between proteins, urea, and sucrose were analyzed using two approaches. First, we counted the number of interacting molecules in a system based on a minimum distance of 5 Å between any heavy atom pairs between different molecules. Second, to quantify interactions for a given protein, we counted the number of heavy atoms from other molecules (*i*.*e*. crowders) within 7 Å of the Cα position of each amino acid residue of the protein.

To obtain contact survival times, a contact autocorrelation function was calculated according to **Eq. 4** as in previous work^15^:

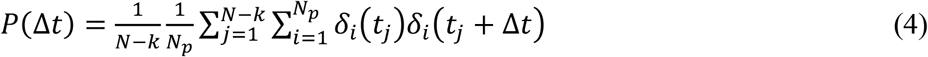

where *N* is the number of trajectory snapshots, *N*_*p*_ is the number of molecule pairs, *Δt* is the *k*-th time interval, and *δ*_*i*_ is 1 if a contact is present and 0 otherwise. A triple-exponential function according to **Eq. 5** was then fitted to the resulting correlation function from **Eq. 4** to obtain characteristic times τ_1_, τ_2_, and τ_3_:

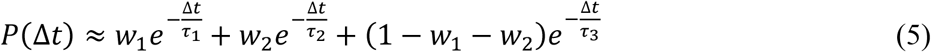

### Dissociation constants *KD* from radial distribution functions

Following previous work^16^, we calculated second virial coefficients (*B*_2_) by integrating the first peaks of protein-protein radial distribution functions (RDFs) according to:

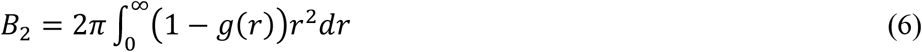

The obtained *B2* values were related to the dimensionless Baxter parameter (*τ*_*B*_)^59^ according to **Eq. 7**^60^:

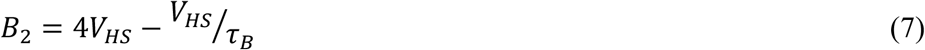

The Baxter parameter (*τ*_*B*_) captures sticky interactions between hard spheres, with a smaller value indicating more stickiness. Thus, the inverse (1/*τ*_*B*_) can be interpreted as the propensity to form clusters^16^. For homotypic interactions,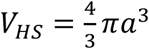, which represents the volume of a hard sphere with radius *a*. Here we will use the molecular volume of the interacting proteins. For heterotypic interactions,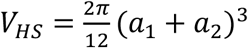 where *a1* and *a2* reflect the radii of two interacting spheres with different sizes. We use here the equivalent radii derived from the molecular volumes of interacting proteins.

The interaction dissociation constant *KD* was then obtained from *τ*_*B*_ according to **Eq. 8**^16^:

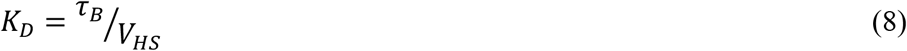

When analyzing SH3-crowder interactions, we set VHS in Eq. 8 as the volume of only SH3 in order to focus on comparing SH3 binding to different crowders instead of crowder binding to SH3.

### Viscosity calculation

Dynamic shear viscosities were calculated from equilibrium MD simulations using the Green-Kubo formalism^61-63^. In this approach, viscosity is obtained from the integral of the auto-correlation function of pressure tensor components according to:

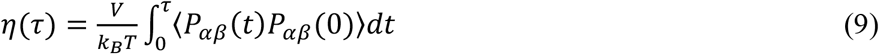

where *P*_*αβ*_ are pressure tensor components, *V* is the system volume, *k*_*B*_ is the Boltzmann constant, and *T* is the temperature (*i*.*e*. 298 K). In principle, any pressure tensor component can be used for isotropic liquids. Here, we averaged the correlation functions from all pressure tensor components, without imposing symmetry, to improve statistics.

Theoretically, one would like to obtain *η* in the limit of *τ → ∞*, but this is not practical with simulations of finite length. Instead, a general approach is to identify a plateau region where *η* remains constant as τ is increased^64^, which can be somewhat arbitrary when the running integral is noisy at longer times^63^. Here, we followed the systematic protocol by Zhang *et al*.^56^ that requires data from multiple trajectories. According to this scheme, *η*(*τ*) was evaluated for each trajectory up to a given value of *τmax*, from which averages and standard deviations across all trajectories (N=150) were obtained. The resulting standard deviation *σ*(*τ*) was fit to a power law function *σ*(*τ*) =*Aτ*^*b*^ to obtain a metric for how the uncertainty of the estimated values of *η*(*τ*) increases as *τ* is increased. A double-exponential function was then fitted to the running integral *η*(*τ*) up to *τmax* using *σ*(*τ*) as weights to be able to extrapolate to *τ → ∞*. Slightly different from the protocol by Zhang *et al*., we finally determined *τ*_max_ by evaluating *η* with increasing values of *τ*_max_ until the estimated value of *η* did not change further within statistical uncertainties from variations across trajectories.

### Translational diffusion calculation

Translational diffusion constants were estimated from linear fits to mean-square displacements (MSD) of molecular centers of mass according to the Einstein relationship:

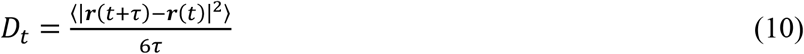

In fitting MSD vs. time, it was necessary to select an appropriate range of time to avoid anomalous behavior at shorter times, where mean-square displacement did not vary linearly with time, and poor statistics at longer times. We found that fitting MSD vs. time over the 2-10 ns interval generally satisfied these criteria.

Initial values of *Dt,PBC* estimated via **Eq. 10** from our simulations with periodic boundary conditions were then corrected for finite-size periodic artifacts^65^ according to:

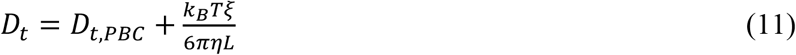

with

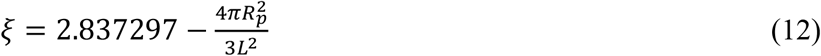

where *L* is the simulation box size, *R*_*p*_ is the size of the protein, η is the viscosity, *k*_*B*_ is the Boltzmann constant, and *T* is the temperature (298 K). For crowded systems, we used the viscosity value obtained via simulation for that system. For dilute systems with only one solute molecule, we used the calculated viscosity for TIP3P water with 0.15 m NaCl salt (0.347 cP) instead of the calculated viscosities of the systems themselves, as the systems were effectively at infinite dilution from the perspective of diffusion.

To compare with experiment, the finite-size corrected diffusion estimates obtained via **Eq. 11** were further corrected for the underestimated viscosity with the TIP3P water model to reach our final estimation of *D*_*t*_ according to:

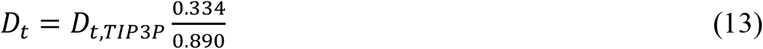

### Rotational diffusion calculation

Rotational diffusion constants were determined using two approaches: from rotational autocorrelation functions^66^ and from the time-dependent covariance matrix of the quaternions describing rotational motion^67^.

In the first approach, following the protocol by Wong *et al*.^66^, trajectories of 1,000 unity vectors with random directions were merged with trajectories of centered molecules. The merged trajectory frames were then rotated onto a reference frame for the molecule using Cα atoms, which also rotates the random vectors. From the rotated random vectors, a rotational auto-correlation function *≪ p*_*2*_(*cosθ*(*t*)*≫* was calculated and fitted to a double-exponential:

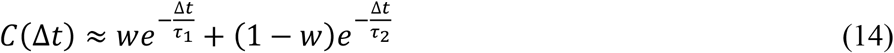

We chose the interval 0-20 ns for the fitting to avoid poor statistics at longer times.

This approach assumes an isotropic diffusion tensor but resolves slow and fast time scales. An overall relaxation time was determined according to:

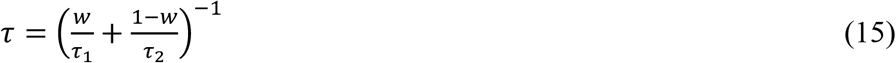

from which the rotational diffusion constant was obtained as:

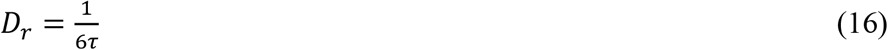

In the second approach, using a tool implemented by Linke *et al*.^67^, a fully anisotropic rotational diffusion tensor was obtained by fitting to the time-dependent quaternion covariance matrix from the evolution of Cα atoms for a given protein. From 20 fits obtained via simulated annealing, the fit with the best agreement to the time evolution of the covariance matrix up to 20 ns was selected.

This approach captured the anisotropy of the diffusion tensor and its time evolution throughout the entire time interval that was being considered (up to 20 ns). An overall relaxation time was then obtained from the three diagonal elements of the diffusion tensor according to^67^:

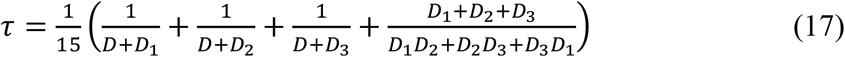

where *D*=(*D*_*1*_+*D*_*2*_+*D*_*3*_)/3 and *D*_*r*_ is then obtained via **Eq. 16**.

Initial estimates of *D*_*r*_ with periodic boundary conditions (*Dr,PBC*) as obtained via **Eq. 16** were corrected according to^68^:

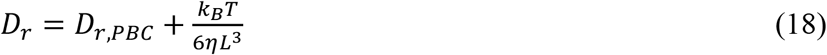

using the calculated viscosities and box length L. As in **Eq. 11** we used the value for salt water for systems with only one solute.

Finally, the finite-size corrected rotational diffusion estimates according to **Eq. 18** were again corrected for the underestimated viscosity with the TIP3P water model according to:

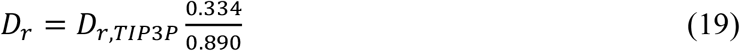

## RESULTS AND DISCUSSION

### Protein stability

The proteins remained generally stable, as indicated by low average root mean-square deviations (RMSD) from their experimental structures (**Table S4**). Average RMSD values were around 2 Å for SH3, 1 Å for GB1, 2.5 Å for lysozyme, and 3 Å for the larger multi-domain proteins BSA and ovalbumin. Individual RMSD time series (**Figs. S2-S7**) indicate that some proteins occasionally exhibited larger deviations, which is expected with dynamic ensembles followed over microsecond time scales. Crowding had no significant impact on GB1 and minimal impact on SH3, aside from a slight, statistically significant destabilization in the presence of urea, consistent with experiments^35^. Lysozyme RMSD values were higher in the crowded systems compared to the monomer, indicating the possibility of crowding-induced destabilization, also consistent with experiments^69^. On the other hand, for BSA and ovalbumin, crowding may have stabilized the native structures slightly, as RMSD values were lower compared to the single-copy simulations.

### Protein interactions

Frequent protein interactions between SH3 and the crowders, as well as among the crowders themselves, were observed (**Table 2** and **Figs. 2-4**). As expected, interactions increased with crowder concentration. For example, SH3 interacted, on average, with about one GB1 crowder at 50 g/L, but with four GB1 crowders at 300 g/L. There was also significant variation depending on the type of crowder. For example, at 50 g/L crowder concentrations, SH3 interacted, on average, with about two lysozyme crowders, compared to just one GB1 or ovalbumin crowder, and only 0.3 BSA crowders. This suggests an interaction preference of SH3 in the order: lysozyme > GB1 = ovalbumin > BSA. Analysis of crowder heavy atom contacts per SH3 residue confirmed the same order of interaction preference. Notably, this difference was most pronounced at 50 g/L but diminished at higher concentrations, at which nearly all crowders interacted with SH3 residues to a similar extent (**Fig. 2**, solid lines). This trend likely resulted from the unavoidable minimum level of interaction at higher concentrations, in addition to intrinsic interaction preferences.

**Table 2.** Crowder contacts.

| System | Interacting crowders<sup>a</sup> |  |  |  | Contacts per residue<sup>b</sup> |  |
| --- | --- | --- | --- | --- | --- | --- |
|  | SH3-crowder<br>(direct) | SH3-crowder<br>(in cluster) | Crowder-crowder | N<sub>crowder</sub> | SH3-crowder | Crowder-crowder |
| *sh3_gbl_50* | 1.07 (0.07) | 1.83 (0.23) | 0.82 (0.08) | 5 | 0.69 (0.05) | 1.49 (0.35) |
| *sh3_gbl_100* | 1.74 (0.10) | 5.99 (0.52) | 1.71 (0.04) | 10 | 0.98 (0.25) | 7.77 (0.61) |
| *sh3_gbl_300* | 3.96 (0.38) | 29.87 (0.11) | 4.09 (0.13) | 30 | 2.17 (0.32) | 38.8 (0.73) |
| *sh3_lys_50* | 2.15 (0.19) | 2.73 (0.06) | 1.32 (0.21) | 3 | 2.04 (0.18) | 1.08 (0.08) |
| *sh3_lys_200* | 3.60 (0.72) | 9.00 (0.00) | 3.37 (0.19) | 9 | 2.83 (0.33) | 8.05 (0.24) |
| *sh3_lys_300* | 4.48 (0.88) | 13.00 (0.00) | 4.16 (0.22) | 13 | 3.55 (0.76) | 12.5 (0.15) |
| *sh3_bsa_50* | 0.26 (0.07) | 0.52 (0.14) | 1.00 (0.00) | 2 | 0.14 (0.07) | 0.19 (0.05) |
| *sh3_bsa_100* | 0.69 (0.15) | 1.13 (0.23) | 0.99 (0.01) | 2 | 0.57 (0.17) | 0.20 (0.03) |
| *sh3_bsa_200* | 1.39 (0.19) | 1.99 (0.00) | 1.00 (0.00) | 2 | 1.47 (0.05) | 0.33 (0.06) |
| *sh3_bsa_300* | 2.07 (0.44) | 4.00 (0.00) | 2.33 (0.20) | 4 | 1.61 (0.19) | 0.86 (0.02) |
| *sh3_ova_50* | 0.82 (0.12) | 0.82 (0.12) | -<sup>c</sup> | 1 | 0.59 (0.16) | -<sup>c</sup> |
| *sh3_ova_300* | 1.56 (0.24) | 2.99 (0.01) | 1.99 (0.01) | 3 | 2.29 (0.41) | 0.46 (0.03) |
| *sh3_urea_50* | 15.0 (0.1) |  |  |  | 2.07 (0.02) |  |
| *sh3_suc_300* | 21.7 (0.4) |  |  |  | 6.40 (0.31) |  |

Averages omitting the first 400 ns of each trajectory; standard errors given in parentheses were estimated from variations between replicate simulations
<sup>a</sup>number of crowder molecules in contact (<5 Å heavy-atom distance)
<sup>b</sup>average number of crowder atoms in contact with Cα per residue (<7 Å heavy-atom distance)
<sup>c</sup>only one homo-dimer present in system

**Figure 2.**
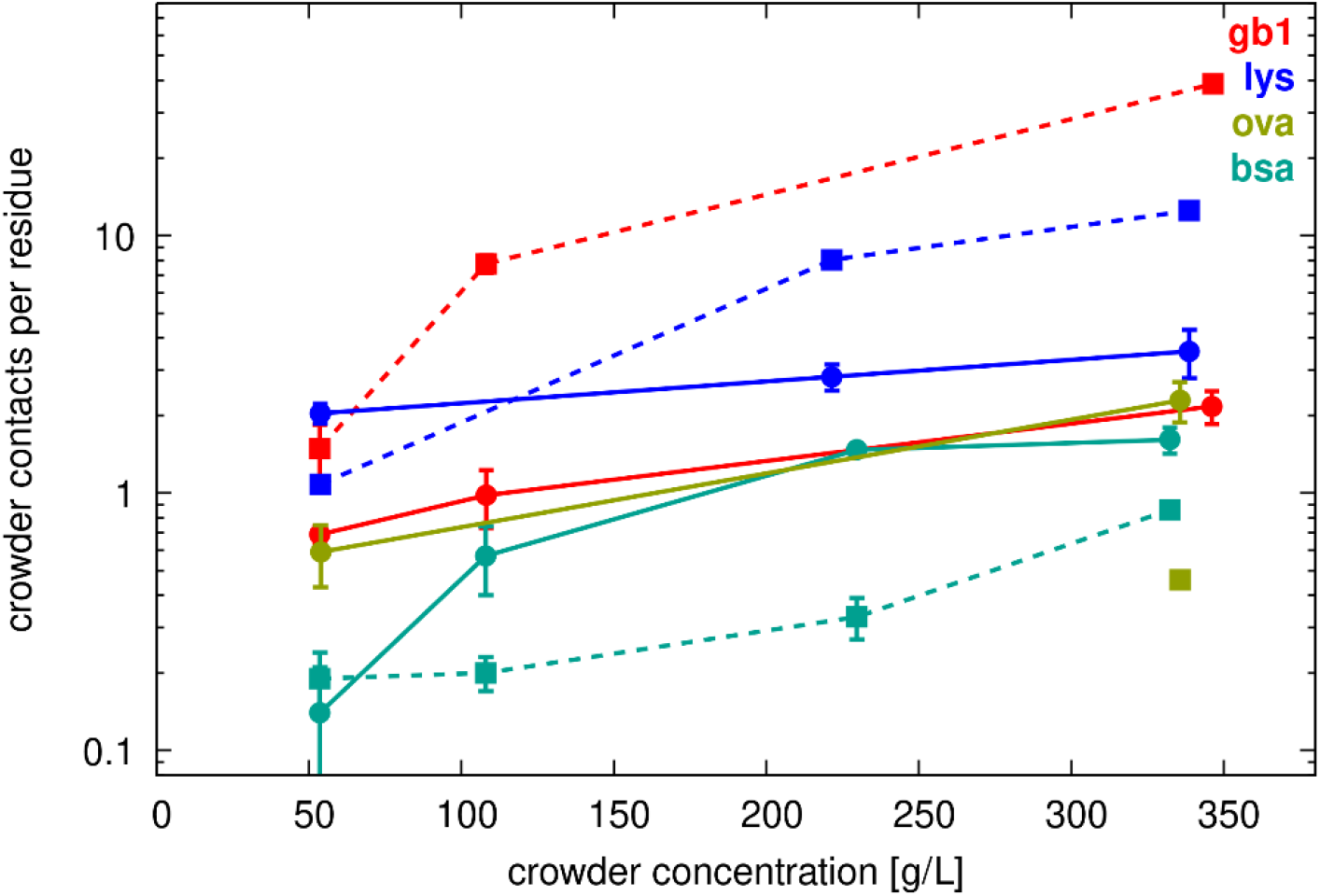
Crowder contacts vs. concentration. Average number of crowder heavy atoms within 7 Å of SH3 Cα atoms (filled circles, solid lines) or other crowder Cα atoms (filled squares, dashed lines) as a function of crowder concentration. Error bars indicate standard errors of the mean across replicate simulations.

Crowder-crowder interactions were similarly extensive, increasing with concentration and varying by crowder type. (**Table 2** and **Fig. 2**). At a given concentration, GB1 had the most extensive self-interactions, followed by lysozyme, while BSA and ovalbumin had relatively few crowder contacts per residue. However, BSA and ovalbumin crowders remained in contact with each other most of the time, as the number of interaction partners given in **Table 2** matched or closely approached the total number of possible interaction partners. For example, in the *sh3_ova_300* systems with three ovalbumin dimers, each ovalbumin dimer was on average in contact with 1.99 (out of possible two) other ovalbumin dimers. This may have resulted from the large molecular sizes of BSA and ovalbumin, which reduced packing efficiency compared to smaller molecules like GB1 and lysozyme, making close interactions involving many atoms less likely.

We compared the number of contacts per residue further (**Fig. 2**), focusing on the competition between crowder self-interactions and SH3-crowder interactions. In the SH3-lysozyme system at 50 g/L, SH3 residues interacted more extensively with lysozyme than lysozyme interacted with itself, despite the higher concentration of lysozyme (three molecules vs. one molecule SH3, **Table**

**1**). However, at higher lysozyme concentrations, SH3-lysozyme interactions were surpassed by lysozyme-lysozyme interactions, indicating that crowder self-interactions became dominant. Moreover, SH3-GB1 interactions were always less extensive than GB1 self-interactions, while SH3-BSA interactions were comparable to BSA-BSA interactions at 50 g/L, though SH3-BSA interactions increased at higher crowder concentrations (**Fig. 2**). This could be because the differing molecular sizes of SH3 and BSA facilitated more extensive interactions between SH3 and BSA compared to BSA self-interactions.

SH3-crowder interactions varied by residue according to solvent-accessible surface area (SASA), with more exposed residues (*i*.*e*. residues with greater SASA) generally involved in more crowder contacts (**Fig. 3**), and some additional variation based on crowder type. For example, ovalbumin interacted more strongly with SH3 near residue 20, whereas GB1 interacted more strongly near residue 30. However, all crowder types interacted with the most exposed residues to some extent, suggesting that SH3-crowder interactions were largely non-specific, albeit with some site preferences. The small-molecule co-solutes urea and sucrose interacted more uniformly with SH3 across all exposed residues, effectively coating the SH3 surface (**Fig. 3**).

**Figure 3.**
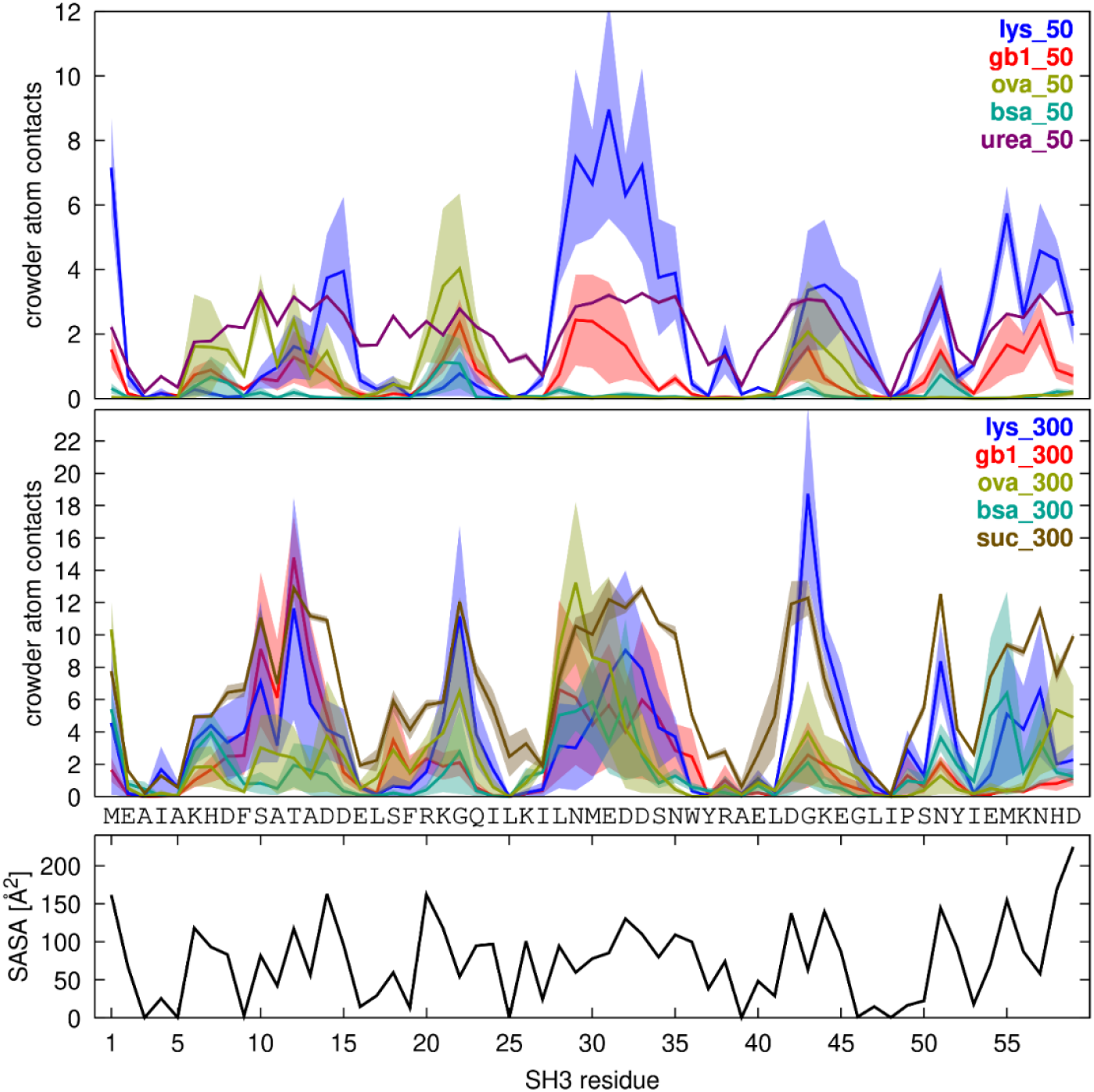
SH3-Crowder contacts. Average number of crowder heavy atoms within 7 Å of SH3 Cα atoms plotted against SH3 residue for crowded systems at 50 g/L (top) and 300 g/L (bottom). Shaded regions indicate standard errors across replicate simulations. For reference, the solvent-accessible surface area (SASA) per residue is shown at the bottom.

Beyond direct contacts, the crowder proteins also formed larger clusters, consistent with previous work^14-16, 31^. Due to the limited number of molecules in our simulations, reliable cluster size distributions could not be determined. Instead, we analyzed the average number of crowders involved in a cluster with SH3 (**Table 2**). When this number exceeded the number of directly interacting crowders, it indicated cluster formation beyond direct contacts. At lower concentrations, we observed significant clustering with lysozyme and GB1, but less with BSA. For example, at 50 g/L, SH3 typically interacted with GB1 clusters involving about two GB1 molecules, only one of which was in direct contact with SH3. At 100 g/L, SH3 interacted with about two GB1 molecules directly, but clusters involved six GB1 molecules on average. At the highest concentrations (>=200 g/L), the number of crowders in the cluster interacting with SH3 was equal to the number of crowders in all the systems. This implies that either the finite-size clusters contain more molecules than our simulations included or that concentrations exceeded solubility limits or other phase thresholds.

Timelines of SH3-crowder contacts (**Fig. 4)** show extensive but highly transient interactions at lower crowder concentrations, with more persistent and specific interactions at the highest concentrations. Crowder contact survival kinetics were quantified by fitting autocorrelations of contact survival (**Figs. S8** and **S9**) to a triple-exponential function (**Eq. 5**), revealing three characteristic time scales with corresponding weights (**Tables S5** and **S6**). We found long contacts persisting for at least 50 ns, intermediate contacts lasting for a few nanoseconds, and short contacts lasting for tens-to-hundreds of picoseconds. SH3-crowder interactions included both long- and short-lived contacts across all systems, indicating their transient nature even up to the highest concentrations. SH3-GB1 interactions showed shorter contact durations (58-113 ns for the longest contacts) compared to SH3-lysozyme interactions (333-541 ns for the longest contacts). SH3-BSA interactions were short-lived at 50 g/L but persisted (>500 ns) at higher concentrations, and SH3-ovalbumin interactions also lasted longer (**Table S5**). Of note, the longer contact times reported here are uncertain due to the small number of crowders and limited simulation times. Contacts between SH3 and the small molecule crowders urea and sucrose were mostly short-lived, but we found a long-time component, with a characteristic time of 76 ns, contributing to the contact survival decay for urea that may indicate specific binding at certain sites.

**Figure 4.**
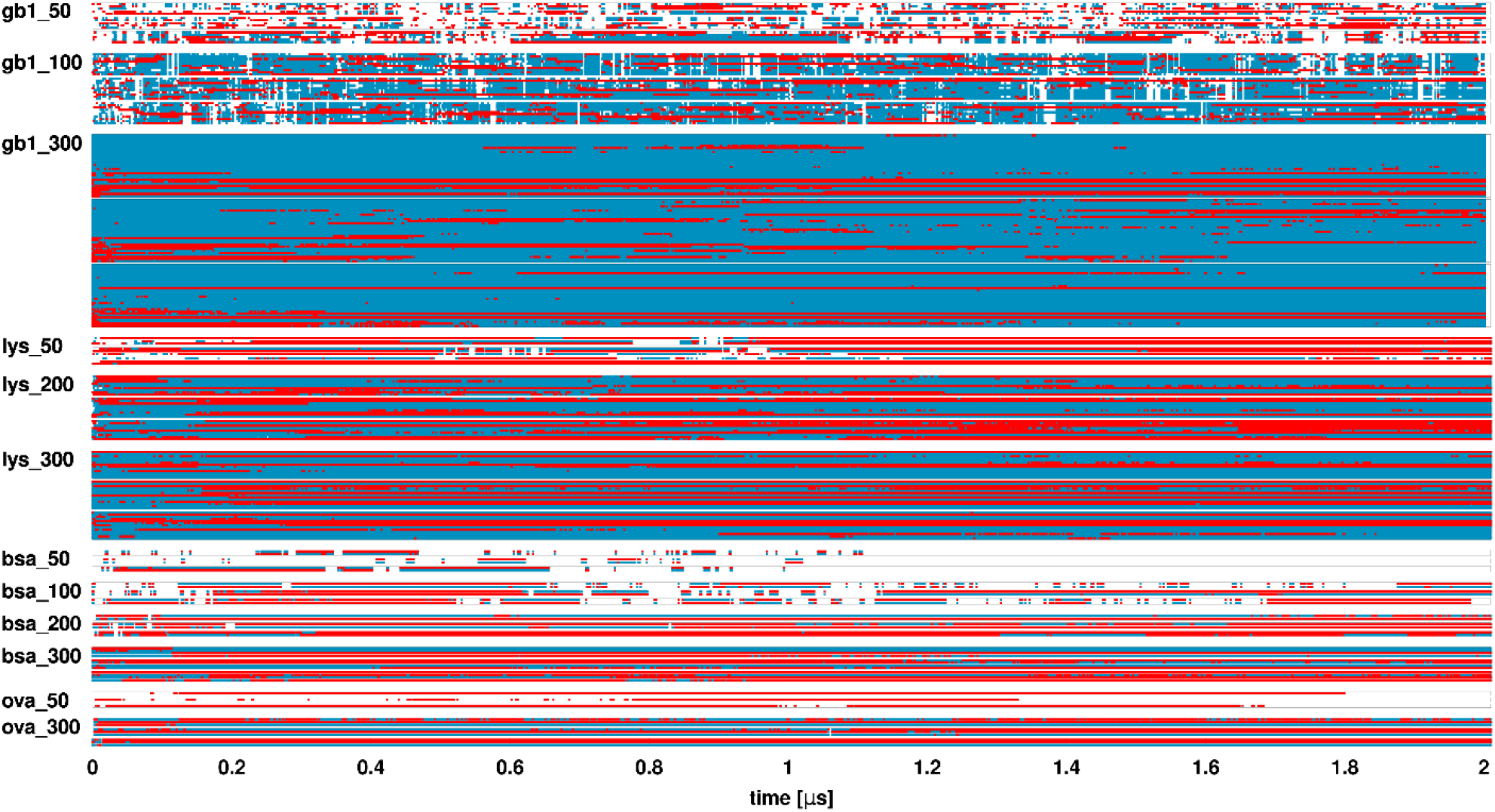
SH3-crowder contact timeline. Contacts between SH3 and crowder proteins as a function of simulation time. Each line corresponds to one crowder molecule. Results for simulation replicates are grouped in boxes. Direct contacts are shown in red, while indirect contacts (crowders in a cluster with SH3 but not in direct contact) are shown in blue.

Crowder contact kinetics were similar for GB1 and lysozyme self-interactions up to the highest concentrations, with GB1 contacts persisting for less time (64-92 ns) than lysozyme contacts (204-320 ns). This seems to suggest coupling of the SH3 and crowder dynamics in the GB1 and lysozyme systems studied here. On the other hand, BSA and ovalbumin self-interactions mostly persisted throughout a given simulation (**Table S6**), as interactions between the large molecules were stabilized across the periodic boundaries of the simulated systems.

To connect with colloid theory, we analyzed radial distribution functions (RDFs) of SH3-crowder and crowder-crowder interactions based on the centers of mass of the interacting molecules. The resulting RDFs at the lowest concentrations (50 g/L) are shown in **Fig. S9**. The RDFs are relatively noisy due to the low number of molecules and limited simulation time. Thus, we did not calculate crowder-crowder RDFs for *sh3_bsa_50* and *sh3_ova_50*. According to the RDFs, there are more interactions between SH3 and lysozyme, compared to GB1. The RDFs for SH3-BSA and SH3-ovalbumin interactions extended over a wider range, reflecting the non-spherical shapes of BSA and ovalbumin, with SH3-BSA interactions appearing much weaker than SH3-ovalbumin interactions. The order of SH3-crowder interactions generally aligns with the contact analysis results. However, crowder-crowder interactions appeared stronger for lysozyme than for GB1, different from the conclusion based on the contact analysis, in which we found more residue contacts between GB1 than lysozyme. This discrepancy suggests that contact analysis may not perfectly reflect interaction strength.

We obtained *KD* values according to **Eqs. 6-8**, and the resulting values shown in **Table 3** are generally in the same range as those reported in previous simulation work^16^. The calculated *KD* for lysozyme-lysozyme interactions (∼ 6 mM) is similar to an experimental value (∼3 mM)^70^. The estimated *KD* values for SH3-crowder interactions again suggest a decreasing interaction strength from lysozyme to GB1 and BSA, with strong SH3-ovalbumin interactions. Estimated *KD* values suggest stronger self-interactions for lysozyme compared to GB1, and the order of the *KD* values aligns with the contact survival times, with lower *KD* values corresponding to longer contact survival. The estimated *KD* values are similar for SH3-crowder and crowder self-interactions for the GB1 and lysozyme systems, again suggesting that interactions are coupled in the mixed systems, while the *KD* value for GB1 without SH3 was about double that of GB1 in the presence of SH3.

**Table 3.** Interaction analysis based on colloid theory.

| Interaction | System | $a_1$<br>[Å] | $a_2$<br>[Å] | $V_{HS}$<br>[Å<sup>3</sup>] | $B_2$<br>[Å<sup>3</sup>] | $r_{max}$<br>[Å]<sup>a</sup> | $\tau_B$ | $K_D$<br>[mM]<sup>c</sup> |
| --- | --- | --- | --- | --- | --- | --- | --- | --- |
| SH3-GB1 | sh3_gbl_50 | 12.07 | 11.70 | 7027 | -18403 | 40 | 0.151 | 34.1 |
| SH3-lysozyme | sh3_lys_50 | 12.05 | 15.40 | 10837 | -199715 | 55 | 0.045 | 10.1 |
| SH3-BSA | sh3_bsa_50 | 12.05 | 25.58 | 27920 | 102037 | 70 | 2.895 | 655.3 |
| SH3-ovalbumin | sh3_ova_50 | 12.05 | 28.11 | 33943 | -653129 | 60 | 0.043 | 9.74 |
| GB1-GB1 | sh3_gbl_50 | 11.70 | - | 6700 | -17024 | 45 | 0.153 | 37.9 |
| lysozyme-lysozyme | sh3_lys_50 | 15.40 | - | 15303 | -203652 | 55 | 0.058 | 6.3 |

<sup>a</sup>upper integration limit when obtaining $B_2$
<sup>b</sup>obtained by dividing $K_D$ in [Å<sup>-3</sup>] by $N_A/L$

### Viscosity

Viscosities were calculated from pressure tensor fluctuations in many short MD simulations via the Green-Kubo formalism according to **Eq. 9**. Stable viscosity estimates with moderate uncertainties were reached with *τ*_max_ = 100 ps (**Fig. S10**). For most systems, 1 ns simulations were sufficient to reach convergence, but for the larger dilute crowded systems (*sh3_gb1_50, sh3_bsa_50, sh3_ova_50*, and *sh3_lys_50*) and the densest systems (*sh3_gb1_300, sh3_bsa_300, sh3_ova_300*, and *sh3_lys_300*), it was beneficial to extend the simulations to 2 ns.

The viscosity values calculated with *τ*_max_ =100 ps are provided in **Table S7** for the crowded systems, single solute systems, and water with/without salt. Of note, viscosity is an intensive property that is not affected by periodic box size^65^. Therefore, viscosities calculated with only one solute reflect the nominal concentration of one molecule in the respective box sizes rather than the infinite dilution condition from the perspective of diffusion when there is only one solute in a periodic box.

Using the CHARMM-modified TIP3P water model^49^, we estimated the viscosity of pure water to be 0.334 ± 0.006 cP, which is within the uncertainty of an early estimate of 0.35 ± 0.02 cP^71^ and only slightly larger than a more recent estimate of 0.322 ± 0.005^72^. Viscosities determined with the standard TIP3P model where hydrogen atoms do not have Lennard-Jones radii are similar^72-74^, but it is well-known that the calculated values significantly underestimate the experimental value of 0.89 cP for pure water at 298 K^75^. Thus, with the expectation that viscosities calculated for the crowded systems differ from experimental values in a similar manner, we focus our subsequent discussion on relative viscosities.

We estimated the viscosity of 0.15 m NaCl solution to be 0.347 ± 0.003 cP, or 1.038 relative to pure water based on our estimate of 0.334 cP. Considering the uncertainties in the calculated viscosities, the calculated relative viscosity is expected to fall between 1.012 and 1.067, which is consistent with the relative viscosity of 1.013 measured experimentally for 0,15 m NaCl at 298K^41^.

Most crowded systems described here contain SH3 as a probe molecule, which generally adds less than 10% to the overall macromolecular concentration, so we assume the presence of SH3 does not significantly affect the following results. Experimental shear viscosities were available for concentrated BSA^5, 20, 21, 25, 36, 37, 76, 77^, lysozyme^5^, and ovalbumin^5^ solutions (**Fig. 5**), but not for GB1 to our knowledge. The data from different experiments is largely in agreement but there are some deviations when salt conditions and/or pH change. For example, the viscosities for lysozyme reported by Roos *et al*.^*13*^ without buffer or salt in D2O (at pD 3.8) are significantly higher than other values obtained with buffer at pH near 5. To compare with our simulations, we focus here on conditions closest to pH 7 and 0.15 m NaCl.

**Figure 5.**
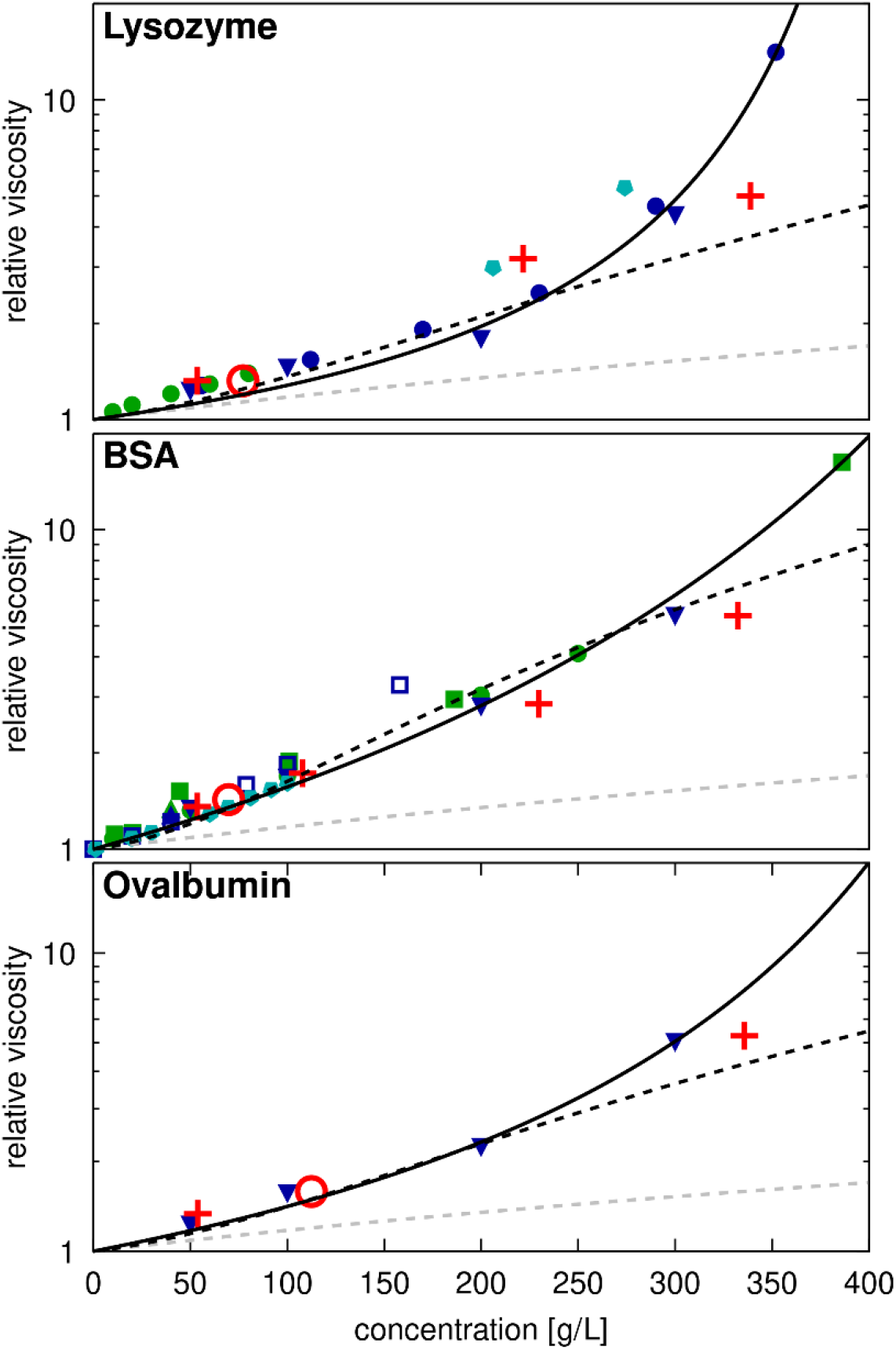
Relative viscosities for protein solutions from experiments and simulations. Calculated relative viscosities as a function of crowder concentrations in systems of lysozyme, BSA, or ovalbumin. For lysozyme, data was taken from Fredericks *et al*.^91^ (green filled circles, pH=7, 0.15 M salt), Wang *et al*.^5^ (blue filled downward triangle, pH=5.4), Riest *et al*.^38^ (blue filled circles, pH=5.4), and Roos *et al*.^*13*^ (cyan filled pentamers, no buffer, pD=3.8, not included in fit). For BSA, experimental data was taken from Castellanos *et al*.^36^ (green filled square, pH=7.4), Sharma *et al*.^37^ (green filled circle, pH=7.4), Yadav *et al*.^21^ (blue and green filled upward triangles, pH=6 and 7.4), Wang *et al*.^5^ (blue filled downward triangle, pH=5.4), Zdovc *et al*.^20^ (blue open square, pH=4.3), and Heinen *et al*.^*25*^ (cyan filled pentamer). Ovalbumin data was taken from Wang *et al*.^5^ (blue filled downward triangle, pH=5.4). Experimental data points were extracted from the literature by digitizing figures when exact values were not given, and relative viscosities were determined with 0.89 cP as the reference value for pure water. Simulation results are shown for systems with SH3 (red ‘+’) and single-copy systems (red open circles) relative to the viscosity of salt water (**Table S7**). Data points are compared with the Einstein function η_*r*_ *=* 1.0 + *2*.5*ϕ* (dashed grey line), and fits to **Eq. 20** (dashed black line) or **Eq. 21** (solid black line). The fitting parameters *b, S*, and *K* are given in **Table S8**.

The well-known Einstein equation for hard-spheres η_*r*_ *=* 1.0 + *2*.5*ϕ* (with the macromolecular volume fraction *ϕ*)^78^ does not fit the data well. A better fit is obtained with a higher order expansion^60^:

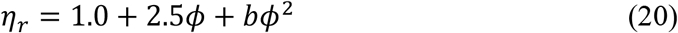

where the parameter *b* has been interpreted to capture attractive interactions between colloidal particles^60, 79^. To convert between concentrations in g/L and volume fraction *ϕ* we use c[g/L] = 1430*ϕ* from fitting concentrations *vs*. volume fraction given in **Table 1** for the systems studied here. We note that the coefficient of 1430 g/L is very similar to experimental protein densities for proteins with larger molecular weights^80^.

Using **Eq. 20**, we fit experimental viscosity data up to concentrations of around 250 g/L, resulting in values of *b*=93 for BSA, *b*=38 for lysozyme, and *b*=48 for ovalbumin. The value for lysozyme is similar to the value found from previous simulations^16^. As *b* reflects interaction strength according to colloid theory^79^, the lower value for lysozyme suggests fewer interactions compared to BSA or ovalbumin, at least up to 250 g/L. This contrasts with the literature, which suggests BSA is non-clustering^81^, while lysozyme clustering and oligomerization are well-documented in experiments^18, 29, 82-85^, suggesting that there should be more interactions between lysozyme than BSA. However, the interpretation of the experimental data is complicated as lysozyme oligomerization is favored at higher ionic strength and high pH (above 7)^81, 84, 85^, while the lysozyme viscosity measurements at higher concentrations were carried out at low pH (3.8-5.4) with little or no added salt^5, 13, 38^, where lysozymes are more likely monomeric^81, 84, 85^.

Nevertheless, the experimental data across the entire range of concentrations is described best by the semi-empirical expression originally proposed by Mooney^86^:

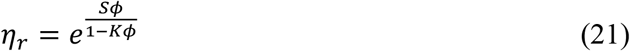

Here, the fitting coefficient *S* in **Eq. 21** is interpreted as an intrinsic viscosity, reflecting the shape and flexibility of the molecule^87^. For hard spheres, S=2.5^86^ as in the Einstein function. When fitted to experimental data, *S* is 2.8 for lysozyme, a mostly spherical protein, whereas larger values of 4.2 and 5.7 are found for ovalbumin and BSA, both of which are less spherical and may be more dynamic internally due to the presence of multiple domains. The second parameter *K* is understood as a self-crowding factor^88^ that captures packing effects. This factor indirectly reflects molecular interactions by describing if and how clusters are formed that would affect both the shape and packing at higher densities, as analyzed in detail for antibody solutions^28^. We found similar *K*values for BSA and ovalbumin, whereas a larger value was found for lysozyme, indicating more efficient packing. It follows that lysozyme exhibits lower viscosities than BSA or ovalbumin at moderate concentrations, but that its viscosity increases more rapidly at higher concentrations.

Our simulation results generally align with experimental data (**Fig. 5**), especially when considering data near pH 7. However, our simulations may slightly overestimate viscosities at lower concentrations, where SH3 occupies a larger fraction of the protein volume, while underestimating viscosities at the highest concentrations. The TIP3P water model may also contribute to underestimations at higher concentrations, consistent with previous studies on concentrated carbohydrate solutions using the same model, which found that viscosity estimations increasingly deviated from uniform scaling beyond volume fractions of 0.20 (approximately 300 g/L)^64^.

We also compared our results with experimental data for sucrose and urea solutions. Experimental data for sucrose-NaCl-water solutions at selected concentrations can be interpolated to other conditions^41^. For the *suc* system with one sucrose in a box of water (0.15 m NaCl and 0.03 m sucrose) the experimental relative viscosity is estimated at 1.043, compared to a calculated value of 1.126 (**Table S7**). For the crowded SH3-sucrose system (*sh3_suc_300*), the calculated relative viscosity was 3.539, similar to the experimental estimate of 3.79 for solution with 0.15 m NaCl and 1.48 m sucrose (matching salt and sucrose concentrations in the *sh3_suc_300* system) (**Table 1**). For the concentrated SH3-urea solution (*sh3_urea_50*, with 0.15 m NaCl and 0.99 m urea), the calculated relative viscosity was 1.26 (**Table S7**), somewhat higher than the experimental value of 1.07 (using 0.89 cP as the reference for pure water) under similar conditions (0.32 m NaCl and 0.96 m urea) at 298K^40^. In this case the calculated value may be higher because of the presence of SH3 as the protein adds about 20% to the solute concentration.

The calculated viscosities for all systems are compared in **Fig. 6**. Surprisingly, we find that the viscosity values for all protein systems, including the systems with only a single protein copy, essentially fall onto a single line. Only the viscosities for the 50 g/L urea and 300 g/L sucrose solutions are significantly lower than for protein solutions at the same concentration (**Fig. 6**). The viscosities for the protein systems can be fit with either **Eq. 20** or **Eq. 21** (**Table S8**), but the fit is slightly better with Mooney’s expression (**Eq. 21**). This is surprising given the significant differences in interactions between GB1, lysozyme, BSA, and ovalbumin crowders in the simulations. To again compare via colloid theory, the Baxter stickiness parameter *τ*_*B*_ introduced above can be related to the second-order viscosity fitting coefficient *b* according to^16, 79^: 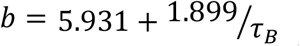. Using this expression with the value of *b*=63.4 for the fit of **Eq. 20** against the relative viscosities from simulations (**Fig. 6**), we obtain *τ*_*B*_=0.033. This value is similar to the value estimated for lysozyme-lysozyme interactions from RDFs (**Table 3**), but it is about an order of magnitude smaller than the RDF-derived *τ*_*B*_ value for GB1-GB1 interactions. Based on our analysis, it appears that interaction strength, described in terms of contacts per residue or stickiness via *τ*_*B*_, does not predict viscosity well. A lack of correspondence between protein-protein interaction strength and viscosity has been reported for antibody solutions^27^, and it may indicate that colloid hard-sphere models are not a good framework for understanding the viscosity in dense protein solutions. One reason could be that water-protein interactions, which do not depend strongly on the protein type^89, 90^, are a major determinant for explaining the increase in viscosity of concentrated protein solutions. This will be explored further below. Another reason could be that the presence of SH3 creates enough perturbation to result in more uniform molecular associations than what would be found with homotypic solutions. Finally, Mooney’s viscosity model suggests that the macroscopic viscosity of highly concentrated solutions may be influenced by larger-scale phenomena such as gelation or cluster formation involving many proteins with varied shapes and packing, which may not be fully captured in our simulations due to the limited number of proteins.

**Figure 6.**
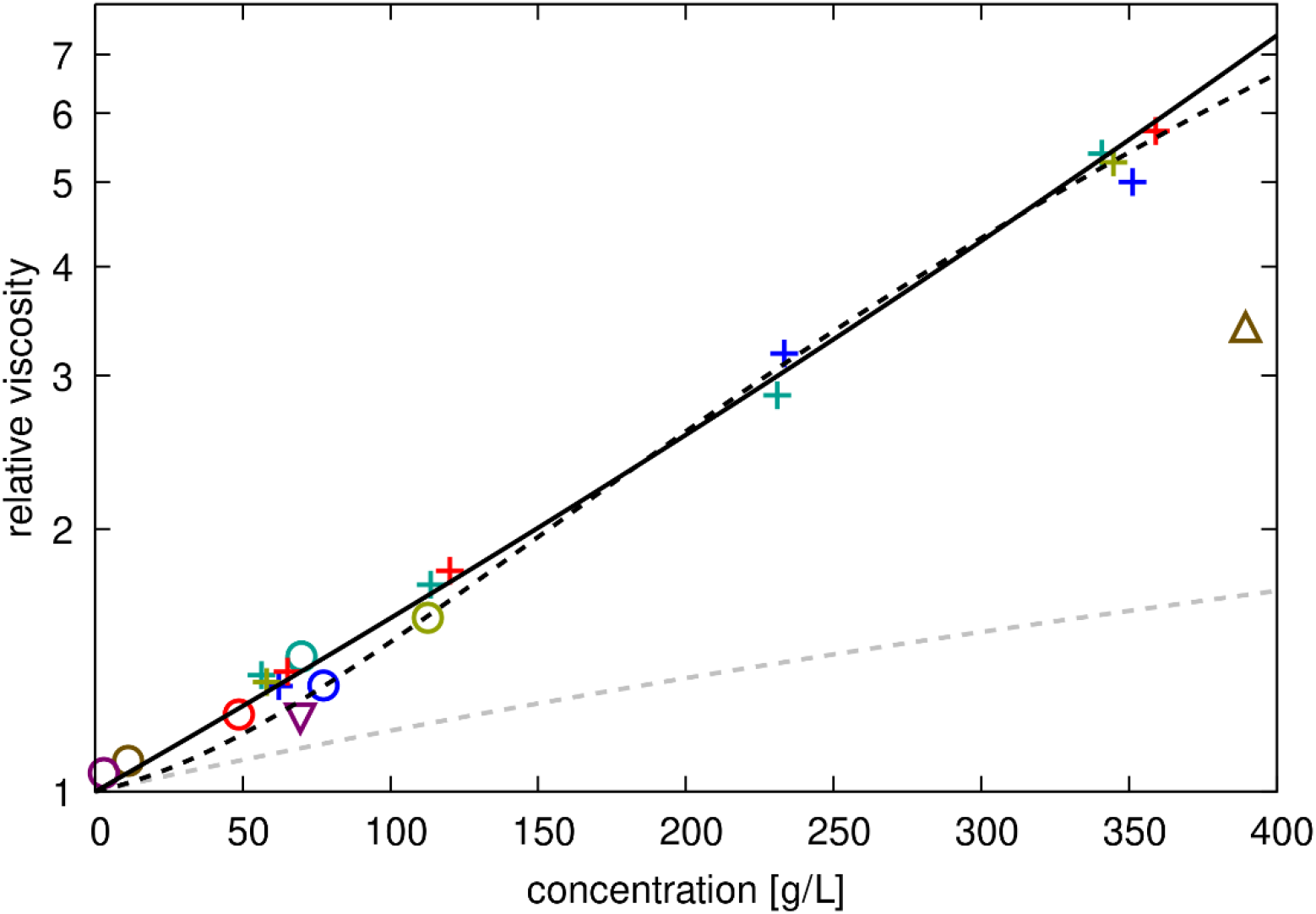
Relative viscosities for protein solutions from simulations. Calculated relative viscosities (with respect to salt water) as a function of crowder concentrations in crowded protein systems (‘+’), crowded urea (open downward triangle), crowded sucrose (open upward triangle) and single copy system (open circles). Colors distinguish between GB1 (red), lysozyme (blue), BSA (green), ovalbumin (tan), urea (purple), and sucrose (brown) crowders. Data points are compared with the Einstein function η_*r*_ *=* 1.0 + 2.5*ϕ* (dashed grey line), a fit to the experimental data to **Eq. 20** (dashed black line) or **Eq. 21** (solid black line). The fitting parameters *b, S*, and *K* are given in **Table S8**.

### Diffusion

Translational diffusion coefficients for SH3 and protein crowders were determined from mean-square displacement (MSD) curves (**Figs. S11-S12**) according to **Eqs. 10-13**. The MSD curves were generally linear in the range of 2-10 ns, and we used this interval to estimate long-term diffusion constants, *Dt*. Rotational diffusion coefficients were obtained from rotational correlation functions (**Figs. S13-S14**) following the protocol by Wong *et al*.^*66*^ (**Eqs. 14-16**) and, in addition, from fitting asymmetric diffusion tensors to the time-dependent quaternion covariance matrix following Linke *et al*.^67^. To allow direct comparisons between estimated and experimental values, our estimates of *Dt* and *Dr* were corrected for periodic box artefacts, and the PBC-corrected values were multiplied by 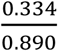 to account for the underestimated viscosity with the TIP3P water model.

Uncorrected and corrected values of *Dt* extracted from the simulations are reported in **Table S9**. Rotational diffusion coefficients *Dr* are reported in **Tables S10-S12**. Fits to the autocorrelation functions (Wong *et al*.^*66*^) and the time-depending quaternion covariance (Linke *et al*.^67^) gave different values, but differences were generally within the estimated statistical uncertainties (**Table S12**). As the results from fitting the rotational autocorrelations gave somewhat better agreement with experiment, we used those values in the subsequent analysis.

Diffusion coefficients are plotted as a function of concentration in **Figs. 7** and **8**. As expected, diffusion rates decrease significantly with concentration. There is generally good agreement with estimates from HYDROPRO^92^ and experimental data, both for SH3 diffusion in the presence of different crowders, and for crowder self-diffusion, especially when experimental data at or near pH 7 is used for comparison since diffusion depends on pH^23^. This is especially apparent for lysozyme, where experimental data for translational diffusion collected at around pH 7^22^ and rotational diffusion collected at pH 9^82^ are in excellent agreement with the simulation results (**Fig. 8**), whereas experimental data at lower pH deviates significantly^13^. Where discrepancies occurred between our simulations and experiments, like with SH3 in the presence of GB1 crowders, simulations tended to underestimate experimental values, meaning diffusion in the simulations was somewhat slower than in the experiments.

**Figure 7.**
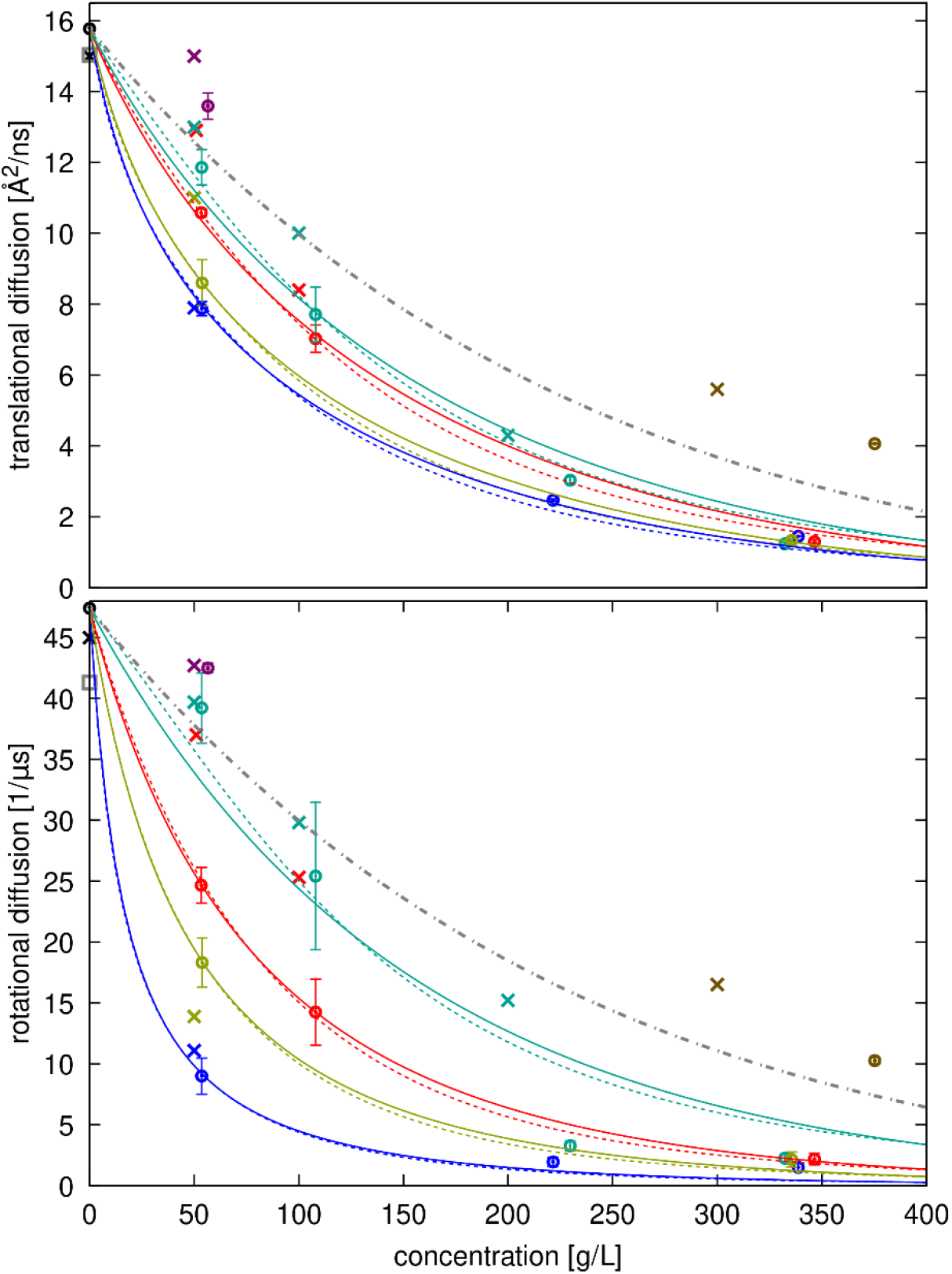
Diffusion of SH3 with different crowders as a function of concentration. Translational (top) and rotational (bottom) diffusion coefficients estimated from simulations after correction for finite-size artefacts and lower viscosity with TIP3P (open circles with error bars) are compared with experimental data^35^ (‘x’) and predictions from HYDROPRO^92^ (grey open square). Colors indicate the type of crowder: black: dilute, red: GB1, blue: lysozyme, green: BSA, tan: ovalbumin, purple: urea, brown: sucrose. The grey dot-dash line shows diffusion decreasing proportionally using **Eq. 21** with the values from **Table S8**. Colored lines show fits to **Eq. 22** for translational diffusion and **Eq. 23** for rotational diffusion with the fitted values of ζ for different crowders given in **Table S12** using either the colloid model for relative viscosity (dashed lines) or Mooney’s expression (solid lines).

**Figure 8.**
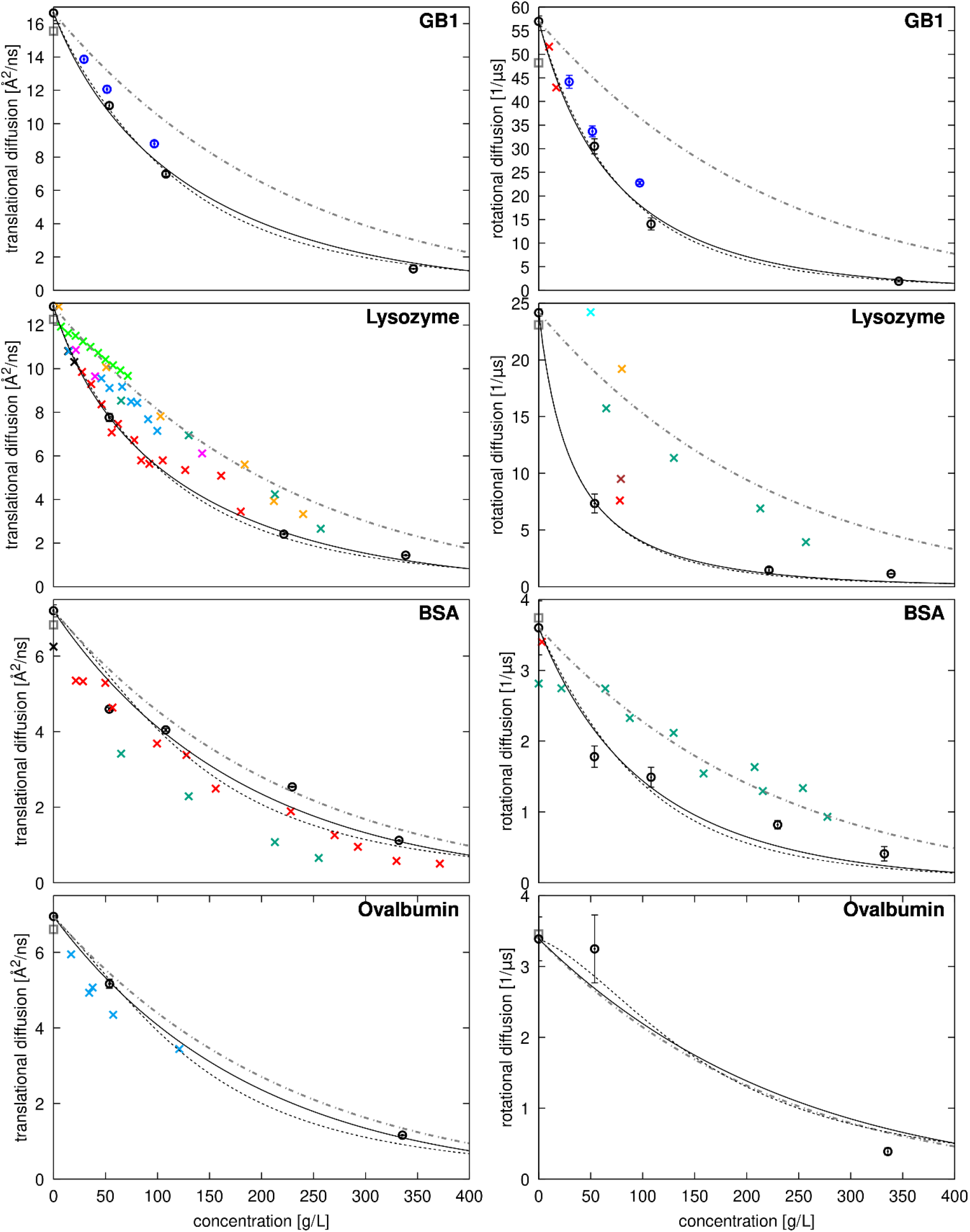
Diffusion of crowders as a function of concentration. Translational (left) and rotational (right) diffusion coefficients estimated from simulations after correction for finite-size artefacts and lower viscosity with TIP3P (open circles with error bars) are compared with experimental data (‘x’) and predictions from HYDROPRO^92^ (grey open square). For GB1, experimental data is unavailable, and simulation results are shown in black. For lysozyme, experimental data for translational diffusion is shown at dilute conditions^94, 95^ (black) and as a function of concentration from Roos *et al*.^13^ at pD=3.8 (green), from Coffman *et al*.^39^ at pH=4.5 (cyan), from Price *et al*.^23^ at pH=4.6 (light green), from Liu *et al*.^24^ at pD=7 to 4.5 from lower to higher concentrations (orange), from Nesmelova *et al*.^*22*^ at pH=7.4-7.8 (red) and from Price *et al*.^23^ at pH=8 (magenta). Experimental data for rotational diffusion is shown from Roos *et al*.^13^at pD=3.8 (green), from Gottschalk *et al*.^82^ at pH=4 without salt (orange), at pH=4 with 0.3 M NaCl (brown), and at pH=9 without salt (red), and from Buck *et al*.^96^ at pH=3.7 without salt. For BSA, experimental data for translational diffusion is shown at dilute conditions^97, 98^ (black) and as a function of concentration from Roos *et al*.^13^ at pD=7.0 without salt (dark green), and from Nesmelova et al.^22^ at pH=4.8-5.2 (red). Experimental data for rotational diffusion is shown at from Roos *et al*.^13^ at pD=7.0 (dark green) and from Wang *et al*.^99^ at pH=8 (red). For ovalbumin, experimental data for translational diffusion is shown for fluorine-labeled ovalbumin as a function of concentration at pH=7.8-8.2 from Coffman *et al*.^39^ (cyan). The dot-dash grey lines show diffusion decreasing proportionally to increased viscosity using **Eq. 21** with the values given in **Table S8**. Lines show fits to **Eq. 22** (translational diffusion) or **Eq. 23** (rotational diffusion) with the fitted values of ζ for different crowders given in **Table S12** using either the colloid model for estimating relative viscosities (dashed black lines) or Mooney’s expression (solid black lines).

Diffusion in the presence of protein crowders generally decreases more rapidly with increasing concentration than expected from the average increase in viscosity (dashed lines in **Figs. 7** and **8** based on **Eq. 20** fitted to viscosity across all simulations). This indicates that factors beyond increased viscosity contribute to the slow-down in diffusion. The reduction in SH3 diffusion varies depending on the crowder and is consistent with experimental observations, with SH3 diffusion decreasing in the order of BSA, GB1, ovalbumin, and lysozyme at the same concentrations (**Fig. 7**). Crowder diffusion is also retarded differently depending on the crowder, with GB1 and lysozyme diffusion retarded more than self-diffusion of BSA and ovalbumin.

The degree of retardation correlates with increasing strength of interactions between SH3 and the crowders (**Fig. 2** and **Table 3**). SH3 interacts relatively weakly with BSA, resulting in less diffusion retardation, while it forms stronger interactions with lysozyme, where diffusion is reduced much more significantly. Crowder diffusion is also slowed down significantly for the strongly interacting crowders lysozyme and GB1, with lysozyme experiencing a greater reduction than GB1, consistent with the stronger interactions between lysozyme molecules as estimated from RDFs (**Table 3**).

We also compared relative viscosities with relative translational and rotational diffusion coefficients using the values calculated from each of the simulations (**Fig. 9**). The generalized Stokes-Einstein relation (**Eq. 3**) applies to systems where points lie on the identity line, as seen for SH3 diffusion in urea and sucrose solutions and, in some cases, for SH3 or crowder diffusion in BSA and ovalbumin solutions. However, in most cases, **Eq. 3** does not apply, as diffusion is reduced more than expected from the increased viscosity. Our simulations and the experiments^35^ both find that SH3 diffusion is significantly slower in the presence of lysozyme and ovalbumin, whereas there is only little deviation from generalized Stokes-Einstein behavior for SH3 diffusion in the presence of BSA at lower concentrations. At higher concentrations (200 g/L), experimental diffusion rates are less reduced than predicted by deviations in viscosity, whereas our simulations do show a significant slow-down at that concentration. SH3 diffusion in urea and sucrose solutions follows Stokes-Einstein behavior in both experiments and our simulations. For GB1, a direct comparison was not possible due to a lack of experimental viscosity data for concentrated GB1 solutions.

**Figure 9.**
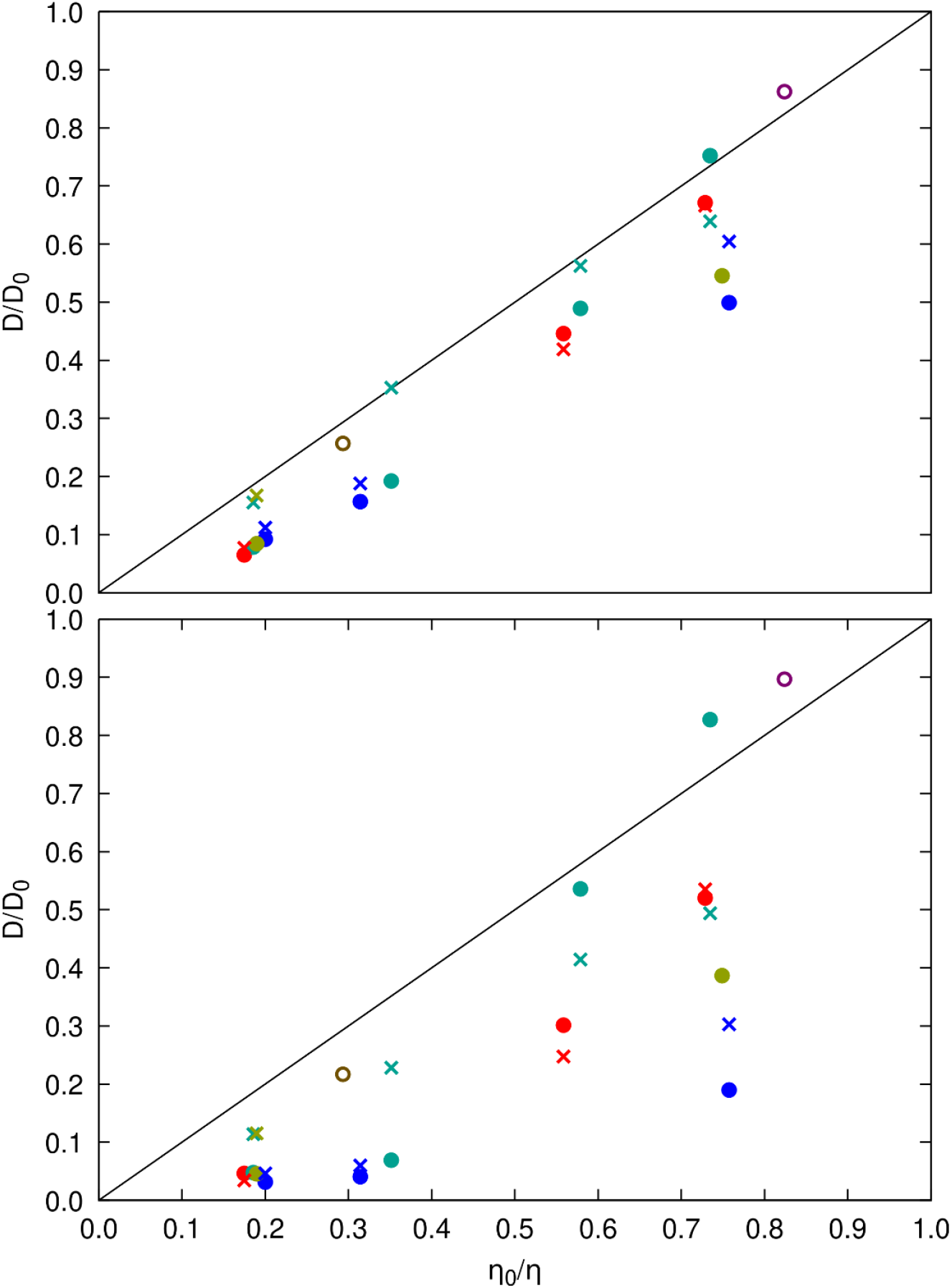
Relative diffusion vs. relative viscosity from simulations. Diffusion relative to dilute values according to data from **Table S9** (translational diffusion) and **Table S10** (rotational diffusion) is compared with the inverse of relative viscosities given in **Table S7** for the same systems. Filled circles are for SH3 diffusion in the presence of protein crowders (red: GB1, blue: lysozyme, green: BSA, tan: ovalbumin), open circles are for SH3 diffusion in urea (purple) and sucrose (brown), ‘x’ marks are for crowder self-diffusion using the same color scheme. Results based on translational diffusion are shown at the top; results for rotational diffusion are shown at the bottom.

As in previous work^14-17^, we interpret the deviation from the generalized Stokes-Einstein equation because of larger effective particle sizes (with increased *Rh*) due to clustering. While we previously described how clustering correlates with diffusional slow-down^14, 15^, the connection is clearer when separately calculated viscosity values are available. Using diffusion and viscosity values from the simulations, we can calculate apparent hydrodynamic radii according to the Stokes-Einstein equations (**Eqs. 1** and **2**). As expected, the obtained hydrodynamic radii are significantly increased in the presence of crowders (**Table S11**), both for SH3 and crowders. Where hydrodynamic radii are not increased, like with 50 g/L urea, they are within uncertainties of the dilute value. Interestingly, hydrodynamic radii estimated from translational and rotational diffusion do not always agree, as noted previously^16^. For SH3, GB1, and lysozyme, deviations are generally within statistical uncertainties, but for BSA and ovalbumin, hydrodynamic radii estimated from rotational diffusion are significantly larger than those estimated from translational diffusion. This could indicate a decoupling of translational and rotational diffusion, conceptually similar to what has been described previously in crowded BSA solutions^13^. However, unlike how rotational diffusion was reduced less than expected in previous experimental work, the larger calculated radii in our work likely result from finite-size effects, where interactions between the small number of large BSA and ovalbumin molecules across periodic boundaries hinder rotational diffusion more than translational motion.

To further quantify the effect of clustering on diffusion, we fitted the diffusion constants from simulation to the following expressions^16^:

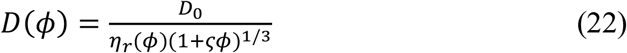

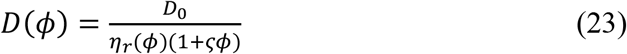

where *D*_0_ is the infinite-dilution diffusion constant, η_*r*_ is the relative viscosity according to **Eq. 20** or **Eq. 21**, and *ϕ* is the volume fraction obtained from concentration according to c[g/L] = 1430*ϕ*. The term 1 + *ςϕ* is interpreted as an effective cluster size, with *ς =* 0 indicating no clustering. We found *ς* values ranging from near zero for BSA and ovalbumin to almost 95 for SH3 diffusion in the presence of lysozyme (**Table S12**).

**Fig. 10** compares the cluster sizes (1 + *ςϕ*) with the number of interacting molecules from **Table 2** and estimated clusters sizes based on colloid theory using *ς =* 1/*τ*_*B*_ ^16^, with the values of Baxter’s stickiness parameter *τ*_*B*_ obtained from RDFs (**Table 3)**. The comparison is only shown for systems with GB1, lysozyme, and BSA crowders, as the ovalbumin system at 50 g/L has only one crowder (dimer) molecule. The RDF-based cluster size estimates were significantly lower than those obtained via diffusion fitting in all cases, which is expected because the RDF-based estimates capture direct interactions and therefore apply best to lower concentrations where larger clusters do not form.

**Figure 10.**
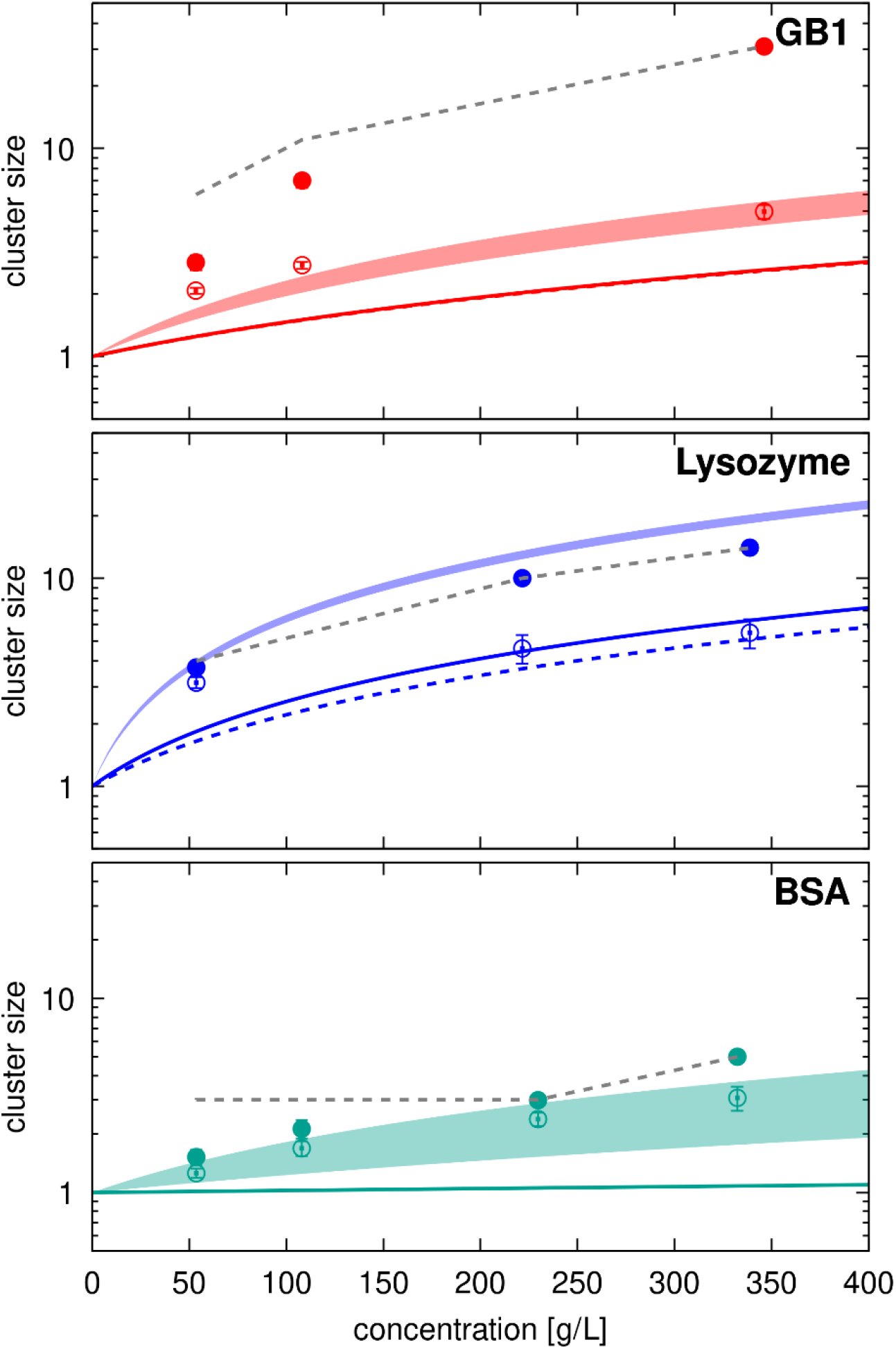
Cluster sizes estimated from diffusion *vs*. observed cluster sizes. Cluster size estimates according to 1+ζφ are compared with average SH3-crowder interactions from **Table 2**. Filled curves show range of estimates using ζ values from **Table S11** between fitting to **Eq. 22** (translational diffusion) and to **Eq. 23** (rotational diffusion) with viscosities according to **Eq. 21**. Colored solid (SH3-crowder) and dashed (crowder-crowder) lines show estimates using ζ=1/τ with τ from **Table 3**. Average direct SH3-crowder interaction counts from **Table 2** are shown as open circles with error bars, indirect interactions that include cluster interactions are shown as filled circles. Grey dashed lines indicate the maximum number of molecules that could be in a cluster given the number of molecules in the simulations.

Diffusion-based estimates align well with average cluster sizes for lysozyme and BSA, considering the limited number of crowder molecules in the simulations. For GB1, however, diffusion-based cluster size estimates are significantly lower than the actual cluster sizes, more closely reflecting the number of direct contacts (**Fig. 10** and **Table 2**). This could be due to the more dynamic nature of SH3-GB1 and GB1-GB1 interactions compared to lysozyme or BSA (**Fig. 4** and **Tables S5-S6**). Short-lived contacts, relative to the timescale of diffusion, are expected to have less impact on diffusion than long-lasting contacts. For example, in the 100 g/L SH3-GB1 system, the uncorrected translational diffusion constant for SH3 is in the presence of GB1 is 9 Å^2^/ns (**Table S9**), whereas typical long-time contact survival times are 60 ns (**Tables S5-S6**). This translates to a distance of 23 Å, which is only slightly larger than the hydrodynamic radii of SH3 or GB1. In other words, molecules would diffuse approximately their own size before losing a long-time contact, meaning clustering may only partially impact diffusion when contact life times are short. Consequently, effective cluster sizes estimated from diffusion appear smaller than actual cluster sizes.

Taken together, we relate diffusion in crowded systems to both increased viscosity and clustering. Clustering is directly related to protein interaction strength, while viscosity depends primarily on concentration, with little variation by crowder type. This is illustrated experimentally for lysozyme, which exhibits significantly weaker interactions at acidic pH and low ionic strength^81, 84, 85^, where diffusion is minimally reduced compared to basic pH, where stronger interactions lead to clustering^82, 84^. Moreover, the translational diffusion of proteins that are strongly repulsive is reduced more than rotational diffusion^13^, as few clusters form to hinder rotation, but long-range translation is limited by the presence of other proteins. Crystallin, which avoids aggregation at high concentrations in the eye lens, exemplifies this behavior^13^. BSA was also found to be slightly repulsive in that work^13^, though we found BSA to be weakly attractive in our simulations. We also found that SH3 with BSA and GB1 crowders showed slightly lower diffusion rates than in the experiments by Stadmiller *et al*.^*35*^, although other results are in excellent agreement with experimental data as described above. This may indicate a slight bias by the CHARMM c36m force field towards increased protein-protein interactions, which has been a long-standing issue with atomistic force fields^33, 93^. However, the bias here, to the extent that it is present, is clearly much smaller than with older force fields that led to strong aggregation without modifying protein-water interactions^9, 15, 33^.

### Water diffusion

Finally, we analyze the translational diffusion of water in the presence of the protein crowders, building on our previous work^100^. In that study, we concluded water diffusion slows significantly when protein crowders are present, primarily due to the fraction of water molecules in solvation shells that diffuse slower when associated with the protein surface^101^. We expected similar results for the systems studied here, but with the availability of separate viscosity data, we can add additional insight into how water diffusion correlates with the increased viscosities.

Translational diffusion coefficients for water extracted from the simulations are reported in **Table S13**. As with proteins, we estimated diffusion from mean-square displacements (**Eq. 10**), corrected the values for periodic artefacts (**Eq. 12**), and rescaled to correct for the underestimated viscosity with the TIP3P water model (**Eq. 13**). In pure water, we obtained an unscaled value of 622.5 Å^2^/ns, the same as in previous calculations that include corrections for periodic size artifacts^72^, and a rescaled value of 233.6 Å^2^/ns, very close to the experimental value of around 230 Å^2^/ns^102, 103^. Water self-diffusion in the presence of 0.15 m NaCl was slightly slower at 227.6 Å^2^/ns, with the slight decrease compared to pure water also consistent with experiment^104^. We note that the values for water self-diffusion reported here differ slightly from those in our previous work^100^, where a Langevin thermostat with a larger friction coefficient was used and the results were not corrected for the reduced viscosity with the TIP3P model.

The relative slow-down of diffusion in the crowded systems is plotted vs. viscosity in **Fig. 11**. As expected, diffusion decreases as viscosity increases. However, unlike protein diffusion, the slow-down is about half of what would be expected based solely on the viscosity of the different systems. According to the Stokes-Einstein relations (**Eqs. 1** and **2**), this would mean that water diffusion is partially decoupled form the solution viscosity. However, considering that water diffusion is primarily slowed by proximity to protein surfaces^100^, it could be argued that this reduced diffusion, essentially due to loss of bulk water because of limited space, accounts for about half of the reduced viscosity in the crowded systems, with the other half arising from protein-protein interactions and altered hydrodynamic interactions due to crowding. Because water-protein interactions vary relatively little between different proteins^100^, this may explain why we find little variation in viscosity between different systems, despite significant differences in crowder interactions.

**Figure 11.**
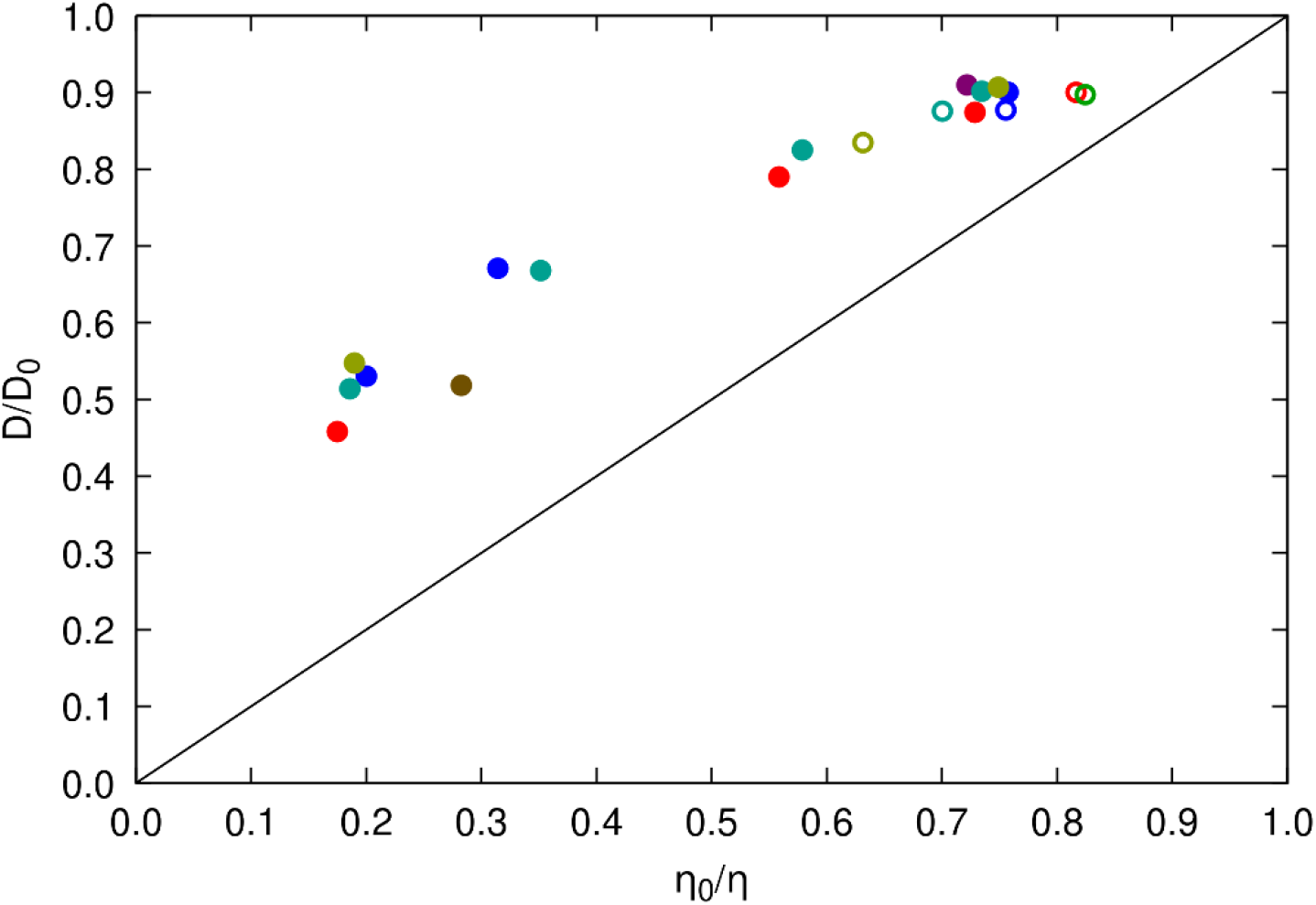
Relative translational diffusion of water vs. relative viscosity of crowded protein solutions. Diffusion and viscosity data was taken from **Tables S13** and **S7**. Reference values for determining ratios were taken from calculated diffusion and viscosity for water with 0.15 m NaCl. Filled spheres correspond to crowded systems with SH3, open spheres are for systems with a single solute. Coloring indicates the type of crowder as in **Fig. 9**.

## CONCLUSIONS

This study describes an analysis of diffusion and viscosity of crowded protein systems using extensive simulations, which showed excellent agreement with experimental data across a wide range of concentrations and extended results reported in other works^16, 17^. Our results validate the use of the CHARMM c36m force field and computationally efficient TIP3P water model (with fixed factor scaling adjustments) in capturing the delicate balance between protein-protein and protein-water interactions across dilute to highly crowded conditions.

The separate calculation of viscosity and diffusion with relatively low uncertainties allowed a careful re-evaluation of the Stokes-Einstein framework. Our central finding was that diffusion is slowed not only by increased viscosity but also by transient clustering that depends on protein interaction strength and leads to effectively larger diffusing particles. As a result, the generalized Stokes-Einstein relationship does not hold, as diffusion is reduced more than expected from an increase in viscosity alone. This extends previous findings^16^ to higher concentrations and to mixtures of proteins for which experimental data is available, in particular diffusion of SH3 in the presence of different crowders. We also highlight that the kinetics of protein contact formation matter, since contacts must persist long enough to fully affect diffusion. In addition, we find that the viscosity of protein solutions does not strongly depend on protein interactions. Considering this finding and our analysis of water diffusion, increased viscosity may largely result from reduced water mobility, as most water is bound to protein surfaces, rather than from protein-protein interactions.

Our results imply that varying degrees of transient interactions and clustering between proteins are common across a wide range of protein systems. There are examples of repulsive proteins where diffusive behavior deviates from Stokes-Einstein in the other direction, with rotational diffusion being reduced less than expected from viscosity^13^, but it appears that this may not be the typical case for many proteins. This also means that the use of largely repulsive synthetic crowders would not be an ideal mimic for protein crowding. This is not a new finding^105^, and while the perspective here is on diffusion, the same conclusion has also been found based on crowding effects on protein stability^106^.

We believe that computer simulations are now sufficiently advanced to broadly explore diffusion and viscosity for a variety of mixtures of proteins, which will bring us closer to fully understanding diffusion in crowded cellular environments.

## ACKNOWLEDGEMENTS

We thank Beatrice Caviglia and Fabio Sterpone for advice on calculating viscosities from crowded protein systems. Funding was provided by the National Science Foundation grants MCB 1817307 and by the National Institute of Health (NIGMS) grant R35 GM126948. We used computational resources at the Institute for Cyber-Enabled Research/High Performance Computing Cluster (ICER/HPCC) at Michigan State University.

