## Supplemental Tables and Figures for "Diffusion and Viscosity in Mixed Protein Solutions"

for

**Tables S1-S13**
**Figures S1-S14**
**Supplementary References**

**Table S1. Amino acid sequences of simulated systems**

| Protein | Uniprot | Residues *(numbering in full sequence)* | Sequence |
|---|---|---|---|
| SH3 | Q08012 | 1-59 | MEAIAKHDFS ATADDELSFR KGQILKILNM EDDSNWYRAE LDGKEGLIPS NYIEMKNHD |
| GB1 | P06654 | 228-282 | MQYKLILNGK TLKGETTTEA VDAATAEKVF KQYANDNGVD GEWTYDDATK TFTVTE |
| LYS | P00698 | 19-147 | KVFGRCELAA AMKRHGLDNY RGYSLGNWVC AAKFESNFNT QATNRNTDGS TDYGILQINS RWWCNDGRTP GSRNLCNIPC SALLSSDITA SVNCAKKIVS DGNGMNAWVA WRNRCKGTDV QAWIRGCRL |
| BSA | P02769 | 27-607 | HKSEIAHRFK DLGEEHFKGL VLIAFSQYLQ QCPFDEHVKL VNELTEFAKT CVADESHAGC EKSLHTLFGD ELCKVASLRE TYGDMADCCE KQEPERNECF LSHKDDSPDL PKLKPDPNTL CDEFKADEKK FWGKYLYEIA RRHPYFYAPE LLYYANKYNG VFQECCQAED KGACLLPKIE TMREKVLTSS ARQRLRCASI QKFGERALKA WSVARLSQKF PKAEFVEVTK LVTDLTKVHK ECCHGDLLEC ADDRADLAKY ICDNQDTISS KLKECCDKPL LEKSHCIAEV EKDAIPENLP PLTADFAEDK DVCKNYQEAK DAFLGSFLYE YSRRHPEYAV SVLLRLAKEY EATLEECCAK DDPHACYSTV FDKLKHLVDE PQNLIKQNCD QFEKLGEYGF QNALIVRYTR KVPQVSTPTL VEVSRSLGKV GTRCCTKPES ERMPCTEDYL SLILNRLCVL HEKTPVSEKV TKCCTESLVN RRPCFSALTP DETYVPKAFD EKLFTFHADI CTLPDTEKQI KKQTALVELL KHKPKATEEQ LKTVMENFVA FVDKCCAADD KEACFAVEGP KLVVSTQTAL A |
| OVA | P01012 | 2-386 | GSIGAASMEF CFDVFKELKV HHANENIFYC PIAIMSALAM VYLGAKDSTR TQINKVVRFD KLPGFGDSIE AQCGTSVNVH SSLRDILNQI TKPNDVYSFS LASRLYAEER YPILPEYLQC VKELYRGGLE PINFQTAADQ ARELINSWVE SQTNGIIRNV LQPSSVDSQT AMVLVNAIVF KGLWEKAFKD EDTQAMPFRV TEQESKPVQM MYQIGLFRVA SMASEKMKIL ELPFASGTMS MLVLLPDEVS GLEQLESIIN FEKLTEWTSS NVMEERKIKV YLPRMKMEEK YNLTSVLMAM GITDVFSSSA NLSGISSAES LKISQAVHAA HAEINEAGRE VVGSAEAGVD AASVSEEFRA DHPFLFCIKH IATNAVLFFG RCVSP |

Deviations from native sequence are indicated in red.

**Table S2. Molecular dynamics simulation details**

| System | Replicates | Length[a] [µs] | Average box size [Å] | Atoms | Ensemble | Temperature [K] | Pressure [atm] |
|---|---|---|---|---|---|---|---|
| sh3_gb1_50 | 3 | 2 | 98.86 | 98,235 | NPT | 298 | 1 |
| sh3_gb1_100 | 3 | 2 | 98.49 | 97,663 | NPT | 298 | 1 |
| sh3_gb1_300 | 3 | 2 | 96.35 | 93,629 | NPT | 298 | 1 |
| sh3_lys_50 | 3 | 2 | 109.90 | 134,678 | NPT | 298 | 1 |
| sh3_lys_200 | 3 | 2 | 98.84 | 99,241 | NPT | 298 | 1 |
| sh3_lys_300 | 3 | 2 | 96.99 | 94,572 | NPT | 298 | 1 |
| sh3_bsa_50 | 3 | 1 | 160.06 | 415,939 | NPT | 298 | 1 |
| sh3_bsa_100 | 3 | 2 | 126.72 | 207,398 | NPT | 298 | 1 |
| sh3_bsa_200 | 3 | 2 | 100.03 | 103,019 | NPT | 298 | 1 |
| sh3_bsa_300 | 3 | 2 | 109.79 | 137,415 | NPT | 298 | 1 |
| sh3_ova_50 | 3 | 1.2 | 138.19 | 267,604 | NPT | 298 | 1 |
| sh3_ova_300 | 3 | 2 | 108.25 | 131,428 | NPT | 298 | 1 |
| sh3_urea_50 | 3 | 2 | 96.00 | 89,777 | NPT | 298 | 1 |
| sh3_suc_300 | 3 | 2 | 92.82 | 84,254 | NPT | 298 | 1 |
| sh3 | 3 | 1 | 61.57 | 23,693 | NPT | 298 | 1 |
| gb1 | 3 | 1 | 59.52 | 21,405 | NPT | 298 | 1 |
| lys | 3 | 1 | 67.50 | 31,227 | NPT | 298 | 1 |
| bsa | 3 | 1 | 116.27 | 159,611 | NPT | 298 | 1 |
| ova | 3 | 1 | 108.01 | 128,282 | NPT | 298 | 1 |
| urea | 3 | 1 | 32.15 | 3,356 | NPT | 298 | 1 |
| suc | 3 | 1 | 36.98 | 5,113 | NPT | 298 | 1 |
| wat_salt[b] | | | 49.276[c] | 11,938 | NVT | 298 | 1 |
| wat[b] | | | 49.288[d] | 11,982 | NVT | 298 | 1 |

[a]common trajectory length in all three replicates used for analysis
[b]water systems were run for viscosity calculations and to extract translational diffusion
[c]fixed to achieve density of 1.005 g/L
[d]fixed to achieve density of 0.997 g/L

**Table S3. Experimental reference structures**

| **Protein** | **PDB** | **Method** | **Resolution** |
|---|---|---|---|
| SH3 | 2A37[1] | NMR | |
| GB1 | 2LGI[2] | NMR | |
| LYS | 3WL2 | X-ray | 0.95 Å |
| BSA | 3V03[3] | X-ray | 2.70 Å |
| OVA | 1OVA[4] | X-ray | 1.95 Å |

**Table S4. Coordinate root mean square deviations of proteins**

| System | Average[a] Cα-RMSD [Å][b] | | | | |
|---|---|---|---|---|---|
| | **SH3** | **GB1** | **Lysozyme** | **BSA** | **Ovalbumin** |
| sh3_gb1_50 | 2.15 *(0.03)* | 1.05 *(0.02)* | | | |
| sh3_gb1_100 | 2.20 *(0.14)* | 1.15 *(0.02)* | | | |
| sh3_gb1_300 | 2.10 *(0.11)* | 1.12 *(0.03)* | | | |
| sh3_lys_50 | 2.24 *(0.13)* | | 2.70 *(0.05)* | | |
| sh3_lys_200 | 2.21 *(0.16)* | | 2.72 *(0.11)* | | |
| sh3_lys_300 | 2.11 *(0.08)* | | 2.53 *(0.07)* | | |
| sh3_bsa_50 | 2.23 *(0.03)* | | | 3.02 *(0.20)* | |
| sh3_bsa_100 | 2.19 *(0.12)* | | | 2.86 *(0.23)* | |
| sh3_bsa_200 | 2.51 *(0.34)* | | | 3.11 *(0.04)* | |
| sh3_bsa_300 | 2.16 *(0.12)* | | | 3.41 *(0.14)* | |
| sh3_ova_50 | 2.65 *(0.54)* | | | | 2.94 *(0.14)* |
| sh3_ova_300 | 2.07 *(0.15)* | | | | 3.08 *(0.08)* |
| sh3_urea_50 | 2.45 *(0.09)* | | | | |
| sh3_suc_300 | 2.20 *(0.19)* | | | | |
| sh3 | 2.14 *(0.07)* | | | | |
| gb1 | | 1.04 *(0.13)* | | | |
| lys | | | 1.57 *(0.28)* | | |
| bsa | | | | 3.54 *(0.06)* | |
| ova | | | | | 3.28 *(0.09)* |

[a]omitting the first 400 ns of each trajectory; standard errors given in parentheses were estimated from variations between replicate simulations
[b]with respect to the PDB structures given in **Table S3** after optimal superposition

**Table S5. SH3-Crowder contact survival**

| System | SH3-crowder | | | | | |
|---|---|---|---|---|---|---|
| | **$\tau_1$ [ns]** | **$\tau_2$ [ns]** | **$\tau_3$ [ns]** | **$w_1$** | **$w_2$** | **$w_3$[b]** |
| sh3_gb1_50 | 69.1 | 3.5 | 0.11 | 0.77 | 0.16 | 0.07 |
| sh3_gb1_100 | 57.6 | 2.7 | 0.08 | 0.80 | 0.13 | 0.07 |
| sh3_gb1_300 | 112.6 | 1.6 | 0.02 | 0.55 | 0.19 | 0.26 |
| sh3_lys_50 | 541.0 | 3.2 | 0.08 | 0.95 | 0.03 | 0.02 |
| sh3_lys_200 | 332.9 | 2.2 | 0.05 | 0.74 | 0.08 | 0.18 |
| sh3_lys_300 | 336.7 | 1.7 | 0.04 | 0.75 | 0.06 | 0.19 |
| sh3_bsa_50 | 47.4 | 1.0 | 0.22 | 0.64 | 0.36 | 0.00 |
| sh3_bsa_100 | >10 μs[a] | 9.6 | 0.29 | 0.70 | 0.22 | 0.08 |
| sh3_bsa_200 | 599.9 | 3.5 | 0.08 | 0.85 | 0.07 | 0.08 |
| sh3_bsa_300 | 503.8 | 2.0 | 0.07 | 0.90 | 0.03 | 0.07 |
| sh3_ova_50 | 1074.4 | 0.80 | 0.06 | 0.99 | 0.01 | 0.00 |
| sh3_ova_300 | 650.4 | 1.2 | 0.10 | 0.77 | 0.05 | 0.18 |
| sh3_urea_50 | 75.5 | 0.68 | 0.08 | 0.04 | 0.52 | 0.44 |
| sh3_suc_300 | 11.5 | 2.5 | 0.15 | 0.33 | 0.44 | 0.23 |

from triple-exponential fits (**Eq. 5**) to combined survival functions from all replicates up to 20 ns
[a]exact results from the numerical fits are not reported when the fitted time scales exceed 10 μs
[b]$w_3=1.0-w_2-w_1$

**Table S6. Crowder-Crowder contact survival**

| System | Crowder-crowder | | | | | |
|---|---|---|---|---|---|---|
| | $\tau_1$ [ns] | $\tau_2$ [ns] | $\tau_3$ [ns] | $w_1$ | $w_2$ | $w_3$[b] |
| sh3_gb1_50 | 70.1 | 5.0 | 0.18 | 0.70 | 0.22 | 0.08 |
| sh3_gb1_100 | 63.7 | 3.5 | 0.10 | 0.70 | 0.20 | 0.10 |
| sh3_gb1_300 | 92.2 | 1.7 | 0.02 | 0.57 | 0.15 | 0.28 |
| sh3_lys_50 | 320.0 | 8.3 | 0.20 | 0.91 | 0.07 | 0.03 |
| sh3_lys_200 | 204.4 | 1.6 | 0.08 | 0.71 | 0.11 | 0.18 |
| sh3_lys_300 | 312.4 | 1.5 | 0.05 | 0.66 | 0.09 | 0.25 |
| sh3_bsa_50 | -[c] | | | | | |
| sh3_bsa_100 | >10 μs[a] | 2483 | 0.31 | 0.53 | 0.47 | 0.00 |
| sh3_bsa_200 | >10 μs[a] | - | - | 1.00 | 0.00 | 0.00 |
| sh3_bsa_300 | 679.9 | 1.6 | 0.05 | 0.86 | 0.03 | 0.11 |
| sh3_ova_50 | -[d] | | | | | |
| sh3_ova_300 | >10 μs[a] | - | - | 1.0 | 0.00 | 0.00 |

from triple-exponential fits (**Eq. 5**) to combined survival functions from all replicates up to 20 ns
[a]exact results from the numerical fits are not reported when the fitted time scales exceed 10 μs
[b]$w_3$=1.0-$w_2$-$w_1$
[c]contacts persisted for the entire trajectory
[d]only one dimer present in system

**Table S7. Viscosity estimates from simulation**

| System | Viscosity[a] $\eta$ [cP] | rel. viscosity $\eta/\eta_0$[c] | rel. viscosity $\eta/\eta_0$[d] |
|---|---|---|---|
| sh3_gb1_50[b] | 0.476 *(0.003)* | 1.425 | 1.373 |
| sh3_gb1_100 | 0.621 *(0.012)* | 1.859 | 1.791 |
| sh3_gb1_300[b] | 1.985*(0.002)* | 5.943 | 5.725 |
| sh3_lys_50[b] | 0.458 *(0.002)* | 1.371 | 1.321 |
| sh3_lys_200 | 1.103 *(0.025)* | 3.302 | 3.181 |
| sh3_lys_300[b] | 1.733 *(0.022)* | 5.189 | 5.000 |
| sh3_bsa_50[b] | 0.472 *(0.002)* | 1.413 | 1.361 |
| sh3_bsa_100 | 0.599 *(0.009)* | 1.793 | 1.727 |
| sh3_bsa_200 | 0.987 *(0.015)* | 2.955 | 2.847 |
| sh3_bsa_300 [b] | 1.868 *(0.024)* | 5.593 | 5.388 |
| sh3_ova_50 [b] | 0.463 *(0.004)* | 1.386 | 1.335 |
| sh3_ova_300 [b] | 1.828 *(0.016)* | 5.473 | 5.273 |
| sh3_urea_50 | 0.421 *(0.003)* | 1.260 | 1.214 |
| sh3_suc_300 | 1.182 *(0.012)* | 3.539 | 3.409 |
| sh3 | 0.424 *(0.003)* | 1.259 | 1.213 |
| gb1 | 0.425 *(0.004)* | 1.272 | 1.225 |
| lys | 0.459 *(0.003)* | 1.374 | 1.324 |
| bsa | 0.495 *(0.003)* | 1.482 | 1.428 |
| ova | 0.549 *(0.007)* | 1.644 | 1.584 |
| urea | 0.364 *(0.004)* | 1.090 | 1.050 |
| suc | 0.376 *(0.003)* | 1.126 | 1.085 |
| wat_salt | 0.347 *(0.003)* | 1.038 | 1.0 |
| wat | 0.334 *(0.006)* | 1.0 | 0.963 |

[a]averages with $\tau_{max}$=100 ps from 150 runs of 1 ns each unless noted otherwise; standard errors were obtained from dividing the total number of runs into 10 blocks consisting of 15 runs each.
[b]averages over 150 runs of 2 ns each
[c]relative to the viscosity of pure water
[d]relative to the calculated viscosity of water with 150 mM salt

**Table S8. Fitting coefficients for experimental and simulated viscosity data**

| **Protein** | **Collloid model (Eq. 20)** | | | **Mooney (Eq. 21)** | | | |
|---|---|---|---|---|---|---|---|
| | fit range | b | $\chi^2$ | fit range | S | K | $\chi^2$ |
| lysozyme | 0-250 g/L | 38.2 | 0.02 | 0-400 g/L | 2.78 | 3.01 | 0.05 |
| BSA | 0-250 g/L | 93.1 | 0.05 | 0-400 g/L | 5.67 | 1.67 | 0.09 |
| ovalbumin | 0-250 g/L | 48.2 | 0.18 | 0-400 g/L | 4.19 | 2.18 | 0.02 |
| MD proteins | 0-250 g/L | 63.4 | 0.016 | 0-250 g/L | 6.38 | 0.38 | 0.007 |

### Table S9. Translational diffusion coefficients from simulations and experiments

| Protein | System | Translational Diffusion [Å$^2$/ns] | | | | |
|---|---|---|---|---|---|---|
| | | $D_{t,PBC}$[a] | $D_{t,TIP3P}$[b] | $D_{t,MD}$[c] | $D_{t,exp}$ | $D_{t,HYDROPRO}$[5] |
| SH3 | sh3 | 14.79 *(1.92)* | 42.01 *(1.91)* | 15.77 *(0.72)* | 15[6] | 15.03 |
| | sh3_gb1_50 | 15.40 *(0.30)* | 28.19 *(0.30)* | 10.58 *(0.11)* | 13[6] | |
| | sh3_gb1_100 | 8.94 *(1.04)* | 18.74 *(1.04)* | 7.03 *(0.39)* | 8.4[6] | |
| | sh3_gb1_300 | 0.35 *(0.15)* | 3.46 *(0.15)* | 1.30 *(0.05)* | | |
| | sh3_lys_50 | 8.89 *(0.52)* | 20.96 *(0.52)* | 7.87 *(0.20)* | 7.9[6] | |
| | sh3_lys_200 | 1.02 *(0.08)* | 6.57 *(0.08)* | 2.47 *(0.03)* | | |
| | sh3_lys_300 | 0.27 *(0.01)* | 3.86 *(0.01)* | 1.45 *(0.00)* | | |
| | sh3_bsa_50 | 23.48 *(1.29)* | 31.60 *(1.29)* | 11.86 *(0.50)* | 13[6] | |
| | sh3_bsa_100 | 12.50 *(1.98)* | 20.54 *(1.98)* | 7.71 *(0.77)* | 10[6] | |
| | sh3_bsa_200 | 1.95 *(0.23)* | 8.08 *(0.23)* | 3.03 *(0.09)* | 4.3[6] | |
| | sh3_bsa_300 | 0.34 *(0.08)* | 3.30 *(0.08)* | 1.24 *(0.03)* | | |
| | sh3_ova_50 | 13.28 *(1.68)* | 22.91 *(1.68)* | 8.60 *(0.65)* | 11[6] | |
| | sh3_ova_300 | 0.47 *(0.15)* | 3.54 *(0.15)* | 1.33 *(0.06)* | | |
| | sh3_urea_50 | 21.25 *(0.97)* | 36.21 *(0.97)* | 13.59 *(0.37)* | 15[6] | |
| | sh3_suc_300 | 5.31 *(0.03)* | 10.81 *(0.03)* | 4.06 *(0.01)* | 5.6[6] | |
| GB1 | gb1 | 16.23 *(1.23)* | 44.36 *(1.25)* | 16.65 *(0.47)* | | 15.56 |
| | sh3_gb1_50 | 16.74 *(0.52)* | 29.54 *(0.52)* | 11.09 *(0.19)* | | |
| | sh3_gb1_100 | 8.79 *(0.33)* | 18.61 *(0.33)* | 6.98 *(0.13)* | | |
| | sh3_gb1_300 | 0.33 *(0.02)* | 3.43 *(0.02)* | 1.29 *(0.01)* | | |
| Lysozyme | lys | 9.93 *(0.92)* | 34.23 *(0.92)* | 12.85 *(0.35)* | 12.85[7]<br>10.3 (@20g/L)[8]<br>10.8 (@14g/L)[9]<br>10.8 (@15g/L)[10] | 12.26 |
| | sh3_lys_50 | 8.75 *(0.47)* | 20.68 *(0.47)* | 7.76 *(0.18)* | 9.1[10] | |
| | sh3_lys_200 | 0.95 *(0.03)* | 6.42 *(0.03)* | 2.41 *(0.01)* | | |
| | sh3_lys_300 | 0.28 *(0.01)* | 3.83 *(0.01)* | 1.44 *(0.00)* | | |
| BSA | bsa | 5.00 *(0.43)* | 19.19 *(0.43)* | 7.20 *(0.16)* | 6.2[11], 6.3[12] | 6.83 |
| | sh3_bsa_50 | 4.39 *(0.10)* | 12.27 *(0.10)* | 4.60 *(0.04)* | | |
| | sh3_bsa_100 | 3.13 *(0.10)* | 10.79 *(0.10)* | 4.05 *(0.04)* | | |
| | sh3_bsa_200 | 1.12 *(0.02)* | 6.78 *(0.02)* | 2.54 *(0.01)* | | |
| | sh3_bsa_300 | 0.21 *(0.00)* | 2.99 *(0.00)* | 1.12 *(0.00)* | | |
| Ovalbumin | ova | 3.09 *(0.04)* | 18.51 *(0.04)* | 6.95 *(0.02)* | 8.0[13]<br>6.0 (@20 g/L)[10] | 6.61 |
| | sh3_ova_50 | 4.63 *(0.32)* | 13.77 *(0.32)* | 5.17 *(0.12)* | 5[10] | |
| | sh3_ova_300 | 0.28 *(0.00)* | 3.10 *(0.00)* | 1.16 *(0.00)* | | |

[a]estimated directly from MSD curves according to **Eq. 10** over 2-10 ns
[b]corrected for periodic box artefacts according to **Eq. 11**
[c]corrected for underestimated viscosity with TIP3P water model according to **Eq. 13**

**Table S10. Rotational diffusion coefficients from simulations via Wong et al.[14]**

| **Protein** | **System** | $\tau_1$ [ns] | $\tau_2$ [ns] | **w** | $\tau_c$[a] [ns] | $D_{r,PBC}$[b] [1/μs] | $D_{r,TIP3P}$[c] [1/μs] |
|---|---|---|---|---|---|---|---|
| SH3 | sh3 | 1.4 | | 1.00 | 1.41 *(0.03)* | 118.0 *(3.0)* | 126.3 *(3.0)* |
| | sh3_gb1_50 | 7.5 | 1.8 | 0.38 | 2.61 *(0.16)* | 64.3 *(3.9)* | 65.7 *(3.9)* |
| | sh3_gb1_100 | 16.1 | 2.2 | 0.62 | 4.91 *(1.09)* | 36.9 *(7.3)* | 38.0 *(7.3)* |
| | sh3_gb1_300 | 634 | 1.3 | 0.96 | 33.34 *(6.29)* | 5.47 *(1.25)* | 5.83 *(1.25)* |
| | sh3_lys_50 | 15.4 | 2.4 | 0.80 | 7.69 *(1.14)* | 22.9 *(3.9)* | 24.0 *(3.9)* |
| | sh3_lys_200 | 561 | 1.4 | 0.96 | 41.75 *(10.86)* | 4.52 *(1.03)* | 5.16 *(1.03)* |
| | sh3_lys_300 | 1773 | 0.6 | 0.99 | 51.78 *(10.34)* | 3.58 *(0.89)* | 4.01 *(0.89)* |
| | sh3_bsa_50 | 17.2 | 1.3 | 0.18 | 1.62 *(0.13)* | 104.1 *(7.6)* | 104.5 *(7.6)* |
| | sh3_bsa_100 | 24.8 | 1.4 | 0.49 | 2.89 *(0.86)* | 67.2 *(16.1)* | 67.7 *(16.1)* |
| | sh3_bsa_200 | 96.7 | 1.6 | 0.94 | 21.38 *(2.53)* | 8.05 *(1.08)* | 8.74 *(1.08)* |
| | sh3_bsa_300 | 399 | 1.5 | 0.95 | 30.48 *(4.82)* | 5.74 *(0.86)* | 6.01 *(0.86)* |
| | sh3_ova_50 | 21.1 | 0.9 | 0.75 | 3.53 *(0.36)* | 48.3 *(5.4)* | 48.8 *(5.4)* |
| | sh3_ova_300 | 437 | 1.1 | 0.96 | 35.50 *(8.35)* | 5.47 *(1.66)* | 5.77 *(1.65)* |
| | sh3_urea_50 | 3.7 | 1.4 | 0.06 | 1.50 *(0.01)* | 111.4 *(1.1)* | 113.3 *(1.1)* |
| | sh3_suc_300 | 6.5 | 3.1 | 0.69 | 6.27 *(0.18)* | 26.6 *(0.8)* | 27.3 *(0.8)* |
| GB1 | gb1 | 1.2 | | 1.0 | 1.17 *(0.02)* | 142.6 *(3.0)* | 151.8 *(3.0)* |
| | sh3_gb1_50 | 6.6 | 1.4 | 0.44 | 2.10 *(0.11)* | 79.8 *(4.3)* | 81.3 *(4.3)* |
| | sh3_gb1_100 | 17.7 | 1.9 | 0.66 | 4.67 *(0.47)* | 36.4 *(3.4)* | 37.5 *(3.4)* |
| | sh3_gb1_300 | 274 | 1.6 | 0.96 | 35.6 *(3.8)* | 4.78 *(0.47)* | 5.15 *(0.47)* |
| Lysozyme | lys | 2.9 | | 1.0 | 2.87 *(0.04)* | 58.1 *(0.8)* | 64.4 *(0.8)* |
| | sh3_lys_50 | 15.3 | 3.4 | 0.82 | 9.34 *(1.18)* | 18.4 *(2.2)* | 19.5 *(2.2)* |
| | sh3_lys_200 | 461 | 1.6 | 0.97 | 53.5 *(7.3)* | 3.25 *(0.48)* | 3.89 *(0.48)* |
| | sh3_lys_300 | 1024 | 0.9 | 0.99 | 65.3 *(1.2)* | 2.55 *(0.05)* | 2.99 *(0.85)* |
| BSA | bsa | 20.6 | | 1.0 | 20.6 *(2.6)* | 8.34 *(1.00)* | 9.58 *(1.00)* |
| | sh3_bsa_50 | 47.9 | 20.0 | 0.67 | 38.6 *(3.8)* | 4.40 *(0.41)* | 4.76 *(0.41)* |
| | sh3_bsa_100 | 55.1 | 93.1 | 0.81 | 49.9 *(5.0)* | 3.42 *(0.38)* | 3.98 *(0.38)* |
| | sh3_bsa_200 | 232 | 3.2 | 0.99 | 113.5 *(11.2)* | 1.50 *(0.16)* | 2.19 *(0.16)* |
| | sh3_bsa_300 | 3329 | 1.5 | 0.99 | 246.9 *(66.8)* | 0.81 *(0.26)* | 1.09 *(0.26)* |
| Ovalbumin | ova | 22.8 | | 1.0 | 22.8 (2.5) | 7.48 *(0.82)* | 9.03 *(0.82)* |
| | sh3_ova_50 | 29.2 | 2.9 | 0.87 | 21.5 *(3.0)* | 8.10 *(1.27)* | 8.67 *(1.27)* |
| | sh3_ova_300 | 2538 | 1.0 | 1.00 | 239.8 *(36.8)* | 0.73 *(0.12)* | 1.03 *(0.12)* |

[a]not corrected for PBC
[b]estimated from fitted rotational autocorrelation functions according to **Eqs. 14-16**
[c]corrected for periodic box artefacts according to **Eq. 18**

**Table S11. Rotational diffusion coefficients from simulations via Linke *et al.*[15]**

| Protein | System | $D_{r2}/D_{r1}$ | $D_{r3}/D_{r1}$ | $D_{r,PBC}$ [1/μs] | $D_{r,TIP3P}$[a] [1/μs] |
|---|---|---:|---:|---:|---:|
| SH3 | sh3 | 0.87 *(0.02)* | 0.69 *(0.04)* | 115.8 *(1.6)* | 124.2 *(1.6)* |
| | sh3_gb1_50 | 0.84 *(0.02)* | 0.74 *(0.02)* | 57.5 *(6.1)* | 59.0 *(6.1)* |
| | sh3_gb1_100 | 0.91 *(0.05)* | 0.75 *(0.07)* | 28.1 *(5.2)* | 29.2 *(5.2)* |
| | sh3_gb1_300 | 0.83 *(0.08)* | 0.70 *(0.08)* | 4.53 *(0.85)* | 4.89 *(0.84)* |
| | sh3_lys_50 | 0.84 *(0.04)* | 0.64 *(0.04)* | 18.4 *(2.8)* | 19.5 *(2.8)* |
| | sh3_lys_200 | 0.79 *(0.07)* | 0.68 *(0.05)* | 2.90 *(0.65)* | 3.54 *(0.65)* |
| | sh3_lys_300 | 0.88 *(0.09)* | 0.75 *(0.10)* | 1.86 *(0.45)* | 2.29 *(0.45)* |
| | sh3_bsa_50 | 0.88 *(0.06)* | 0.68 *(0.01)* | 90.3 *(10.6)* | 90.7 *(10.6)* |
| | sh3_bsa_100 | 0.86 *(0.06)* | 0.77 *(0.03)* | 48.0 *(13.2)* | 48.6 *(13.2)* |
| | sh3_bsa_200 | 0.71 *(0.13)* | 0.53 *(0.10)* | 4.82 *(0.75)* | 5.51 *(0.75)* |
| | sh3_bsa_300 | 0.82 *(0.07)* | 0.66 *(0.05)* | 3.66 *(0.64)* | 3.94 *(0.64)* |
| | sh3_ova_50 | 0.85 *(0.05)* | 0.73 *(0.04)* | 25.4 *(10.0)* | 26.0 *(10.0)* |
| | sh3_ova_300 | 0.71 *(0.03)* | 0.50 *(0.06)* | 3.23 *(0.91)* | 3.52 *(0.91)* |
| | sh3_urea_50 | 0.88 *(0.02)* | 0.79 *(0.05)* | 108.3 *(5.3)* | 110.1 *(5.3)* |
| | sh3_suc_300 | 0.83 *(0.03)* | 0.74 *(0.02)* | 24.9 *(0.9)* | 25.6 *(0.9)* |
| GB1 | gb1 | 0.74 *(0.07)* | 0.62 *(0.05)* | 144.7 *(2.9)* | 154.0 *(3.0)* |
| | sh3_gb1_50 | 0.78 *(0.02)* | 0.69 *(0.02)* | 66.6 *(3.7)* | 68.1 *(3.7)* |
| | sh3_gb1_100 | 0.78 *(0.02)* | 0.67 *(0.02)* | 28.0 *(2.1)* | 29.1 *(2.1)* |
| | sh3_gb1_300 | 0.74 *(0.01)* | 0.61 *(0.01)* | 4.29 *(0.13)* | 4.65 *(0.13)* |
| Lysozyme | lys | 0.73 *(0.11)* | 0.56 *(0.06)* | 56.0 *(1.6)* | 62.3 *(1.6)* |
| | sh3_lys_50 | 0.72 *(0.03)* | 0.57 *(0.02)* | 15.8 *(1.7)* | 16.9 *(1.7)* |
| | sh3_lys_200 | 0.73 *(0.02)* | 0.60 *(0.02)* | 2.14 *(0.19)* | 2.79 *(0.19)* |
| | sh3_lys_300 | 0.77 *(0.03)* | 0.64 *(0.03)* | 1.59 *(0.07)* | 2.02 *(0.07)* |
| BSA | bsa | 0.76 *(0.04)* | 0.52 *(0.03)* | 8.17 *(0.74)* | 9.41 *(0.74)* |
| | sh3_bsa_50 | 0.59 *(0.02)* | 0.45 *(0.02)* | 3.41 *(0.04)* | 3.76 *(0.04)* |
| | sh3_bsa_100 | 0.51 *(0.03)* | 0.45 *(0.02)* | 3.01 *(0.03)* | 3.57 *(0.03)* |
| | sh3_bsa_200 | 0.52 *(0.09)* | 0.39 *(0.10)* | 1.19 *(0.06)* | 1.88 *(0.06)* |
| | sh3_bsa_300 | 0.88 *(0.02)* | 0.20 *(0.03)* | 1.26 *(0.03)* | 1.54 *(0.03)* |
| Ovalbumin | ova | 0.84 *(0.05)* | 0.63 *(0.04)* | 7.36 *(0.59)* | 8.90 *(0.59)* |
| | sh3_ova_50 | 0.70 *(0.04)* | 0.59 *(0.03)* | 6.53 *(0.13)* | 7.09 *(0.13)* |
| | sh3_ova_300 | 0.45 *(0.09)* | 0.15 *(0.02)* | 1.21 *(0.04)* | 1.51 *(0.04)* |

[a]corrected for periodic box artefacts according to **Eq. 18**

### Table S12. Rotational diffusion coefficients from simulations and experiments

| **Protein** | **System** | **$D_{r,MD}$**[a] [1/µs] Wong *et al.* | **$D_{r,MD}$**[a] [1/µs] Linke *et al.* | **$D_{r,exp}$**[b] [1/µs] | **$D_{r,HYDROPRO}$** [1/µs] |
|---|---|---:|---:|---:|---:|
| SH3 | sh3 | 47.41 *(1.11)* | 46.61 *(0.59)* | 45.0 *(1.2)*[6] | 41.3 |
| | sh3_gb1_50 | 24.66 *(1.47)* | 22.13 *(2.30)* | 37.0 *(0.8)*[6] | |
| | sh3_gb1_100 | 14.25 *(2.72)* | 10.95 *(1.96)* | 25.3 *(0.4)*[6] | |
| | sh3_gb1_300 | 2.19 *(0.47)* | 1.84 *(0.32)* | | |
| | sh3_lys_50 | 9.00 *(1.48)* | 7.33 *(1.03)* | 11.1 *(0.7)*[6] | |
| | sh3_lys_200 | 1.94 *(0.38)* | 1.33 *(0.24)* | | |
| | sh3_lys_300 | 1.50 *(0.34)* | 0.86 *(0.17)* | | |
| | sh3_bsa_50 | 39.21 *(2.87)* | 34.02 *(3.98)* | 39.7 *(0.9)*[6] | |
| | sh3_bsa_100 | 25.42 *(6.05)* | 18.23 *(4.95)* | 29.8 *(0.5)*[6] | |
| | sh3_bsa_200 | 3.28 *(0.41)* | 2.07 *(0.28)* | 15.2 *(1.3)*[6] | |
| | sh3_bsa_300 | 2.26 *(0.32)* | 1.48 *(0.24)* | | |
| | sh3_ova_50 | 18.32 *(2.02)* | 9.75 *(3.76)* | 13.9 *(1.1)*[6] | |
| | sh3_ova_300 | 2.16 *(0.62)* | 1.32 *(0.34)* | | |
| | sh3_urea_50 | 42.51 *(0.40)* | 41.33 *(2.00)* | 42.7 *(1.1)*[6] | |
| | sh3_suc_300 | 10.26 *(0.28)* | 9.60 *(0.33)* | 16.5 *(0.3)*[6] | |
| GB1 | gb1 | 56.98 *(1.13)* | 57.77 *(1.11)* | 51.8[16, 17]<br>(@ 10 g/L, pH=5.3) | 48.2 |
| | sh3_gb1_50 | 30.50 *(1.60)* | 25.54 *(1.40)* | | |
| | sh3_gb1_100 | 14.07 *(1.29)* | 10.93 *(0.80)* | | |
| | sh3_gb1_300 | 1.93 *(0.18)* | 1.75 *(0.05)* | | |
| Lysozyme | lys | 24.17 *(0.31)* | 23.37 *(0.61)* | 19.2[18]<br>(@ 80 g/L, pH=4, no salt)<br>9.5[18]<br>(@ 79 g/L, pH=4, 0.3 M NaCl)<br>7.6[18]<br>(@ 78 g/L, pH=9, no salt<br>24.2[19]<br>(@ 50 g/L, pH=3.8, no salt) | 23.1 |
| | sh3_lys_50 | 7.33 *(0.83)* | 6.34 *(0.64)* | | |
| | sh3_lys_200 | 1.46 *(0.18)* | 1.04 *(0.07)* | | |
| | sh3_lys_300 | 1.12 *(0.02)* | 0.76 *(0.03)* | | |
| BSA | bsa | 3.60 *(0.38)* | 3.53 *(0.28)* | 3.4[20]<br>(@ 3 g/L, pH=8, no salt) | 3.74 |
| | sh3_bsa_50 | 1.78 *(0.15)* | 1.41 *(0.02)* | | |
| | sh3_bsa_100 | 1.49 *(0.14)* | 1.34 *(0.01)* | | |
| | sh3_bsa_200 | 0.82 *(0.06)* | 0.70 *(0.02)* | | |
| | sh3_bsa_300 | 0.41 *(0.10)* | 0.58 *(0.01)* | | |
| Ovalbumin | ova | 3.39 *(0.31)* | 3.34 *(0.22)* | | 3.46 |
| | sh3_ova_50 | 3.25 *(0.48)* | 2.66 *(0.05)* | | |
| | sh3_ova_300 | 0.39 *(0.05)* | 0.57 *(0.01)* | | |

[a]corrected for underestimated viscosity with TIP3P water model according to **Eq. 19**
[b]obtained from experimental values of $\tau_c$ via $D_r=1/(6\tau_c)$

**Table S13. Hydrodynamic radii according to Stokes-Einstein**

| Protein | System | $R_h$ [Å] from $D_{t,TIP3P}$[a] | $R_h$ [Å] from $D_{r,TIP3P}$[b] Wong *et al.* | $R_h$ [Å] from $D_{r,TIP3P}$[b] Linke *et al.* |
|---|---|---|---|---|
| SH3 | sh3 | 15.03 *(0.65)* | 15.52 *(0.12)* | 15.60 *(0.07)* |
| | sh3_gb1_50 | 16.30 *(0.18)* | 17.40 *(0.34)* | 18.11 *(0.68)* |
| | sh3_gb1_100 | 18.87 *(1.08)* | 19.40 *(1.27)* | 21.12 *(1.25)* |
| | sh3_gb1_300 | 31.92 *(1.28)* | 24.63 *(1.59)* | 25.95 *(1.36)* |
| | sh3_lys_50 | 22.75 *(0.56)* | 24.88 *(1.25)* | 26.57 *(1.21)* |
| | sh3_lys_200 | 30.12 *(0.36)* | 31.25 *(2.27)* | 35.24 *(2.05)* |
| | sh3_lys_300 | 32.58 *(0.07)* | 29.24 *(1.94)* | 35.08 *(2.07)* |
| | sh3_bsa_50 | 14.68 *(0.58)* | 14.95 *(0.38)* | 15.75 *(0.67)* |
| | sh3_bsa_100 | 18.11 *(1.89)* | 16.43 *(1.56)* | 18.55 *(2.03)* |
| | sh3_bsa_200 | 27.40 *(0.79)* | 26.83 *(1.03)* | 31.36 *(1.41)* |
| | sh3_bsa_300 | 35.48 *(0.81)* | 24.66 *(1.23)* | 28.42 *(1.41)* |
| | sh3_ova_50 | 20.94 *(1.60)* | 19.49 *(0.68)* | 25.92 *(3.84)* |
| | sh3_ova_300 | 33.82 *(1.39)* | 25.77 *(2.17)* | 30.26 *(2.49)* |
| | sh3_urea_50 | 14.34 *(0.38)* | 15.09 *(0.05)* | 15.24 *(0.24)* |
| | sh3_suc_300 | 17.28 *(0.12)* | 17.18 *(0.16)* | 17.57 *(0.21)* |
| GB1 | gb1 | 14.20 *(0.40)* | 14.59 *(0.10)* | 14.53 *(0.09)* |
| | sh3_gb1_50 | 15.62 *(0.29)* | 16.21 *(0.29)* | 17.36 *(0.36)* |
| | sh3_gb1_100 | 19.06 *(0.34)* | 19.23 *(0.62)* | 21.65 *(0.58)* |
| | sh3_gb1_300 | 32.10 *(0.15)* | 25.31 *(0.81)* | 26.42 *(0.22)* |
| Lysozyme | lys | 18.40 *(0.50)* | 19.42 *(0.08)* | 19.65 *(0.17)* |
| | sh3_lys_50 | 23.12 *(0.47)* | 26.50 *(1.04)* | 28.04 *(0.74)* |
| | sh3_lys_200 | 30.82 *(0.13)* | 33.88 *(1.33)* | 38.47 *(0.74)* |
| | sh3_lys_300 | 32.90 *(0.06)* | 31.62 *(0.17)* | 36.31 *(0.36)* |
| BSA | bsa | 32.81 *(0.72)* | 36.84 *(1.33)* | 36.98 *(1.01)* |
| | sh3_bsa_50 | 37.71 *(0.33)* | 41.92 *(1.25)* | 45.19 *(0.17)* |
| | sh3_bsa_100 | 33.79 *(0.31)* | 41.11 *(1.24)* | 42.47 *(0.14)* |
| | sh3_bsa_200 | 32.61 *(0.09)* | 42.38 *(1.00)* | 44.59 *(0.47)* |
| | sh3_bsa_300 | 39.14 *(0.04)* | 44.21 *(3.28)* | 38.53 *(0.25)* |
| Ovalbumin | ova | 33.98 *(0.08)* | 37.52 *(1.13)* | 37.63 *(0.82)* |
| | sh3_ova_50 | 34.47 *(0.80)* | 34.80 *(1.58)* | 36.88 *(0.23)* |
| | sh3_ova_300 | 38.50 *(0.06)* | 44.60 *(1.71)* | 39.05 *(0.32)* |

[a]obtained from translational diffusion according to $R_h=k_BT/(6\pi\eta D_t)$
[b]obtained from rotational diffusion according to $R_h=(k_BT/(8\pi\eta D_r))^{1/3}$

**Table S12. Fitting coefficients for translational and rotational diffusion from simulation to cluster model**

| Diffuser | Crowder | Colloid model (Eq. 20) | | | | Mooney (Eq. 21) | | | |
|---|---|---|---|---|---|---|---|---|---|
| | | Translation | | Rotation | | Translation | | Rotation | |
| | | $\zeta$ | $\chi^2$ | $\zeta$ | $\chi^2$ | $\zeta$ | $\chi^2$ | $\zeta$ | $\chi^2$ |
| SH3 | GB1 | 27.24 | *0.04* | 16.03 | *0.49* | 18.81 | *0.16* | 13.56 | *0.03* |
| | Lysozyme | 95.07 | *0.26* | 89.56 | *1.92* | 73.29 | *0.11* | 81.20 | *1.66* |
| | BSA | 16.31 | *0.76* | 3.95 | *68.4* | 11.73 | *1.87* | 3.28 | *102.5* |
| | Ovalbumin | 71.36 | *0.02* | 31.20 | *1.09* | 52.12 | *0.00* | 27.03 | *0.84* |
| GB1 | GB1 | 31.27 | *0.14* | 17.50 | *4.94* | 22.15 | *0.39* | 15.03 | *8.28* |
| Lysozyme | Lysozyme | 42.72 | *0.15* | 46.20 | *0.74* | 31.29 | *0.06* | 41.05 | *0.59* |
| BSA | BSA | 10.00 | *1.50* | 10.54 | *0.39* | 4.97 | *0.85* | 8.32 | *0.28* |
| Ovalbumin | Ovalbumin | 9.98 | *0.07* | 0.04 | *0.23* | 3.60 | *0.01* | -0.33 | *0.42* |

fit of simulation data against **Eq. 22** for translational diffusion or **Eq. 23** for rotational diffusion with different estimates of relative viscosities using the coefficients for MD simulations from **Table S8**.

**Table S13. Translational diffusion coefficients of water from simulations**

| Protein | System | Translational Diffusion [Å²/ns] | | |
|---|---|---|---|---|
| | | $D_{t,PBC}$[a] | $D_{t,TIP3P}$[b] | $D_{t,MD}$[c] |
| SH3 | sh3 | 520.4 *(0.3)* | 544.0 *(0.3)* | 204.2 *(0.1)* |
| | sh3_gb1_50 | 516.8 *(0.1)* | 529.9 *(0.1)* | 198.9 *(0.1)* |
| | sh3_gb1_100 | 469.4 *(0.5)* | 479.4 *(0.5)* | 179.9 *(0.2)* |
| | sh3_gb1_300 | 274.6 *(1.3)* | 277.8 *(1.3)* | 104.2 *(0.5)* |
| | sh3_lys_50 | 533.7 *(0.0)* | 546.0 *(0.0)* | 204.9 *(0.0)* |
| | sh3_lys_200 | 401.2 *(1.7)* | 406.9 *(1.7)* | 152.7 *(0.6)* |
| | sh3_lys_300 | 317.7 *(2.5)* | 321.4 *(2.5)* | 120.6 *(0.9)* |
| | sh3_bsa_50 | 538.8 *(0.0)* | 547.0 *(0.0)* | 205.3 *(0.2)* |
| | sh3_bsa_100 | 492.2 *(0.7)* | 500.4 *(0.7)* | 187.8 *(0.3)* |
| | sh3_bsa_200 | 398.7 *(2.0)* | 405.0 *(2.0)* | 152.0 *(0.8)* |
| | sh3_bsa_300 | 308.4 *(1.5)* | 311.4 *(1.5)* | 116.9 *(0.6)* |
| | sh3_ova_50 | 540.6 *(0.3)* | 550.3 *(0.3)* | 206.5 *(0.1)* |
| | sh3_ova_300 | 328.4 *(1.9)* | 331.5 *(1.9)* | 124.4 *(0.7)* |
| | sh3_urea_50 | 536.5 *(0.1)* | 551.9 *(0.1)* | 207.1 *(0.0)* |
| | sh3_suc_300 | 308.6 *(0.1)* | 314.2 *(0.1)* | 117.9 *(0.0)* |
| GB1 | gb1 | 521.5 *(0.1)* | 545.8 *(0.2)* | 204.8 *(0.1)* |
| Lysozyme | lys | 512.0 *(0.5)* | 531.9 *(0.5)* | 199.6 *(0.2)* |
| BSA | bsa | 520.7 *(0.4)* | 531.4 *(0.4)* | 199.4 *(0.2)* |
| Ovalbumin | ova | 495.8 *(0.3)* | 506.2 *(0.3)* | 190.0 *(0.1)* |
| | wat_salt[d] | 568.8 (*0.3*) | 606.4 *(0.3)* | 227.6 *(0.1)* |
| | wat[d] | 584.9 *(0.4)* | 622.5 *(0.4)* | 233.6 *(0.1)* |

[a]estimated directly from MSD curves according to **Eq. 10** over 2-10 ns
[b]corrected for periodic box artefacts according to **Eq. 11**
[c]corrected for underestimated viscosity with TIP3P water model according to **Eq. 13**
[d]diffusion was estimated by averaging over 150 simulations of 1 ns each

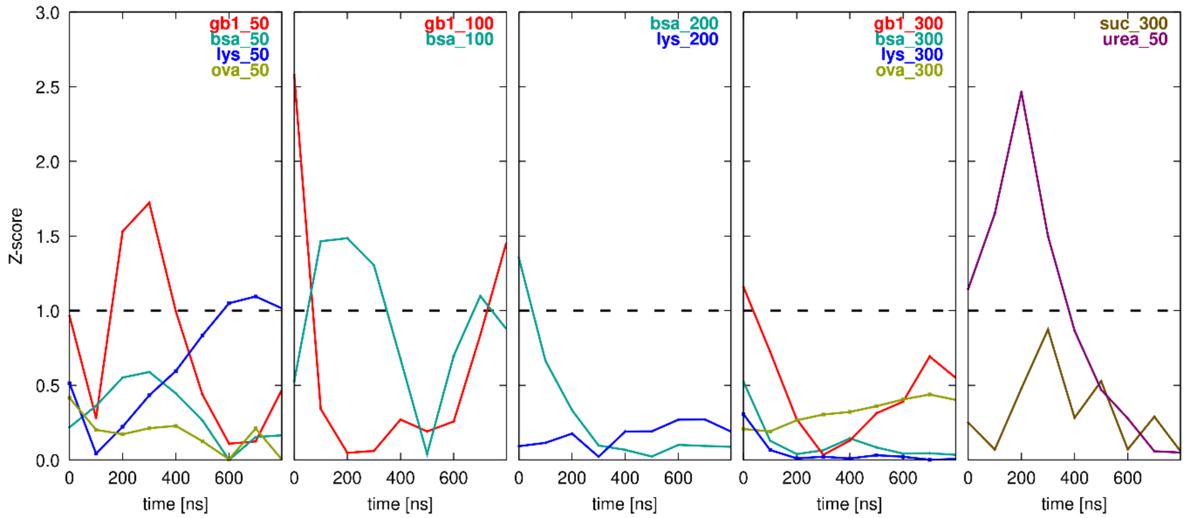


**Figure S1. Simulation convergence.** Based on average number of interactions with crowders a Z-score is shown as a function of simulation time *t* comparing the average over [*t*,*t*+400ns] ($a_{400}$) *vs.* the average over [*t*, $t_{max}$] ($a_{max}$), where $t_{max}$ is the total simulation time used for analysis according to **Table S2**. The Z-score was calculated as $|a_{400} - a_{max}|/(SEM(a_{400}) + SEM(a_{max}))$ using the standard error of the mean (SEM) for the two averages. The dashed lines indicate a Z-score of 1.

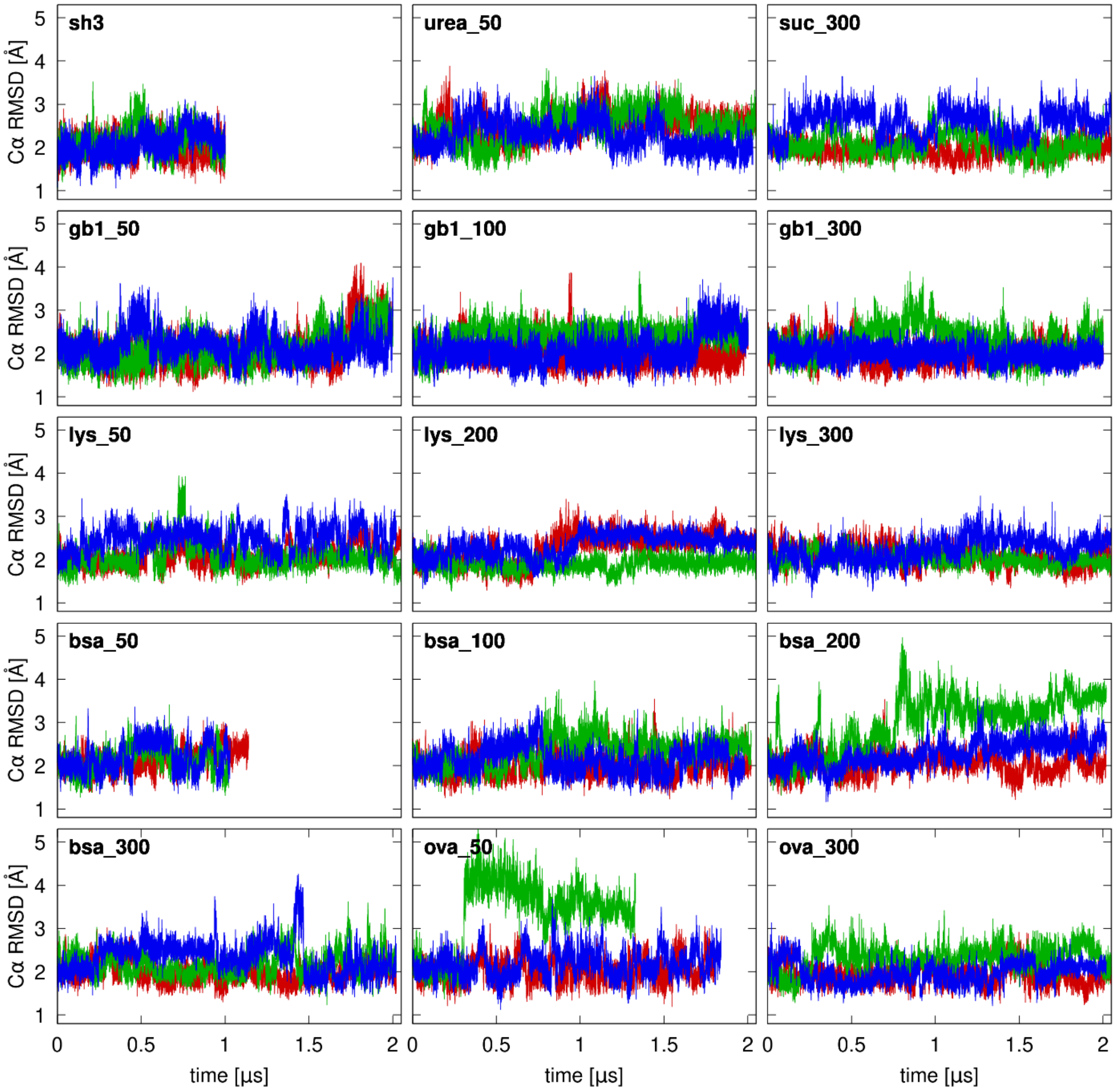


**Figure S2. Coordinate root mean-square deviation time series for SH3.** RMSD values were calculated for Cα atoms with respect to 2A37 after optimal superposition. Colors distinguish different simulation replicates.

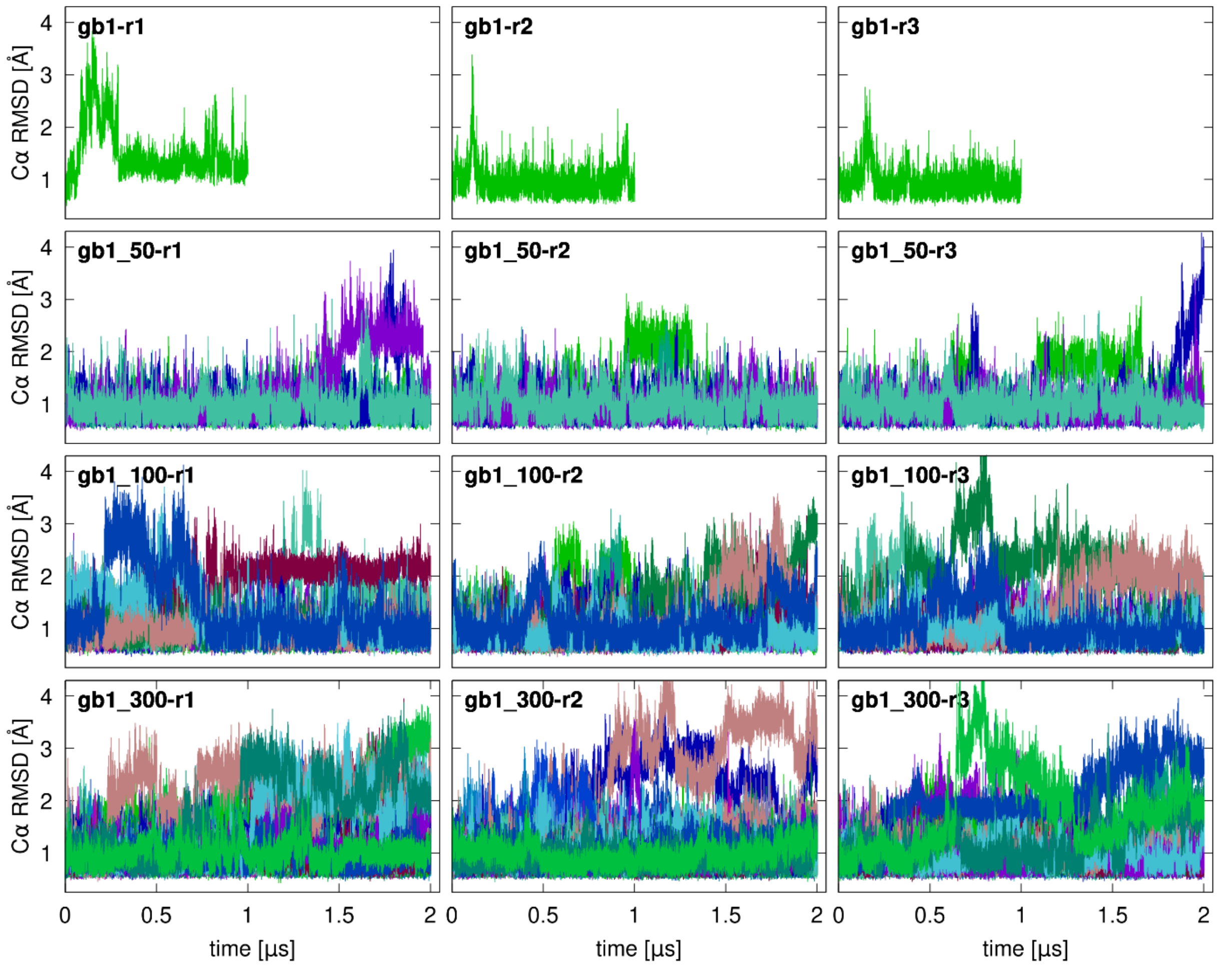


**Figure S3. Coordinate root mean-square deviation time series for GB1.** Results in columns 1-3 correspond to simulation replicates for each system. RMSD values were calculated for Cα atoms with respect to 2LGI after optimal superposition. Colors distinguish different protein copies.

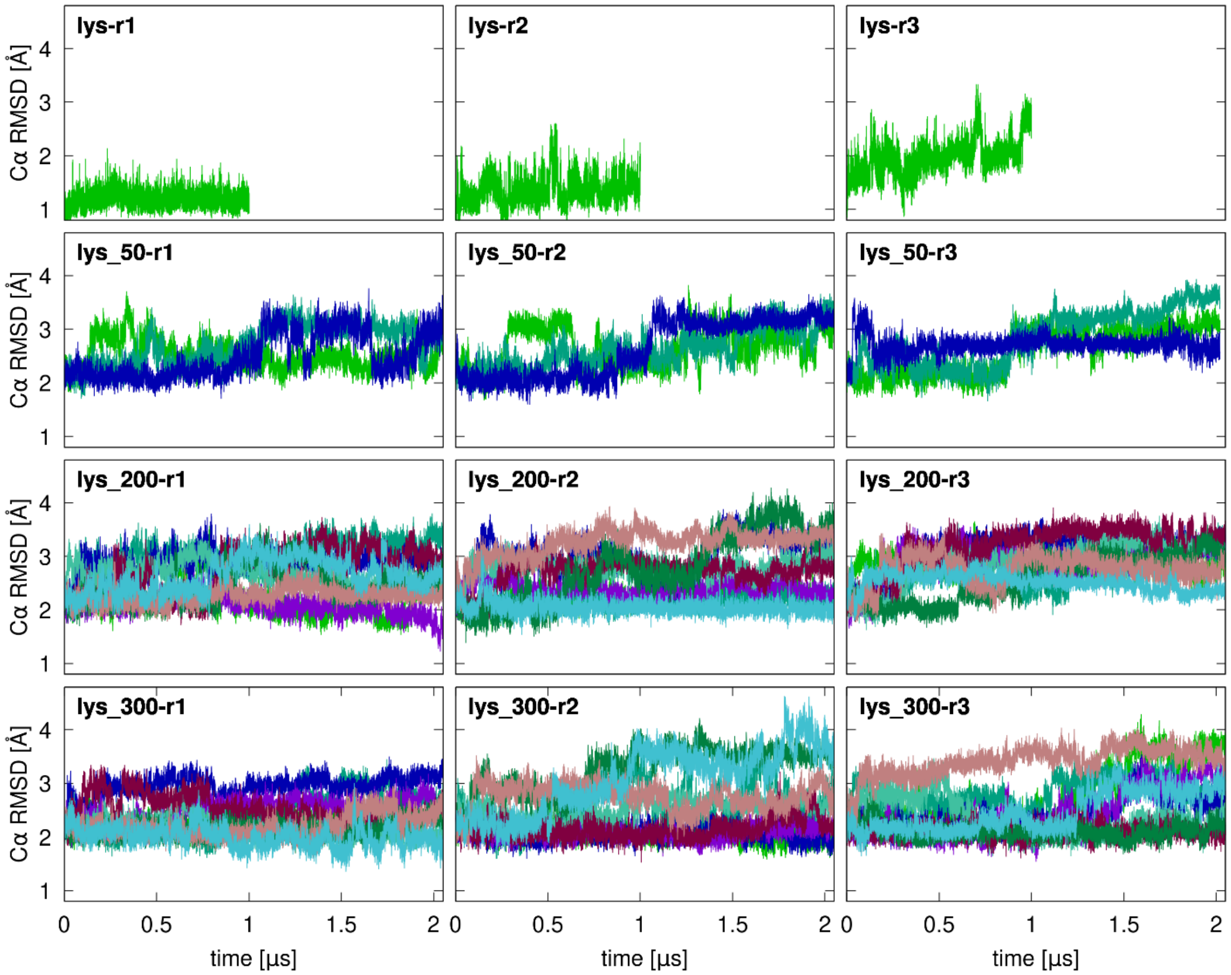


**Figure S4. Coordinate root mean-square deviation time series for lysozyme.** Results in columns 1-3 correspond to simulation replicates for each system. RMSD values were calculated for Cα atoms with respect to 2WL2 after optimal superposition. Colors distinguish different protein copies.

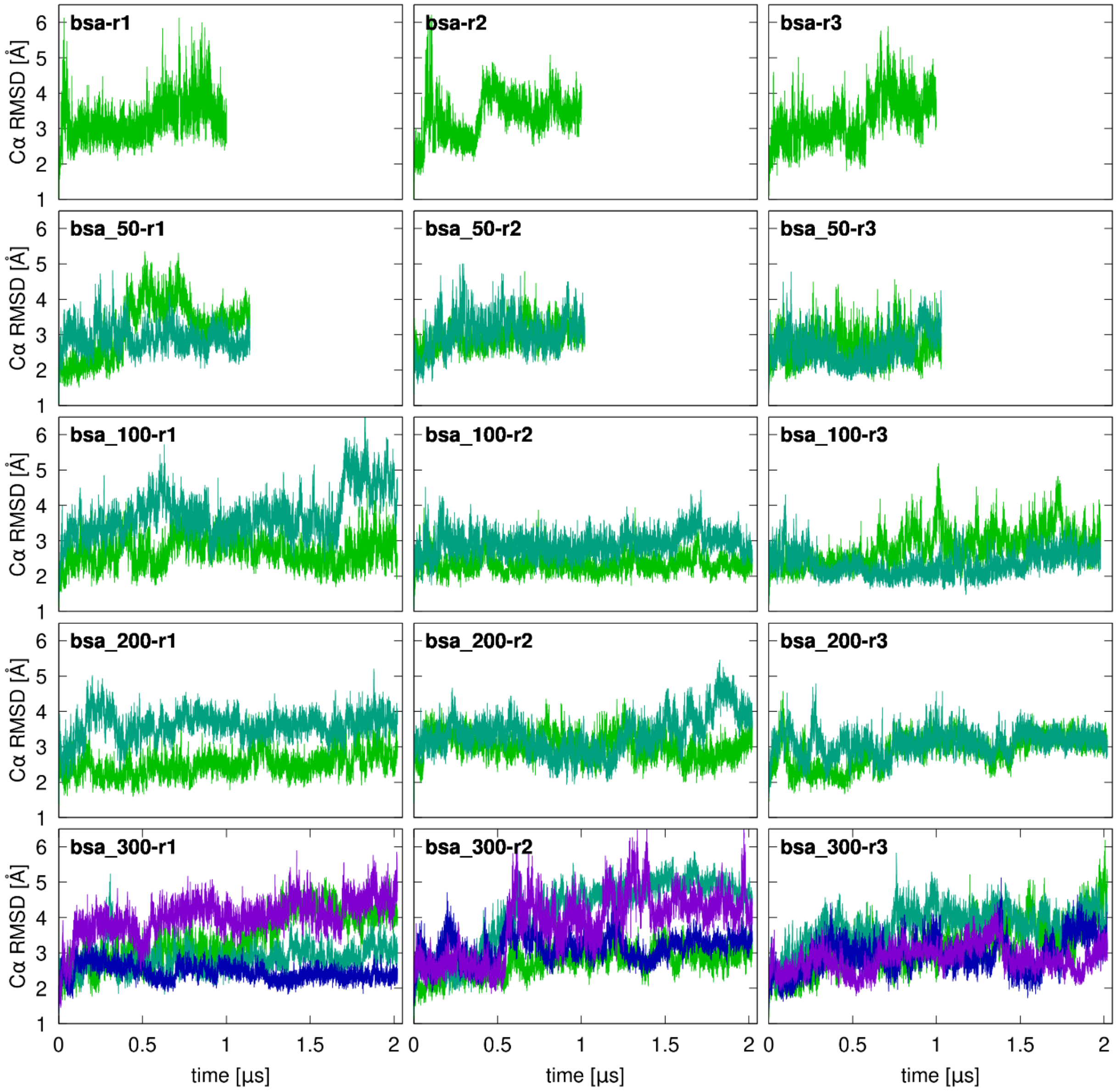


**Figure S5. Coordinate root mean-square deviation time series for BSA.** Results in columns 1-3 correspond to simulation replicates for each system. RMSD values were calculated for Cα atoms with respect to 3V03 after optimal superposition. Colors distinguish different protein copies.

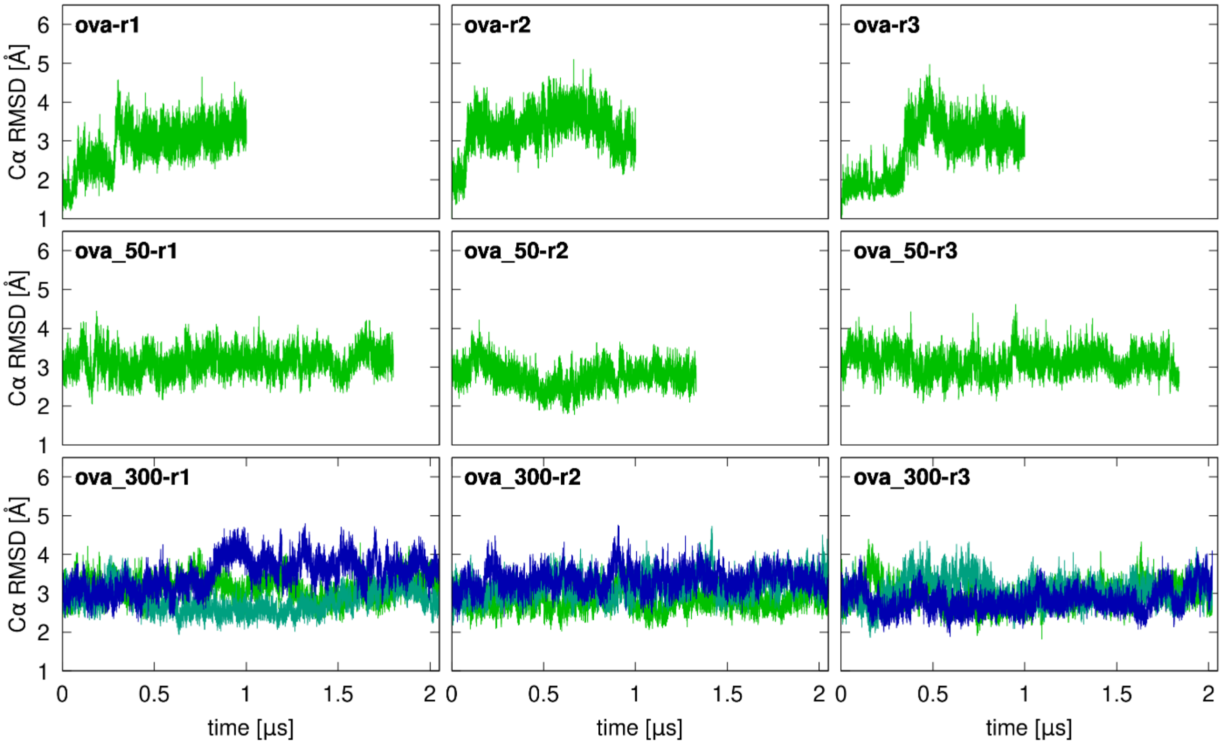


**Figure S6. Coordinate root mean square deviation time series for ovalbumin.** Results in columns 1-3 correspond to simulation replicates for each system. RMSD values were calculated for Cα atoms with respect to 1OVA after optimal superposition. Colors distinguish different protein copies. Note that ovalbumin was modeled as a homo-dimer.

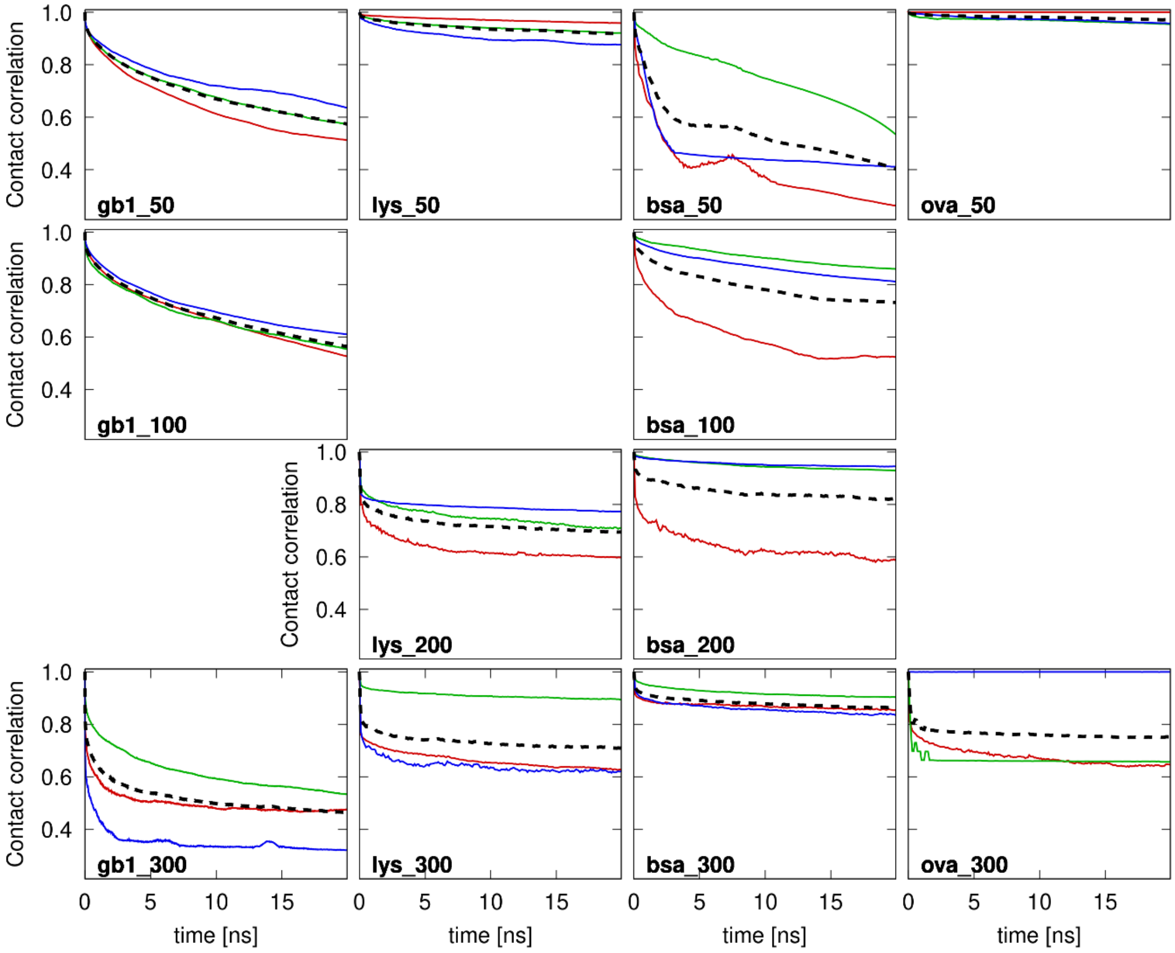


**Figure S7. SH3-protein crowder contact survival functions.** Contact survival functions between SH3 and protein crowder molecules were calculated according to **Eq. 2**. Survival functions averaged over crowder proteins for different replicates are shown in red, blue, and green. Averaged contact functions used for fitting are shown as black dashed lines. Fits are reported in **Table S5**.

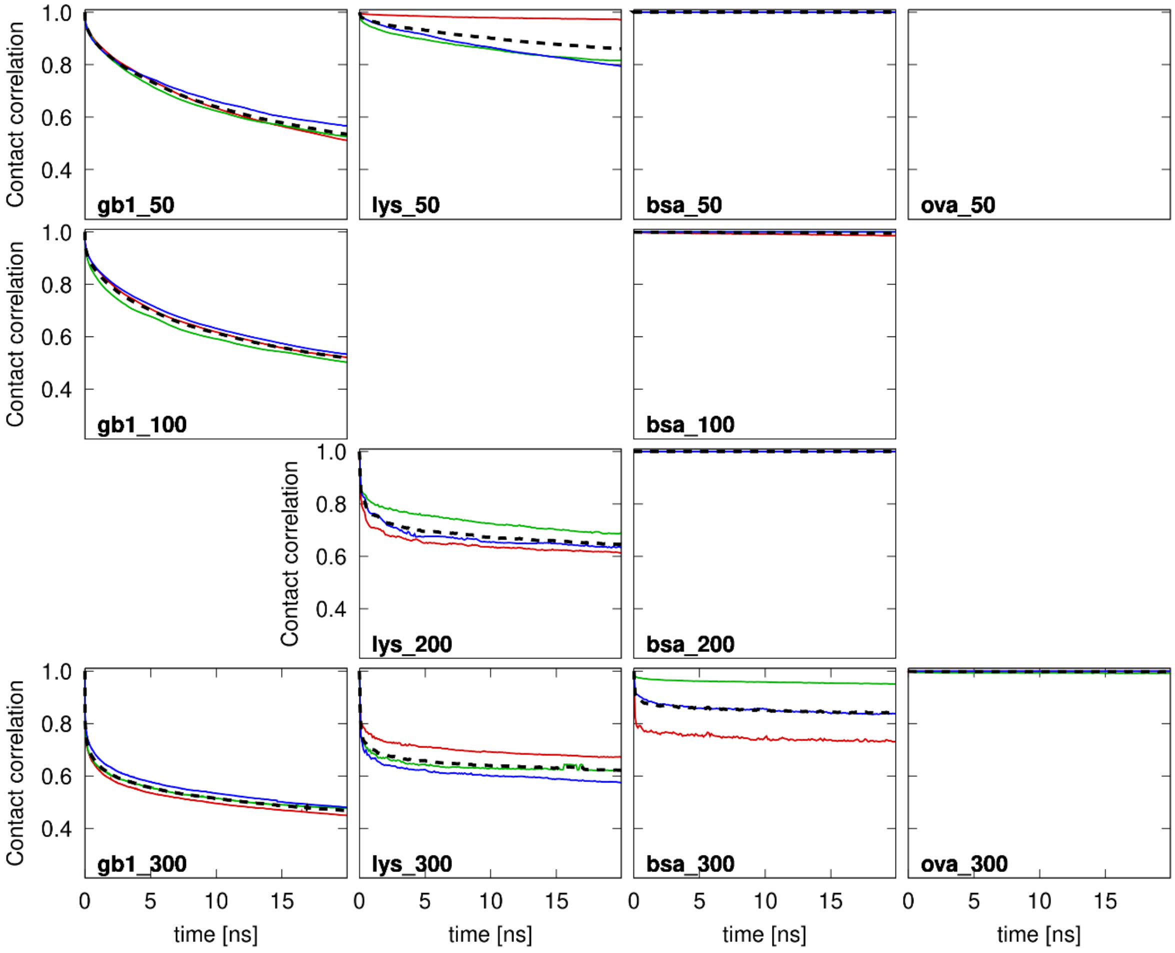


**Figure S8. Crowder contact survival functions for systems with SH3.** Contact survival functions were calculated according to **Eq. 2**. Survival functions averaged over crowders are shown in red, blue, and green for different replicates. Contact functions averaged over replicates that were used for fitting are shown as black dashed lines. Fits are reported in **Table S6**.

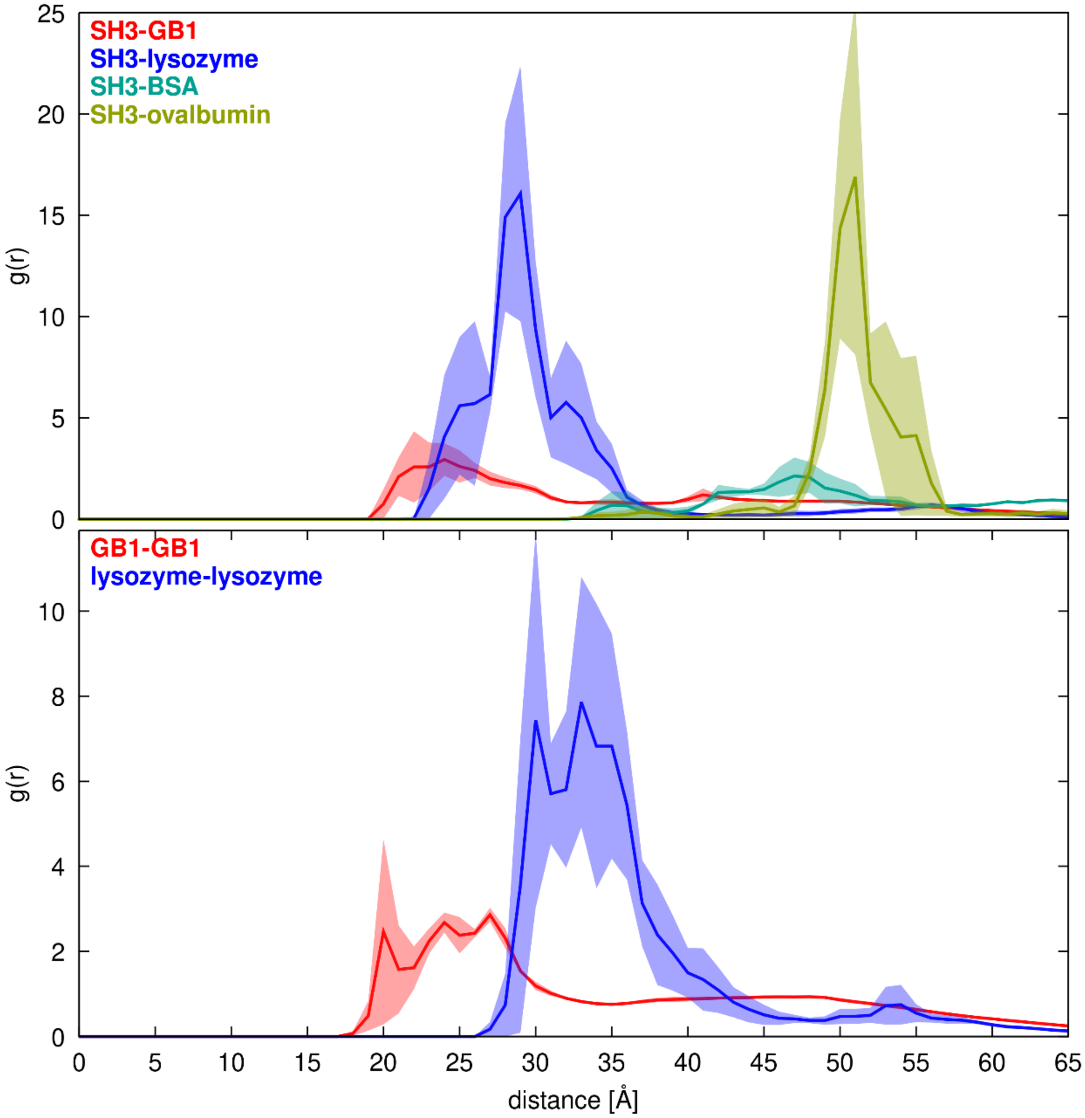


**Figure S9. Protein-protein radial distribution functions.** RDFs were calculated from protein centers of mass between SH3 and crowders (in *sh3_gb1_50*, *sh3_lys_50*, *sh3_bsa_50*, sh3_ova_50) and between GB1 or lysozyme crowders (in *sh3_gb1_50*, *sh3_lys_50*).

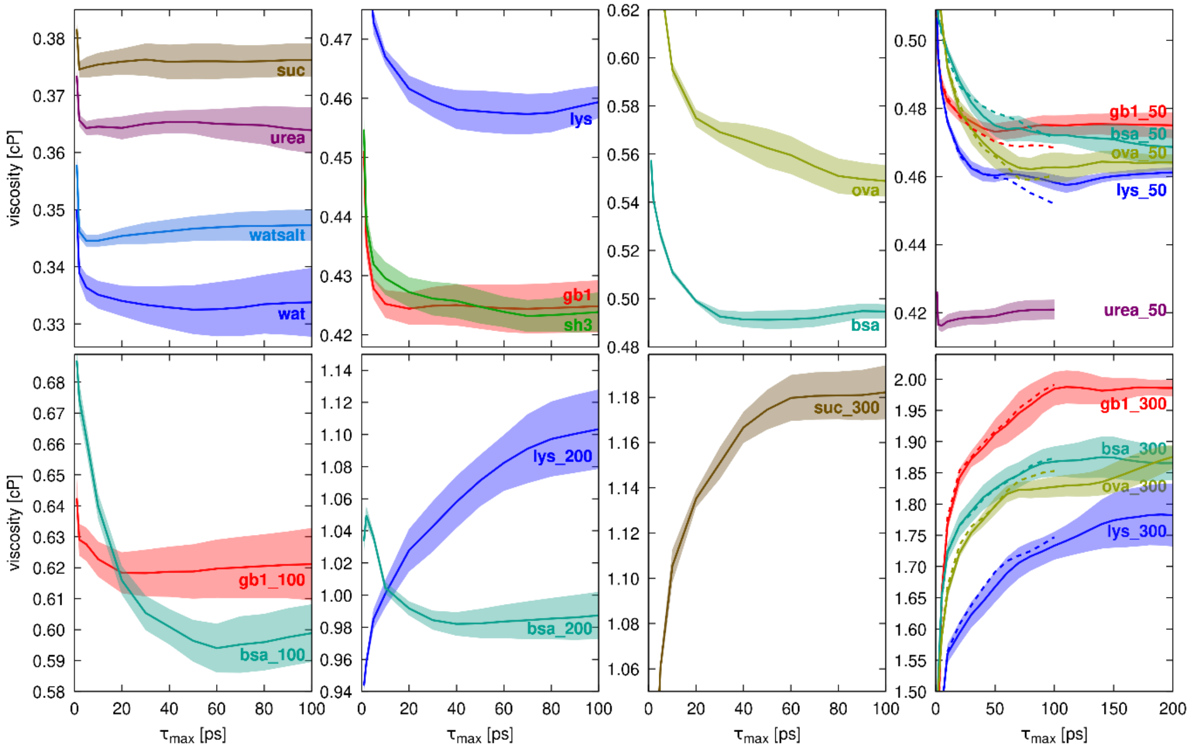


**Figure S10. Convergence of viscosity estimates from simulations.** Viscosities were estimated according to **Eq. 3** following the protocol by Zhang *et al.*[21] as a function of $\tau_{max}$. For each estimate, pressure tensor fluctuation from 150 simulations over 1 or 2 ns were used. The shaded areas indicate the standard error from variations in viscosity estimates obtained from individual trajectories. For systems that were simulated over 2 ns, the dashed line indicates viscosity estimates obtained using only the first 1 ns.

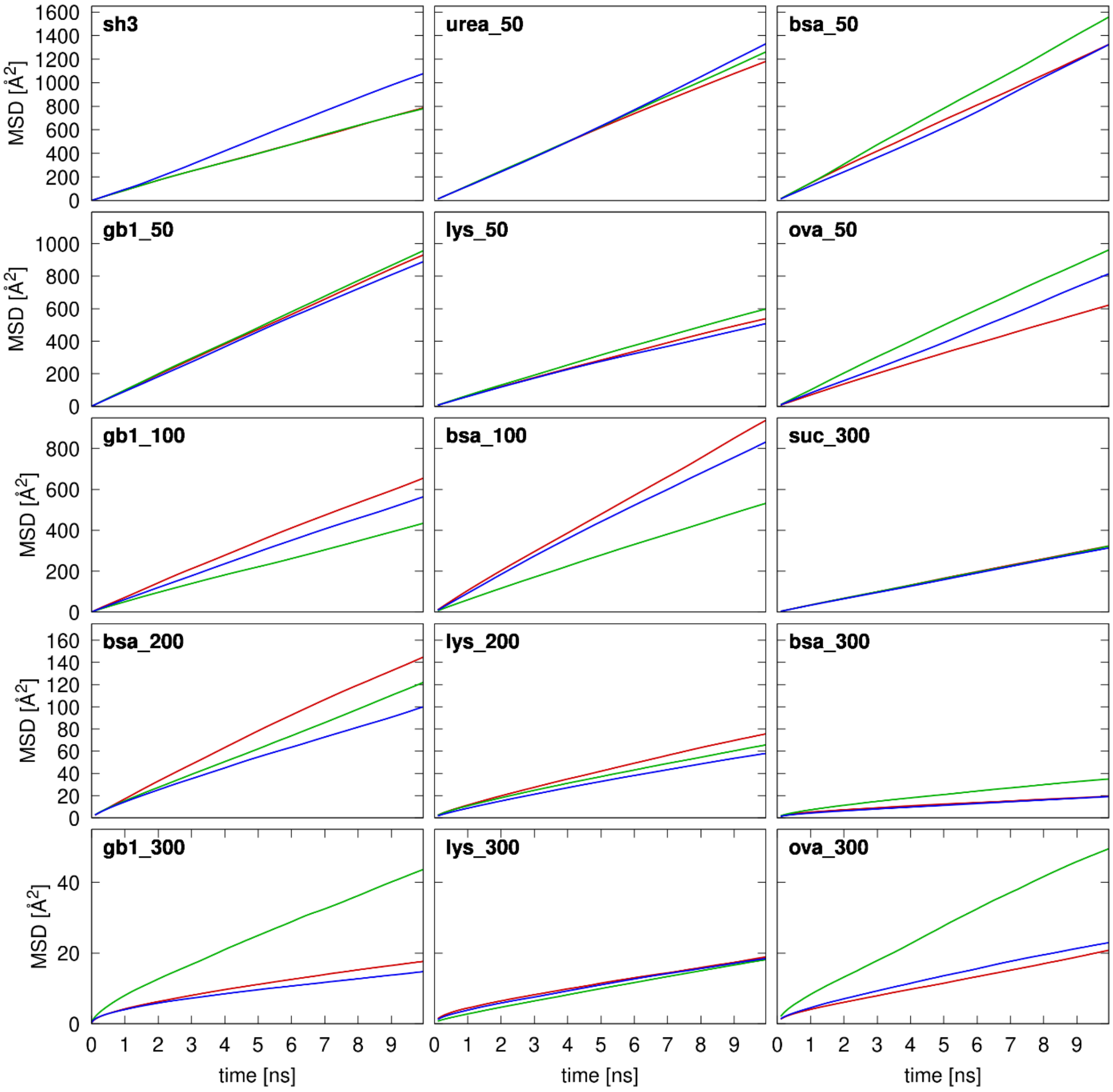


**Figure S11. Mean-square displacement vs. time for SH3.** Colors distinguish results from three replicates.

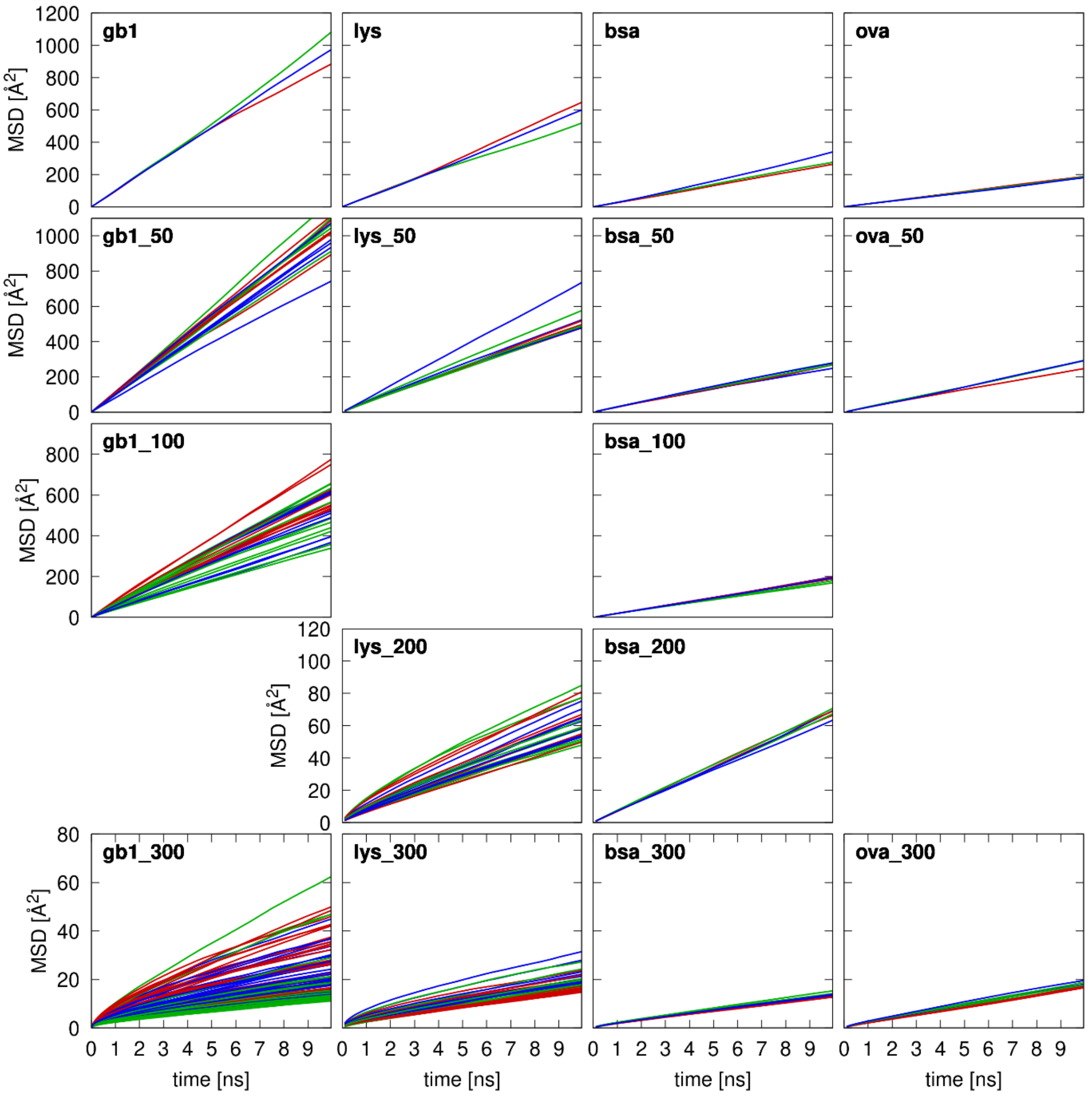


**Figure S12. Mean-square displacement vs. time for crowder proteins in systems with SH3.** Individual curves are for different protein copies. Colors distinguish results from three replicates.

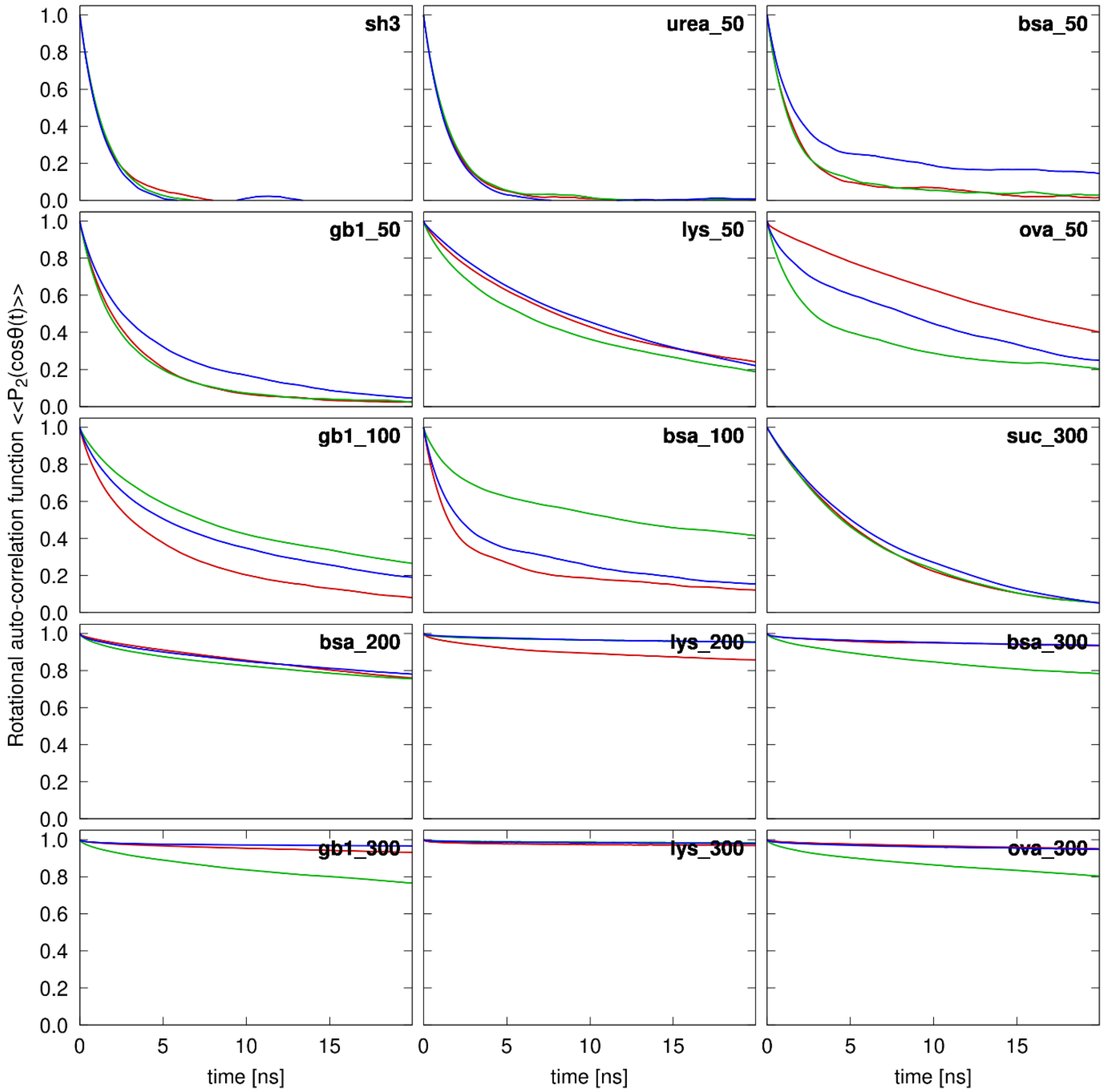


**Figure S13. Rotational correlation functions for SH3.** Colors distinguish results from three replicates.

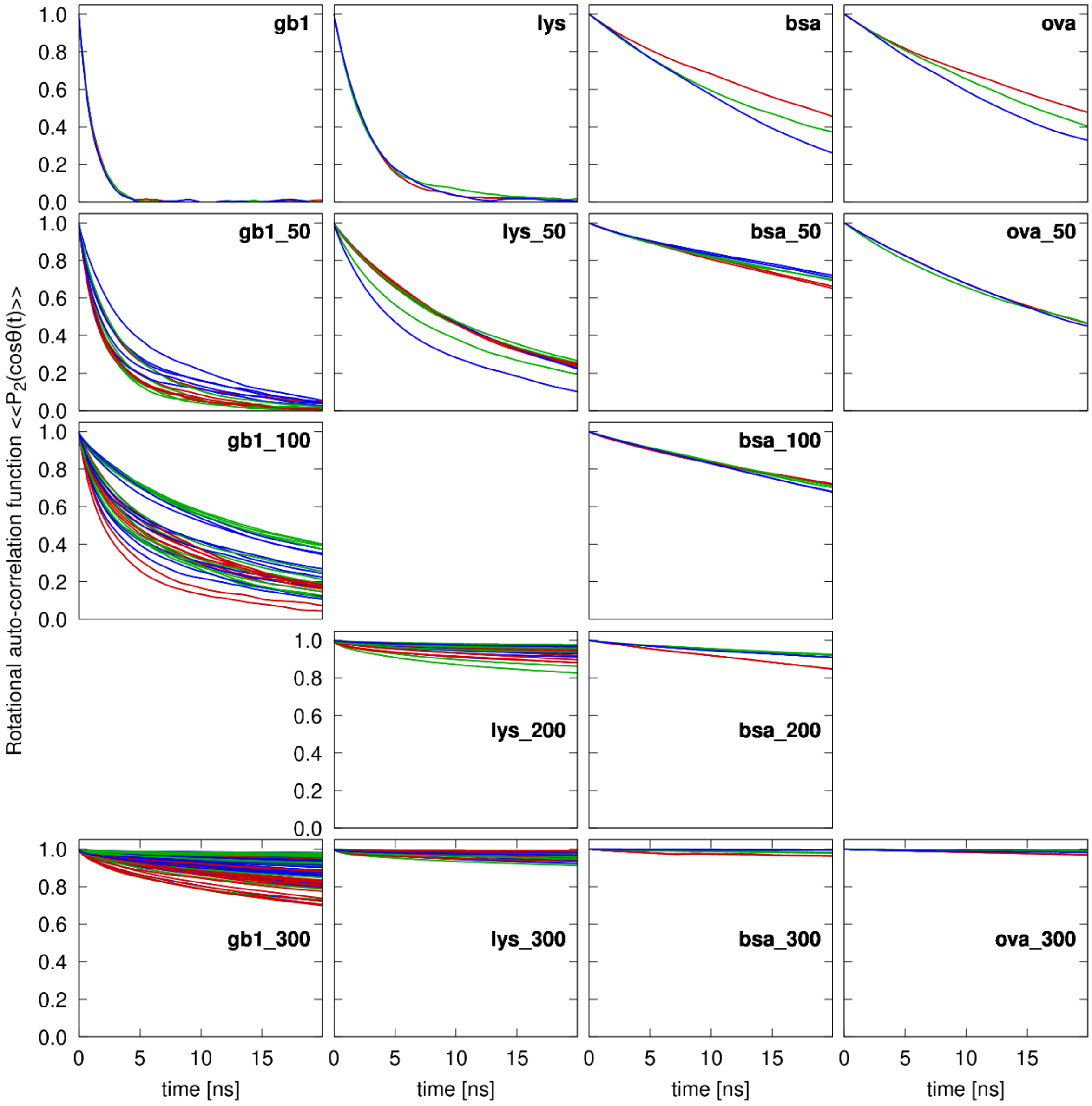


**Figure S14. Rotational correlation functions for crowder proteins in systems with SH3.** Individual curves are for different protein copies. Colors distinguish results from three replicates.
